# CD4-binding site immunogens elicit heterologous anti-HIV-1 neutralizing antibodies in transgenic and wildtype animals

**DOI:** 10.1101/2022.09.08.507086

**Authors:** Harry B. Gristick, Harald Hartweger, Maximilian Loewe, Jelle van Schooten, Victor Ramos, Thiago Y. Oliviera, Yoshiaki Nishimura, Nicholas S. Koranda, Abigail Wall, Kai-Hui Yao, Daniel Poston, Anna Gazumyan, Marie Wiatr, Marcel Horning, Jennifer R. Keeffe, Magnus A.G. Hoffmann, Zhi Yang, Morgan E. Abernathy, Kim-Marie A. Dam, Han Gao, Priyanthi N.P. Gnanapragasam, Leesa M. Kakutani, Ana Jimena Pavlovitch-Bedzyk, Michael S. Seaman, Mark Howarth, Andrew T. McGuire, Leonidas Stamatatos, Malcolm A. Martin, Anthony P. West, Michel C. Nussenzweig, Pamela J. Bjorkman

## Abstract

Passive transfer of broadly neutralizing anti-HIV-1 antibodies (bNAbs) protects against infection, and therefore eliciting bNAbs by vaccination is a major goal of HIV-1 vaccine efforts. bNAbs that target the CD4-binding site (CD4bs) on HIV-1 Env are among the most broadly active, but to date, responses elicited against this epitope in vaccinated animals have lacked potency and breadth. We hypothesized that CD4bs bNAbs resembling the antibody IOMA might be easier to elicit than other CD4bs antibodies that exhibit higher somatic mutation rates, a difficult-to-achieve mechanism to accommodate Env’s N276_gp120_ N-glycan, and rare 5-residue light chain complementarity determining region 3s (CDRL3s). As an initial test of this idea, we developed IOMA germline-targeting Env immunogens and evaluated a sequential immunization regimen in transgenic mice expressing germline-reverted IOMA. These mice developed CD4bs epitope-specific responses with heterologous neutralization, and cloned antibodies overcame neutralization roadblocks including accommodating the N276_gp120_ glycan, with some neutralizing selected HIV-1 strains more potently than IOMA. The immunization regimen also elicited CD4bs-specific responses in animals containing polyclonal antibody repertoires. Thus, germline-targeting of IOMA-class antibody precursors represents a potential vaccine strategy to induce CD4bs bNAbs.

## Introduction

A successful vaccine against HIV-1 would be the most effective way to contain the AIDS pandemic, which so far is responsible for > 36 million deaths in total and 1 - 2 million new infections each year (https://www.unaids.org/en/resources/fact-sheet). Clinical trials of vaccine candidates have revealed disappointing outcomes, and as a result, there is no currently available protective vaccine against HIV-1 (*1*), in part due to the large number of circulating HIV-1 strains (*2*). For the last decade, a major focus of HIV-1 vaccine design has been on eliciting broadly neutralizing antibodies (bNAbs), which neutralize a majority of HIV-1 strains *in vitro* at low concentrations (*1*). Multiple studies have demonstrated that passively administered bNAbs can prevent HIV-1 or simian/human immunodeficiency virus (SHIV) infection (*3–15*), suggesting a vaccination regimen that elicits bNAbs at neutralizing concentrations would be protective.

The HIV-1 Envelope protein (Env), a trimeric membrane glycoprotein comprising gp120 and gp41 subunits that is found on the surface of the virus, is the sole antigenic target of neutralizing antibodies (*16*). An impediment to HIV-1 vaccine design is that most inferred germline (iGL) precursors of known bNAbs do not bind with detectable affinity to native Envs on circulating HIV-1 strains (*17–28*). As a result, potential Env immunogens must be modified to bind and select for bNAb precursors *in vivo* during immunization (i.e., a “germline-targeting” approach). This approach has been used to activate precursors of the VRC01-class of bNAbs that target the CD4 binding site (CD4bs) on gp120 (*25, 29*). Eliciting VRC01-class bNAbs that target the CD4bs would be desirable due to their breadth and potency (*30*). However, the VRC01-class of bNAbs may be difficult to elicit due to their requirement for rare short light chain complementarity region 3 (CDRL3) loops of 5 residues (present in only ∼1% of human antibodies) (*31*) and many somatic hypermutations (SHMs), including a difficult-to achieve sequence of mutations to sterically accommodate the highly-conserved N276_gp120_ glycan (*32*).

Crystal structures of a natively glycosylated HIV-1 soluble Env trimer derived from the clade A BG505 strain (BG505 SOSIP.664) (*33*) complexed with the antibody IOMA, revealed that this CD4bs bNAb exhibits distinct properties from VRC01-class bNAbs (*34*). In common with VRC01-class bNAbs, IOMA is derived from the VH1-2 immunoglobulin heavy chain (HC) gene segment, and it binds Env with a similar overall pose as other VH1-2–derived CD4bs bNAbs, but it is not as potent or broad as many of the VRC01-class antibodies (*34*). However, unlike VRC01-class bNAbs, IOMA includes a normal-length (8 residues) CDRL3 (*34*) and is less mutated with 9.5% HC and 7% light chain (LC) nucleotide mutations to its iGL compared to VRC01 with 30% HC and 19% LC nucleotide mutations (*35, 36*). In addition, IOMA accommodates the N276_gp120_ glycan, a roadblock for raising VRC01-class bNAbs (*32*), using a relatively easy-to-achieve mechanism involving a short helical CDRL1, and four amino acid changes (including a single mutated glycine) that each require single nucleotide substitutions. By contrast, the CDRL1s of VRC01-class bNAbs include either a 3 - 6 residue deletion or large numbers of SHMs that introduce multiple glycines and/or other insertions to create flexible CDRL1 loops (*32*). Thus, IOMA-like antibodies likely represent an easier pathway for vaccine induced maturation of CD4bs precursors to mature CD4bs bNAbs.

Here we report immunogens engineered to elicit IOMA and other CD4bs bNAbs. Using these immunogens, we devised a sequential immunization strategy that elicited broad heterologous serum neutralization in both IOMA iGL knock-in and wildtype (wt) mouse models. Notably, this was achieved using fewer than half of the immunizations in other studies (*37, 38*). Moreover, IOMA-like bNAbs elicited in knock-in mice were more potent than IOMA against some strains. Finally, the immunization regimen developed in knock-in mice also elicited CD4bs-specific responses in multiple wt animals including mice, rabbits, and rhesus macaques providing a rationale for using the IOMA-targeting immunogens described here as part of an effective HIV vaccine.

## Results

### Design of IOMA-targeting immunogens

To create the IOMA iGL antibody, we reverted the HC and LC sequences of mature IOMA (*34*) to their presumptive germline sequences. The IOMA iGL HC sequence was based on human *IGHV1-2*02, IGHD3-22*01* and *IGHJ6*02* and contained 22 amino acid changes compared to the HC of IOMA, all within the V gene. The CDRH3 was unaltered due to uncertainty with respect to D gene alignment and potential P and N nucleotides – the IOMA iGL HC sequence maintains one gp120-contacting residue (W100F; Kabat numbering (*39*)) found in mature IOMA (*34*). The sequence of the IOMA iGL LC was derived from human *IGLV2-23*02* and *IGLJ2*01* containing 16 amino acid changes compared with mature IOMA, including 3 SHMs in CDRL3. Of the 3 mutations in CDRL3, two non-contact amino acids (V96, A97; Kabat numbering (*39*)) at the V-J junction were left as in mature IOMA (Figure S1A, Table S1).

No currently available germline-targeting CD4bs immunogens bind IOMA iGL with detectable affinity (Figure S1B), and IOMA iGL does not neutralize primary HIV-1 strains (Figure S1C). We therefore used *in vitro* selection methods to identify potential IOMA-targeting immunogens (Figure 1A). We chose *S. cerevisiae* yeast display for selecting an IOMA-targeting immunogen for two reasons: (i) Yeast libraries can contain up to 1 x 10^9^ variants (*40, 41*) and therefore allow screening a large number of immunogen constructs, a necessity since we were starting with no detectable binding of IOMA iGL to any CD4bs-targeting immunogens, and (ii) *S. cerevisiae* attach different forms of N-linked glycans to glycoproteins than mammalian cells; e.g., yeast can add up to 50 mannoses to Man_8-9_GlcNAc_2_ (*42*). Such glycan differences may be an advantage because N-glycosylated immunogens selected in a yeast library to bind IOMA iGL and increasingly mature forms of IOMA might stimulate an antibody maturation pathway that is relatively insensitive to the form of N-glycan at any potential N-linked glycosylation site (PNGS) on HIV-1 Env. Promiscuous glycan recognition is desired because Env trimers on viruses exhibit heterogeneous glycosylation at single PNGSs, even within one HIV-1 strain (*43–45*). Using yeast display for immunogen selection, we sought to achieve promiscuous N-glycan accommodation through recognition of a glycan’s core pentasaccharide, a common feature of both complex-type and high-mannose glycans, which we observed as being recognized at some N-glycan sites in structures of antibody Fab-Env complexes (*21, 46, 47*).

**Figure 1.**
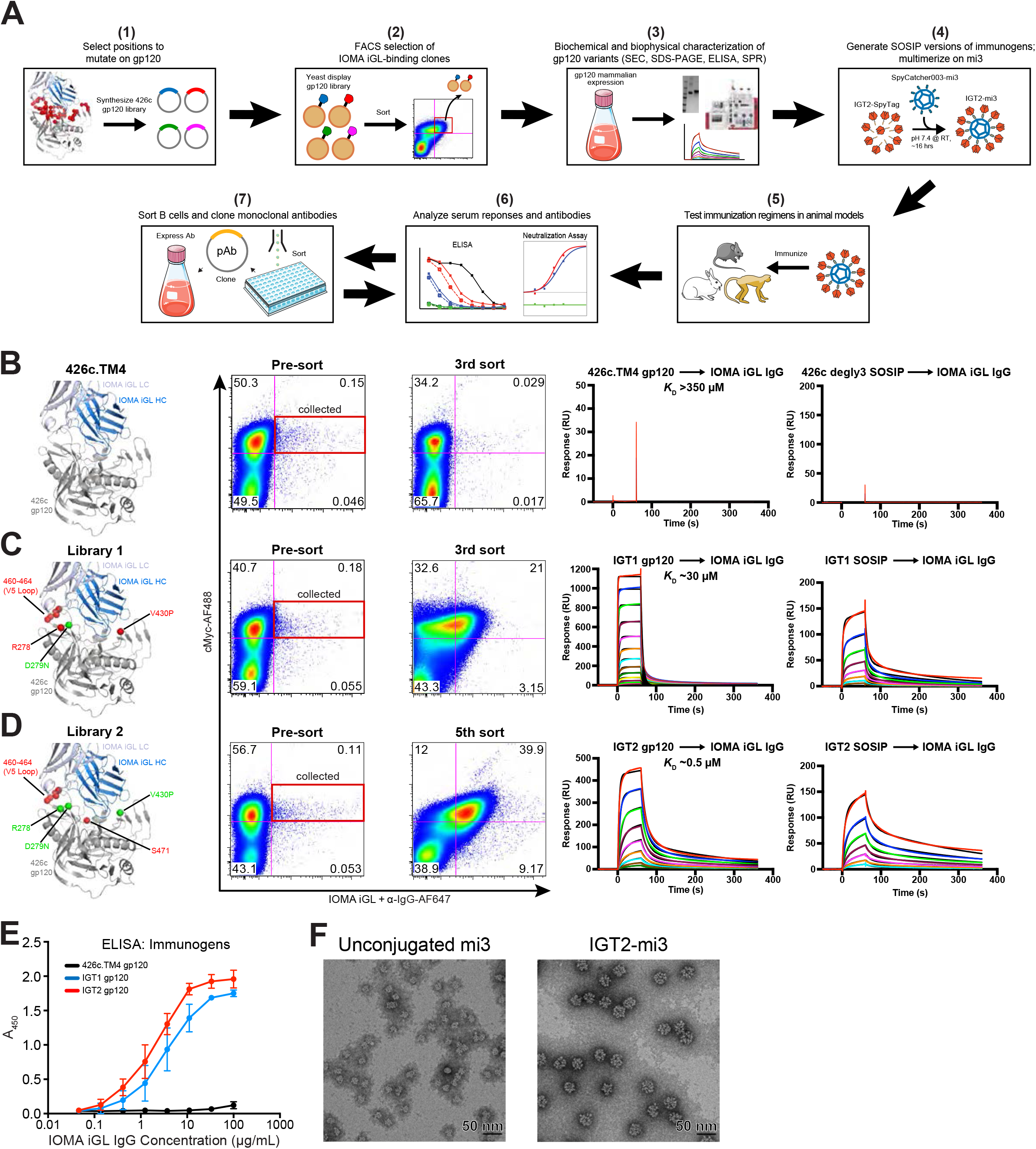
Design and characterization of IOMA-iGL targeting immunogens. **(A)** Overview of strategy to engineer and test immunogens designed to elicit IOMA-like antibodies: **(B-D)** Residues selected from the unmutated starting protein 426c.TM4 gp120 to mutate in yeast display library (left panel), FACS summary (second and third panels), and SPR data for highest affinity immunogen selected from each library as gp120 (fourth panel) and SOSIP (fifth panel) are shown **(B)**, Library 1 **(C)**, and Library 2 **(D)**. Residues shown in red represent degenerate positions in the library, while residues shown in green represent point mutations from 426c.TM4 gp120. Representative sensorgrams are shown in red with the 1:1 binding model fits shown in black. IgG was immobilized to the CM5 chip and gp120 at varying concentrations was flowed over the chip surface (IGT2 gp120: 4.9 nM – 5,000 nM; IGT1 gp120: 2.3 nM – 150,000 nM; 426c.TM4 gp120: 7,000 nM – 609,000 nM; IGT2 SOSIP: 31 nM – 2,000 nM; IGT1 SOSIP: 78 nM – 10,000 nM; 426c degly2 SOSIP: 313 nM – 40,000 nM). **(E)** ELISA demonstrating binding of CD4bs IgGs to various Env proteins. Bars indicate mean and 95% confidence interval. **(F)** Representative negative stain EM micrographs of unconjugated SpyCatcher003-mi3 nanoparticles (left) and IGT2-SpyTag SOSIP conjugated to SpyCatcher003-mi3 nanoparticles (right). Scale bar is 50 nm.

Yeast display libraries were produced using variants of the 426c.NLGS.TM4ΔV1-3 monomeric gp120 immunogen (hereafter referred to as 426c.TM4 gp120), a modified clade C gp120 that was designed to engage VRC01-class precursor antibodies (*25, 48, 49*). We started with a gp120-based immunogen instead of the engineered outer domain immunogens (eODs) previously used to select for VRC01-class bNAb precursors (*27, 29, 50*) because, unlike certain other CD4bs bNAbs, IOMA contacts the inner domain of gp120 (*34*), which is absent in the eOD constructs (*27, 29, 50*). To aid in determining which immunogen residues should be varied to achieve IOMA iGL binding, we solved a 2.07 Å crystal structure of IOMA iGL Fab (Figure S1D, Table S2), which was nearly identical (root mean square deviation, RMSD, of 0.64 Å for 209 Cα atoms) to the mature IOMA Fab structure complexed with BG505 Env trimer (*34*) (Figure S1E). Based on modeling the IOMA iGL Fab structure (Figure 1B-C, Figure S1D) into the mature IOMA Fab-Env structure (*34*), we varied 7 positions in 426c.TM4 gp120. A library with ∼10^8^ variants was produced using degenerate codons so that all possible amino acids were incorporated at the selected positions (see methods). R278_gp120_ was varied because this position might select for IOMA iGL’s unique CDRL1 conformation (*34*). In addition, we introduced a D279N_gp120_ substitution because IOMA is ∼2-3-fold more potent against HIV-1 viruses that have an N at this position (*34*). Next, V430_gp120_ was varied to increase the interaction with the HC of IOMA iGL. Lastly, residues 460_gp120_-464_gp120_ were varied in the V5-loop of 426c gp120 to accommodate and select for IOMA iGL’s normal length CDRL3.

Following three rounds of fluorescence-activated cell sorting (FACS) using one fluorophore for IOMA iGL and another against a C-terminal Myc tag to monitor gp120 expression, there was a > 100-fold enrichment for gp120 variants that bound IOMA iGL (Figure S1F, middle panel), demonstrated by increased staining for IOMA iGL compared to the starting 426c.TM4 gp120 (Figure 1B-C, Figure S1F, left and middle panels). Two clones (from ∼100 sequenced after the third sort) accounted for 50% of the sequences, suggesting that IOMA iGL-binding activity was enriched. IGT1, the best variant identified by the initial yeast display library, had an affinity of ∼30 µM for IOMA iGL, as determined by a surface plasmon resonance (SPR)-based binding assay (Figure 1C, right panel). IGT1 was then used as a guide to construct a second yeast library to select for an immunogen with higher affinity to IOMA iGL (Figure 1D). Based on their selection in IGT1, we maintained residues R278_gp120_, N279_gp120_, and P430_gp120_, while allowing amino acids R/N/K/S to be sampled at position 460. In addition, residues 461-464 and 471 were allowed to be fully degenerate and sample all possible amino acids. Following 7 rounds of sorting, multiple clones were selected including IGT2, which bound to IOMA iGL with a 0.5 µM affinity (Figure 1D, right panel and Figure S1F, right panel).

IOMA-targeting mutations selected by yeast display were transferred onto a 426c soluble native-like Env trimer (a SOSIP.664 construct (*33*)) to hide potentially immunodominant off-target epitopes within the Env trimer core that are exposed in a monomeric gp120 protein. The SOSIP versions of IGT1 and IGT2 were well behaved in size-exclusion chromatography and SDS-PAGE (Figure S1G-H). IGT1 and IGT2 SOSIPs bound to IOMA iGL IgG with higher apparent affinities than IGT1 and IGT2 gp120s due to avidity effects (Figure 1C-D). IGT1 and IGT2 SOSIP- and gp120-based immunogens were also evaluated for binding to a panel of VRC01-class iGL antibodies (*27*) (VRC01, 3BNC60, BG24). IGT2 bound all the iGLs tested, making it the only reported immunogen that binds to iGLs from both IOMA- and VRC01-class CD4bs bNAbs (Figure 1E, Figure S1I-J). Finally, using the SpyCatcher-SpyTag system (*51*), we covalently linked our SpyTagged SOSIP-based immunogens to the designed 60-mer nanoparticle SpyCatcher003-mi3 (*52*) (Figure 1A, 1F), thereby enhancing antigenicity and immunogencity through avidity effects from multimerization (*53, 54*) (Figure S1I), while also reducing the exposure of undesired epitopes at the base of soluble Env trimers (*55–57*). Efficient covalent coupling of the immunogens to SpyCatcher003-mi3 was demonstrated by SDS-PAGE (Figure S1H), and negative stain electron microscopy (EM) showed that these nanoparticles were densely conjugated and uniform in size and shape (Figure 1F).

### Sequential immunization of transgenic IOMA iGL knock-in mice elicits broad heterologous neutralizing serum responses

To evaluate whether our immunogens induced IOMA-like antibody responses, we generated transgenic mouse models expressing the full, rearranged IOMA iGL V_H_ or V_L_ genes in the mouse *Igh* (*Igh^IOMAiGL^*) and *Igk* loci (*Igk^IOMAiGL^*^)^ (Figure S2A-B). Mice homozygous for both chains, termed IOMAgl mice, showed overall normal B cell development with reduced numbers of pre-B cells and late upregulation of CD2 (suggesting accelerated B cell development due to the already rearranged VDJ and VJ genes), a preference for the IOMA iGL Igκ as seen by a reduction of mouse Igλ-expressing cells, and a reduction in IgD expression indicative of low autoreactivity (*58*) (Figure S2C-H). Total B cell numbers in IOMAgl mice were grossly normal, making them suitable to test IOMA germline-targeting immunogens (Figure S2D, S2G).

We primed the IOMAgl mice using mi3 nanoparticles coupled with the SOSIP version of the immunogen with the highest affinity to IOMA iGL (Figure 2A, IGT2-mi3) adjuvanted with the SMNP adjuvant (*59*), and compared binding by ELISA to IGT2 and a CD4bs knockout mutant IGT2 (CD4bs KO: G366R/D368R/D279N/A281T). Priming the IOMAgl mice with IGT2-mi3 elicited only weak responses to the priming and boosting (IGT1-mi3) immunogens (Figure 2B-C). However, boosting with mi3 nanoparticles coupled with IGT1, which bound IOMA iGL with a lower affinity than IGT2 (Figure 1C, middle panel), increased the magnitude and specificity of the serum responses, as demonstrated by an increase in binding to IGT2 and IGT1 compared to IGT2- and IGT1-CD4bs KO (Figure 2B-C). A comparable level of differential binding was preserved throughout the remaining immunizations (Group 1) following boosting with 426c degly2 D279N (degly2: removal of N460_gp120_ and N462_gp120_ PNGSs) followed by mosaic8-mi3, a nanoparticle coupled with 8 different wt SOSIPs chosen from a global HIV-1 reference panel used to screen bNAbs (*60*) (Table S1). Serum binding also increased throughout the immunization regimen for 426c and 426c D279N, a mutation preferred by IOMA, compared to 426c-CD4bs KO (Figure 2D). Terminal bleed sera showed binding to a panel of heterologous wt and N276A Env SOSIPs in (Figure 2E-F) and when screened against a panel of IOMA-sensitive HIV-1 strains, 8 of 12 IOMA iGL knock-in animals neutralized up to 9 of 15 strains (Figure 2G, Figure S3A-M, Table S3). However, one of these mice (ET34) also neutralized the MuLV control virus, suggesting that the neutralization activity from this mouse is at least partially non-specific for HIV.

**Figure 2.**
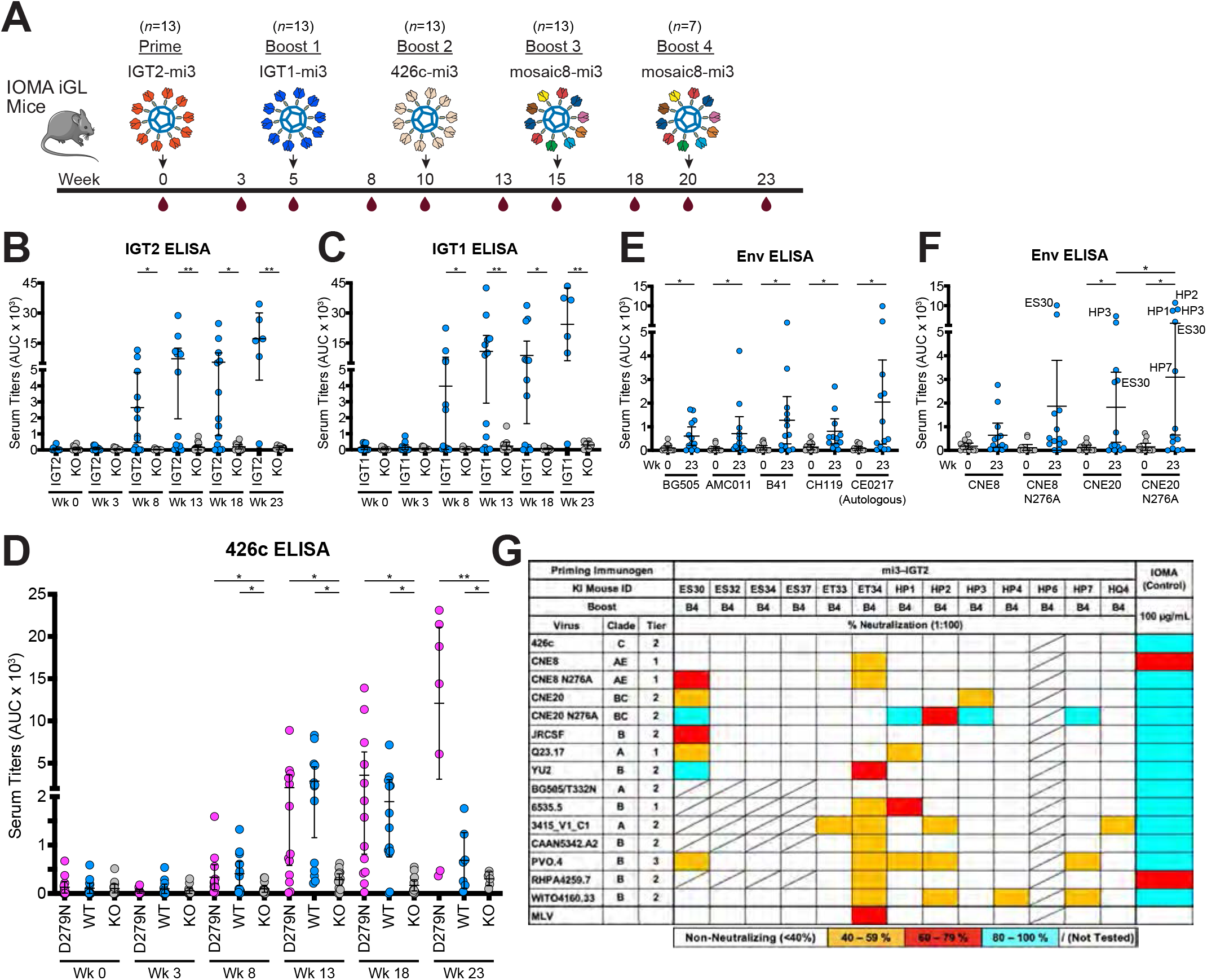
Sequential immunization with IOMA-targeting immunogens elicits heterologous neutralizing serum responses in IOMA iGL transgenic mice. **(A)** Schematic and timeline of immunization regimen for IOMA iGL knock-in mice. **(B-F)** Serum ELISA binding at the indicated time points for IGT2 and IGT2 KO **(B)**, IGT1 and IGT1 KO **(C)**, 426c D279N, 426c, and 426c KO **(D)**, and to a panel of wt and N276A-versions of SOSIP-based Envs **(E-F)**. **(G)** Serum neutralization activity against a panel of 18 HIV pseudoviruses and an murine leukemia virus (MLV) control after terminal bleed. Animal immunization studies were performed as 3 independent experiments. Each dot represents results from one mouse. Bars indicate mean and 95% confidence interval. AUC, area under the curve. *n*, number of animals.

To determine whether a shorter immunization regimen could elicit heterologous neutralizing responses, we tested 7 other immunization regimens in IOMA iGL knock-in mice (Figure S4, groups 2-8). ELISA binding titers against 426c degly2 and 426c SOSIPs using serum from group 1, which was primed with IGT2-mi3 and sequentially boosted with IGT1-mi3, 426c degly2 D279N-mi3, and mosaic8-mi3, were significantly higher than binding titers from the other groups (p: <0.0001) (Figure S4B). These results demonstrate the requirement for germline targeting through sequential immunization to induce IOMA-like antibodies.

### bNAbs isolated from IOMA iGL knock-in mice

To analyze immunization-induced antibodies, we isolated B cells from spleen and mesenteric lymph nodes of three IOMA iGL knock-in mice of group 1 (ES30, HP1, and HP3) following the final boost (Week 18 or 23, Figure 2G). We sorted immunization-induced germinal center B cells or used antigen-bait combinations of 426c degly2 D279N or CNE8 N276A together with 426c degly2 D279N-CD4bs KO (Table S1) to sort epitope-specific B cells (Figure S5). Among the identified HC and LC sequences, we noted a correlation (R^2^ = 0.78 for HCs and R^2^ = 0.62 for LCs) between the total number of V region amino acid mutations and V region mutations with identical or chemical similarity to IOMA. We compared this to unbiased VH1-2*01 or VL2-23*02 sequences derived from peripheral blood of HIV-negative human donors which showed both a lower rate and correlation (R^2^ = 0.52 for HCs and R^2^ = 0.55 for LCs) of IOMA-like mutations indicating that the immunization regimen induced maturation of IOMA iGL towards IOMA (*61*), particularly of the heavy chain which constitutes the majority of contact surface between IOMA and Env-based immunogens (*34*) (Figure 3A). 55 paired sequences were selected for antibody production based on mutation load and similarity to mature IOMA (Figure S6). In addition, 10x Genomics VDJ analysis of germinal center B cells revealed 5207 paired HC and LC sequences, of which another 12 were chosen for recombinant antibody production (Figure S5, S6, S7A-B).

**Figure 3.**
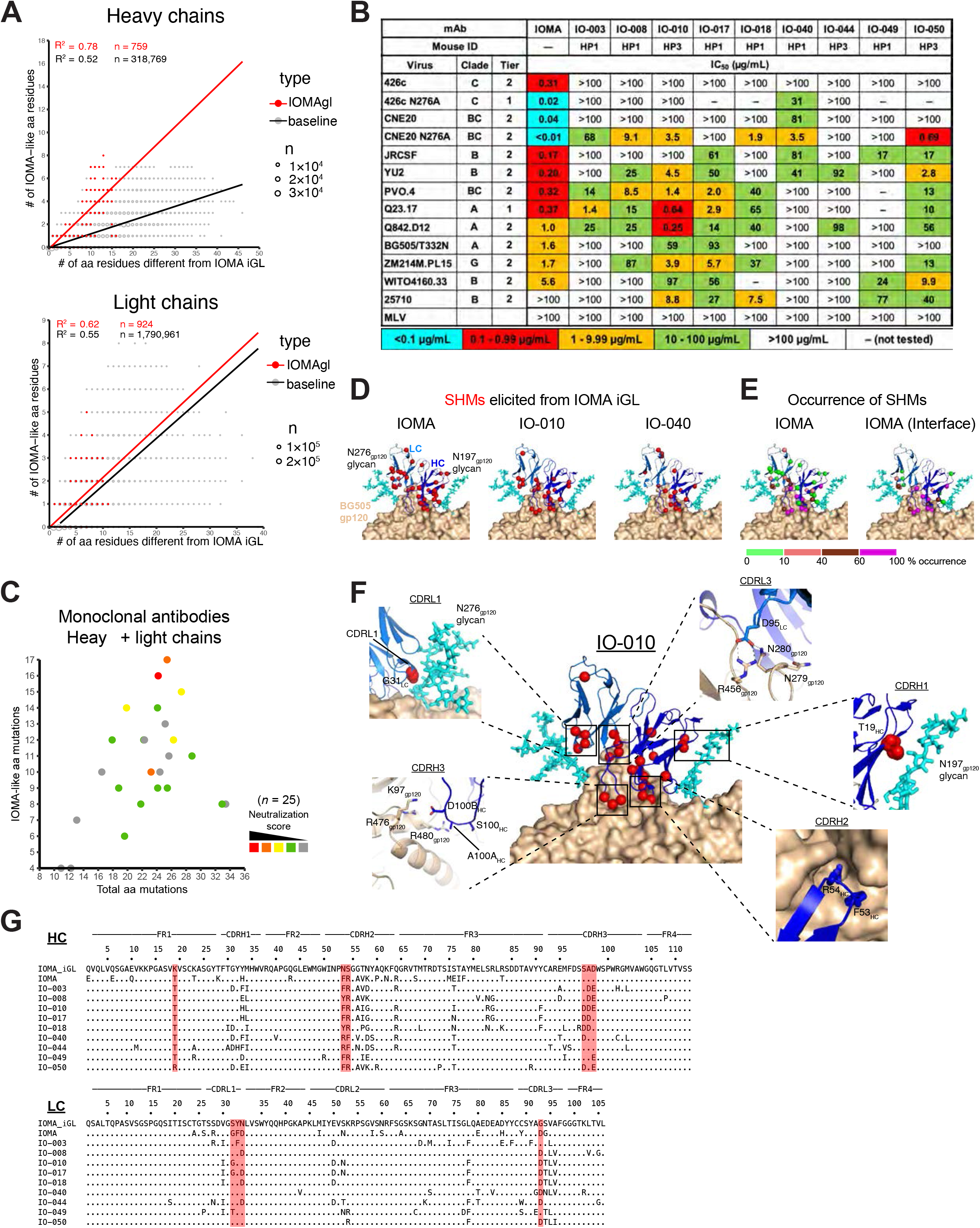
Monoclonal antibodies cloned from IOMA iGL transgenic mice neutralize heterologous HIV strains. **(A)** Graphs show the total number of V region (excluding CDR3) amino acid mutations in HC (top) and LC (bottom) of all antibody sequences (x-axis) vs. the number of mutations that are identical or chemically equivalent to mutations in IOMA for aa positions where IOMA and IOMA iGL differ (y-axis). Sequences derived from IOMAgl mice HP1, HP3 and ES30 from immunization group 1 (red) and baseline human VH1-2*01 or VL2-23*02 sequences from peripheral blood of HIV-negative human donors (gray). The size of the dot is proportional to the number of sequences. Number of sequences (n), determination coefficient (Pearson, R^2^) and linear regression lines are indicated. Chemical equivalence classified in 7 groups as follows: (1) G=A=V=L=I; (2) S=T; (3) C=M; (4) D=N=E=Q; (5)_R=K=H; (6) F=Y=W; (7) P. **(B)** Neutralization titers (IC_50_s) of nine representative monoclonal antibodies isolated from IOMA iGL transgenic mice against a panel of 14 viruses and an MLV control. IC_50_s for IOMA are shown on the far left. **(C)** 3D plot showing neutralization activity (color coded), total number of amino acid mutations in both HC and LC V(D)Js (x-axis), and the number of mutations that are identical or chemically equivalent to mutations in the IOMA (y-axis) for all Env-binding monoclonal antibodies from IOMAgl mice HP1, HP3, and ES30 from immunization group 1. Chemical equivalence is as in (A). For each antibody a neutralization score was calculated (see Methods). Red indicates higher neutralization activity and score. Number of sequences (n) are indicated. **(D)** Residues mutated from IOMA iGL are shown as red spheres mapped onto the crystal structure of mature IOMA (shown in cartoon representation) bound to BG505 gp120 (depicted in surface representation) (PDB 5T3Z). SHMs are depicted for mature IOMA (left panel) as well as two antibodies isolated from IOMAgl mice: the more potent IO-010 (middle panel) and weaker IO-040 (right panel). **(E)** Total SHMs for mature IOMA (left panel) or SHMs found in the IOMA-gp120 interface (right panel) are colored according to their percentages of occurrence from green to magenta (left panel). Structures are depicted as in (D). **(F)** Key mutations essential for IOMA binding to Env that were elicited in our immunization strategy are mapped onto antibody IO-010 and highlighted in each inset box. IO-010 depicted as in (D). Each inset represents a different interaction between IOMA and gp120. **(G)** Amino acid sequence alignment of IOMA V_H_ and V_L_ and monoclonal antibodies from (B) with IOMA iGL as a reference.

The selected monoclonal antibodies were tested for binding to a panel of heterologous Envs by ELISA (Figure S8A). Isolated IOMA-like antibodies that demonstrated binding to the Envs were then evaluated in pseudotyped *in vitro* neutralization assays (*62*), and several exhibited similar neutralization potencies as mature IOMA on a small panel of heterologous HIV-1 strains. Some antibodies neutralized the tier 2 strain 25710, which IOMA does not neutralize, and IO-010 neutralized Q842.D12 better than IOMA (Figure 3B). We also noted that among the Env-binding monoclonal antibodies, stronger neutralization activity tended to occur with antibodies that shared a larger number of critical residues with IOMA (Figure 3C).

Two mature IOMA residues, CDRH2 residues F53_HC_ and R54_HC_, interact with the CD4bs Phe43 binding pocket (*63*) on gp120 and are critical for Env recognition (*34*) (Figure 3D-G, Figure S6). 29 of 67 clones chosen for antibody production (Figure S6) contained both mutations and another 15 contained R54_HC_, 5 of which in combination with Y53_HC_, which is chemically similar to F53_HC_. N53F_HC_ is a rare mutation that is found in only ∼0.13% of VH1-2*02-derived antibodies (*64, 65*). In contrast, our immunization regimen elicited this mutation in ∼45% of antibodies, a ∼350-fold increase. S54R is elicited at slightly higher frequencies in VH1-2*02-derived antibodies (∼2.7%). However, our immunization regimen elicited this mutation at an ∼24-fold higher rate compared to the random frequency of this mutation in VH1-2*02-derived antibodies (Table S4) (*64*). In addition, our sequential immunization regimen selected for a negatively-charged DDE motif in CDRH3 (replacing the IOMA sequence of S100, A100A and D110B) in 23 of 63 sequences and another 27 sequences with at least 1 of the 3 mutations, which was likely selected for by a highly conserved patch of positively-charged residues found at the IOMA-contacting interface of the Envs used in our immunization regimen [K97_gp120_ (90% conserved), R476 _gp120_ (R - 64% conserved, R/K - 98% conserved), and R480_gp120_ (99% conserved)] (Figure 3D-G, Figure S6). To accommodate the N276_gp120_ glycan, IOMA acquired 3 mutations in CDRL1 (S29G_LC_, Y30F_LC_, N31D_LC_). The group 1 immunization regimen elicited all 3 of these substitutions; however, none of the clones contained all these mutations. Of 63 antibodies 7 contained two and another 25 contained one of these mutations (Figure 3D-E, Figure S6). Two of the most potent antibodies elicited by our immunization regimen, IO-010 and IO-017, acquired the S31G_gp120_ mutation, suggesting this mutation is more critical to accommodate the N276_gp120_ glycan (Figure 3D-G) and generating antibody breadth and potency. While accommodation of the N276 glycan is critical for CD4bs bNAbs to develop breadth and potency, CD4bs bNAbs must also acquire mutations to better interact with the N197 glycan, such as K19T_HC_ in FR1. Our immunization strategy elicited the K19T_HC_ mutation in 31 of 67 monoclonal antibodies (∼46%), which is ∼20-fold higher compared to the random frequency of this mutation in VH1-2*02-derived antibodies (Table S4) (*64*). Within the CDRL3, VRC01-class bNAbs acquire a G96E_LC_ mutation that enables interactions with the CD4bs loop, while IOMA acquires a similar G95D_LC_ mutation. Once again, this mutation was elicited in 22 of 67 antibodies (∼33%) by our immunization regimen (Figure 3D-G, Figure S6). An essential interaction of VRC01-class bNAbs involves the germline-encoded N58 residue in FR3_HC_, which makes backbone contacts to the highly (∼95%) conserved R456_gp120_. Due to a shift away from gp120 in CDRL2, IOMA acquires an N58K_HC_ substitution such that the longer lysine sidechain can access R456_gp120_ (*34*). Our immunization regimen elicited substitutions at N58_HC_ to amino acids with longer sidechains in 39 of 67 (∼58%) antibodies and was mutated to N58K_HC_ in 17 of 67 (∼24%) antibodies, a ∼1.5-fold increase over the random frequency of the N58K_HC_ mutation (Figure 3D-G, Figure S6, Table S4). Our immunization regimen elicited additional IOMA-like mutations within CDRH2: G56A_HC_ (∼30%) and T57V_HC_ (∼31%), ∼3-fold and ∼74-fold increases over the random frequency in other VH1-2*02-derived antibodies (Table S4). The 10x Genomics VDJ analysis produced an unbiased view of the extent of SHM elicited in the germinal center over the course of the immunization regimen, which, excluding frame shifts, reached up to 26 amino acid mutations in the HC exceeding the number of mutations of the IOMA HC and up to 10 mutations in the LC (Figure S7C-E).

### Sera from prime-boosted wt mice targeted the CD4bs and displayed heterologous neutralizing activity

We next investigated the same immunization regimen in wt mice (Figure 4A). Since IOMA does not have the same sequence requirements as VRC01-class bNAbs (*34*), we hypothesized that a prime-boost with IGT2-IGT1 could induce IOMA-like antibodies (which we define as recognizing the CD4bs and including a normal-length CDRL3 (*34*)) in wt mice, even though these mice do not contain the VH1-2 germline gene segment. Priming with IGT2-mi3 in wt mice elicited strong serum binding responses that were CD4bs-specific (Figure 4B, p ≤ 0.05), compared to the IOMA iGL knock-in mice, which only responded robustly after boost with IGT1-mi3 (Figure 2B-C). As in the IOMA iGL knock-in mice, the magnitude of these responses increased after boosting with IGT1-mi3, and importantly, a significant fraction of the response was still epitope-specific (p ≤ 0.001). To characterize antibodies in immunized serum, we measured binding to anti-idiotypic monoclonal antibodies raised against IOMA iGL. While naïve serum did not react with either of the anti-idiotypic antibodies, priming with IGT2-mi3 elicited serum responses that bound both anti-idiotypic antibodies, and boosting with IGT1-mi3 increased these responses (Figure 4C). After further boosting with 426c-mi3 and mosaic8-mi3 (Figure 4A), we measured binding to heterologous wt Envs. Our immunization regimen elicited significantly increased binding responses to all 9 Envs (Figure 4D-E, p ≤ 0.05 to < 0.001) in the majority of mice. Importantly, serum binding to CNE8 N276A_gp120_ and CNE20 N276A_gp120_ was significantly higher compared to CNE8 and CNE20 (p ≤ 0.05), suggesting that these responses were at least partially specific to the CD4bs (Figure 4E). Finally, we evaluated neutralization activity against a panel of heterologous HIV-1 strains and detected weak heterologous neutralization in the serum of 7 of 16 wt animals (Figure 4F, Figure S3 N-X, Table S5).

**Figure 4.**
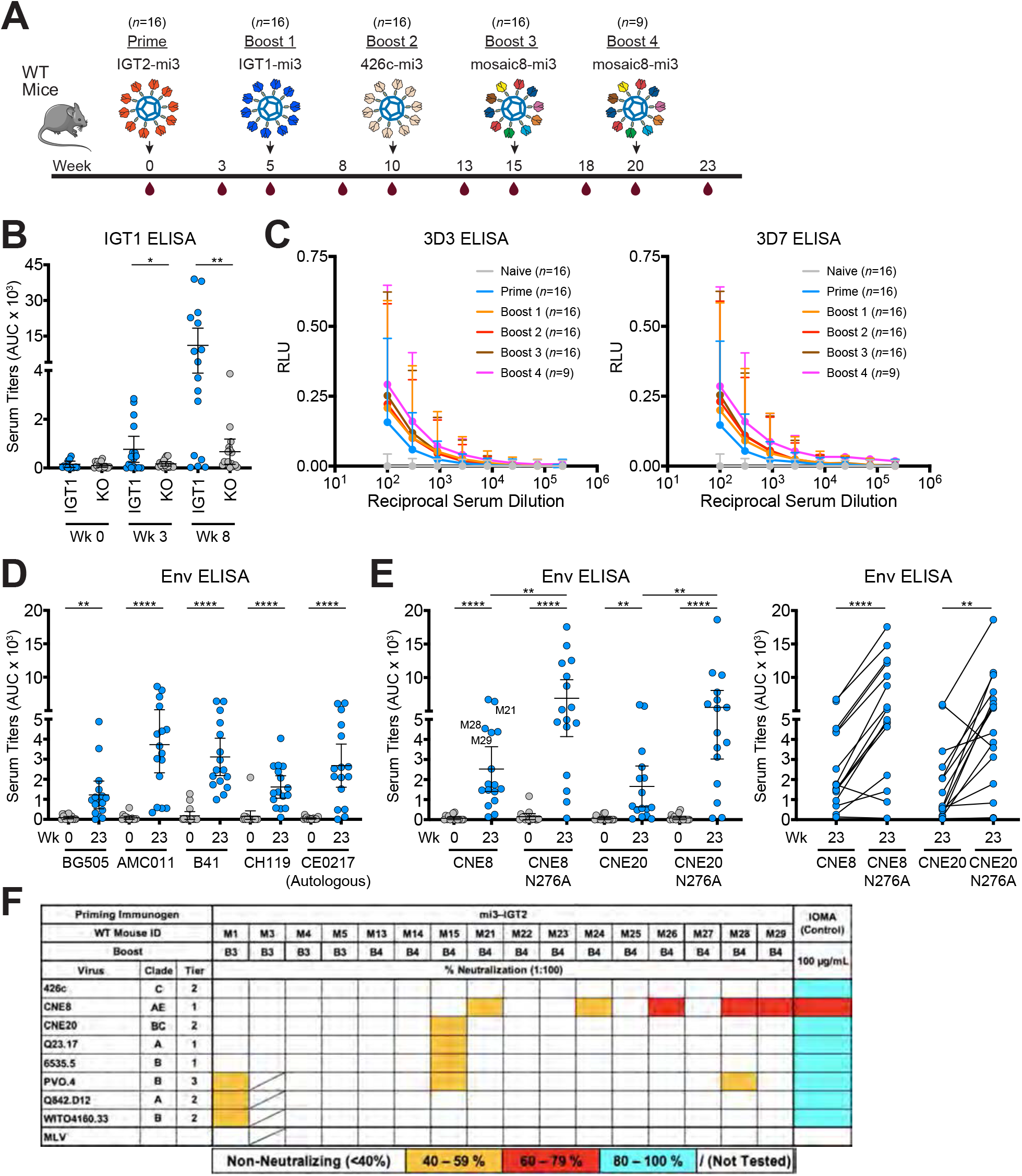
Sequential immunization with IOMA-targeting immunogens elicits CD4bs-specific responses and heterologous neutralizing serum responses in wildtype mice. **(A)** Schematic and timeline of immunization regimen for wt mice. **(B)** Serum ELISA binding at the indicated time points to IGT1 or IGT1 CD4bs-KO (KO). **(C-D)** Serum ELISA binding to anti-idiotypic monoclonal antibodies raised against IOMA iGL (left, 3D3) and IOMA iGL + mature IOMA (right, 3D7). Mean ± SEM of 9 to 16 mice per time point are depicted. **(D-E)** Serum ELISA binding at the indicated time points to a panel of WT and N276A-versions of SOSIP-based Envs. **(F)** Serum neutralization against a panel of 18 viruses and an MLV control at week 23 of wt mice. Animal immunization studies were performed as 3 independent experiments. Each dot represents results from one mouse. Bars indicate mean and 95% confidence interval. AUC, area under the curve.

### Immunization of rabbits and rhesus macaques elicited CD4bs-specific responses

To evaluate this immunization regimen in other wt animals with more potential relevance to humans, we started by immunizing rabbits and rhesus macaques with IGT2-mi3 followed by IGT1-mi3 (Figure 5A). For these experiments, we assayed only for binding antibody responses since we did not achieve heterologous neutralization after a prime or a prime/single boost of a different HIV-1 immunogen in rabbits or non-human primates (NHPs) (*57, 66*). As with the wt mouse immunizations, the IGT2-mi3 immunization elicited robust responses that were partially epitope-specific as evaluated by comparing binding to IGT1 versus to IGT1-CD4bs KO (Figure 5B). When boosted with IGT1-mi3, the responses showed significant increases in epitope specificity to the CD4bs in both rabbits and non-human primates (NHPs) (p ≤ 0.05) (Figure 5B). In addition, post-prime and post-boost sera exhibited potent neutralization of pseudoviruses generated from the IGT2 and IGT1 immunogens (Figure 5C). As stated above, we did not evaluate neutralization of heterologous pseudoviruses since our previous results using a different HIV-1 immunogen in rabbits and NHPs showed heterologous neutralization only after a second boost (*57*). The increase in epitope specificity and serum neutralization titers following boosting with IGT1 suggests that our immunization strategy is well optimized to elicit CD4bs antibody responses.

**Figure 5.**
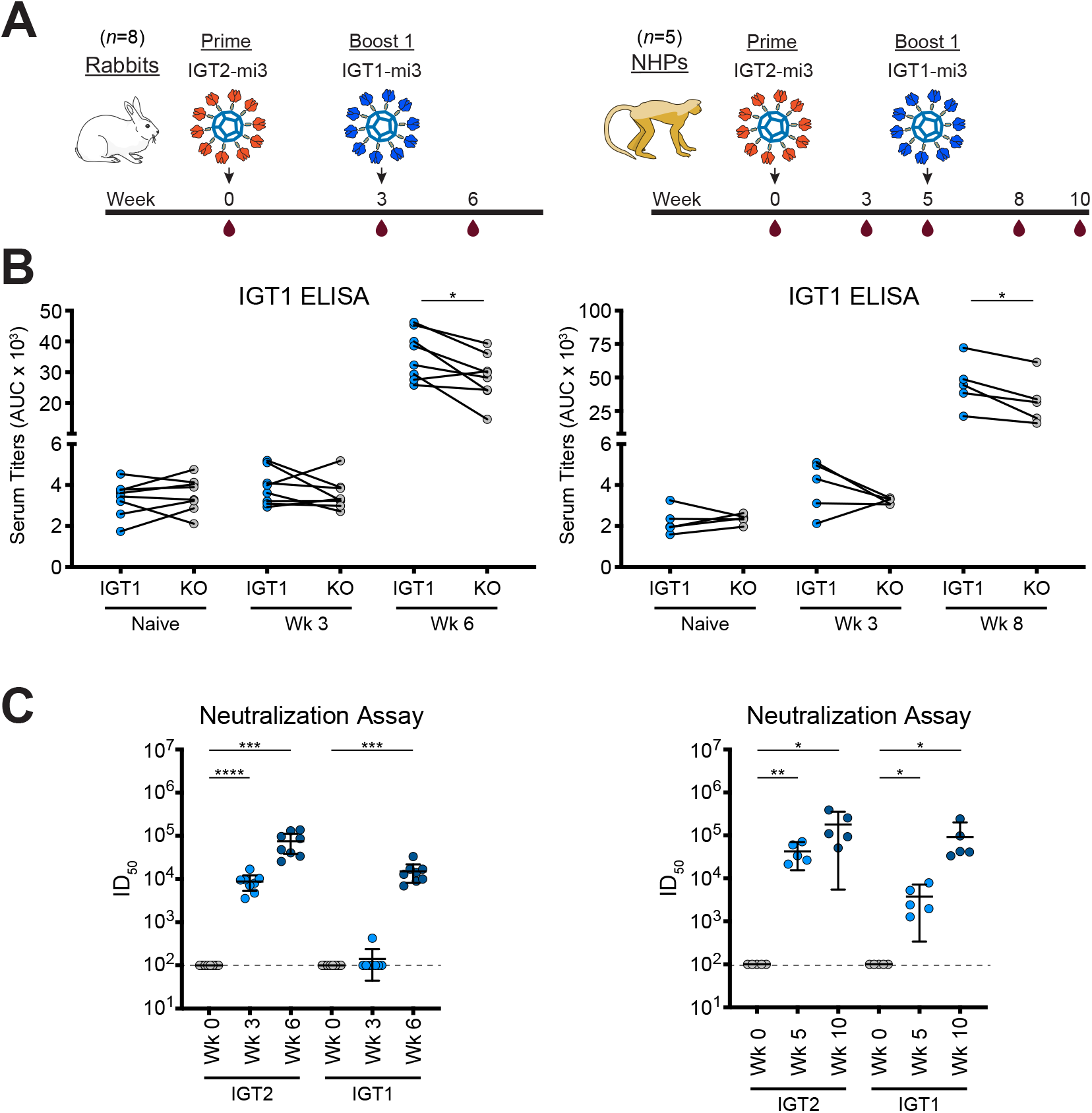
Prime-boost with IGT2-IGT1 elicits CD4bs-specific responses and potent autologous neutralization in rabbits and rhesus macaques. **(A)** Schematic and timeline of immunization regimen for rabbits and rhesus macaques. **(B)** Serum ELISA binding to IGT1 and IGT1 KO for rabbits (left) and rhesus macaques (right). **(C)** Serum neutralization ID_50_s of IGT2 and IGT1 pseudoviruses for rabbits (left) and rhesus macaques (right). The dotted line at y = 10^2^ indicates the lowest dilution evaluated. Significance was demonstrated using a paired t test (p ≤ 0.05).

## Discussion

Here we describe an immunization regimen to elicit antibodies to the CD4bs epitope on HIV Env using engineered immunogens targeting IOMA-like CD4bs antibody precursors. The ultimate goal of the germline-targeting approach is the induction of bNAbs at protective concentrations (*1*), but to date, no study has been able to accomplish this feat, although a recent study involving mRNA delivery of HIV-1 Env and gag genes reported reduced risk of SHIV infection in immunized NHPs (*67*). A previous study using a transgenic mouse expressing diverse VRC01 germline precursors demonstrated that priming with eOD-GT8 followed by sequential boosting with more native-like Envs elicited VRC01-like bNAbs (*38*). However, that study required 9 immunizations over 81 weeks to elicit VRC01-class antibodies with heterologous neutralization. By comparison, our study elicited bNAbs with similar breadth and potency using only 4-5 immunizations in 18-23 weeks. In addition, sequence analysis of the monoclonal antibodies elicited in the IOMA iGL transgenic mice revealed that our immunization regimen was much more efficient at eliciting critical mutations required for bNAb development compared to the immunogens used in the attempts to elicit VRC01-class bNAbs (*38*). Finally, the neutralization profiles of monoclonal antibodies often correlated with serum neutralization from the mouse they were isolated from. For example, IO-010 and IO-017, which neutralized PVO.4 and Q23.17, were isolated from HP3 and HP1, whose serum also demonstrated neutralization activity against these strains (Figure 2G and Figure S3A-M).

Accommodation of the N276_gp120_ glycan is considered the major impediment to the elicitation of bNAbs targeting the CD4bs (*32*). To accommodate the N276 _gp120_ glycan, VRC01-class bNAbs require a 2 - 6 residue deletion or the selection of multiple glycines within CDRL1 (*32*). IOMA requires simpler substitution of 4 residues in CDRL1 (S27AR_LC_, S29G_LC_, Y30F_LC_, N31D_LC_) (*34*). These mutations were elicited in our immunization regimen, although no single clone contained all 4 of these residues. The two most potent monoclonal antibodies isolated from immunized iGL mice, IO-010 and IO-017, contained the S31G mutation, suggesting this residue is most critical for accommodating the N276_gp120_ glycan in IOMA-like antibodies and to the development of bNAbs capable of potent heterologous neutralization. Although these antibodies were cloned from mice following the 4^th^ or 5^th^ immunization, sera from week 8 of our immunization regimen displayed significant binding to 426c Envs containing the N276_gp120_ glycan (Figure 2D), suggesting these mutations were elicited following only two immunizations. In contrast, in the same study noted above (*38*), mutations within CDRL1 of VRC01 required to accommodate the N276_gp120_ glycan occurred only after the ninth immunization at 81 weeks (Figure S8B). Additional mutations known to be important for binding to the CD4bs were also elicited earlier and at higher efficiencies in our immunization regimen compared to previous studies (Figure S8B). Importantly, no other reported vaccination regimen to elicit CD4bs antibodies has elicited all of the required SHMs to accommodate the N276_gp120_ glycan (*23, 37, 38, 68–70*), making our results an important achievement in the pursuit to elicit CD4bs bNAbs, although these mutations need to be elicited more efficiently and at higher frequencies in a protective vaccine. Since the CDRL1 of IOMA iGL was already in a helical conformation, the CDRL1 of the IOMA precursor cells selected by priming and boosting with IGT2 and IGT1 might have been in a conformation that allowed it to accommodate the N276_gp120_ glycan and therefore not required additional SHMs to accommodate the N276_gp120_ glycan introduced in the third immunization using 426c. Thus, boosting with Envs that incorporate only high-mannose glycans at N276_gp120_ followed by boosting with Envs that only incorporate complex-type glycans at N276_gp120_ starting at the second or third immunizations might force IOMA precursor cells to adapt to more diverse and branching glycan moieties and acquire these critical SHMs.

Utilizing the strategy that we developed in IOMA iGL knock-in mice, we immunized wt mice with the same immunization regimen (Figure 4A). A prime-boost sequence with IGT2-mi3 (prime) and IGT1-mi3 (boost) elicited robust CD4bs-specific responses. Importantly, the antibodies elicited by these immunogens resembled IOMA based on binding to an anti-idiotypic antibody raised against IOMA iGL using previously described methods (*71, 72*). Subsequent immunization with more native-like Envs, 426c degly2 and mosaic8, generated serum responses capable of neutralizing heterologous HIV strains. Importantly, serum neutralization correlated with ELISA binding titers; e.g., mice that elicited the highest serum binding titers against CNE8 (M21, M28, and M29) also elicited heterologous neutralizing activity against CNE8 pseudovirus. To our knowledge, these results represent the first time CD4bs-specific responses and heterologous neutralization were elicited in wt mice, thereby setting a new standard by which to evaluate HIV immunogens in wt mice, although additional work is required to determine whether antibodies targeting the CD4bs were responsible for the neutralizing responses. Due to the success of our immunogens in wt mice, we tested them in additional animals with polyclonal antibody repertoires - rabbits and rhesus macaques. Once again, our priming immunogens elicited CD4bs-specifc responses in both animal models, representing the first time a germline-targeting immunogen designed to target CD4bs Abs elicited epitope-specific responses in rabbits and rhesus macaques.

As a final boost, we used a mosaic8 nanoparticle presenting eight different wt Envs on the surface, with the intention of more efficiently selecting cross-reactive B cells and increasing neutralization breadth, a strategy that was employed to elicit cross-neutralizing responses to influenza or to zoonotic coronaviruses of potential pandemic interest (*73, 74*). Indeed, serum isolated from both wt and transgenic mice after a mosaic8-mi3 boost bound to heterologous Envs in ELISAs and neutralized a panel of heterologous HIV pseudoviruses, although additional experiments need to be completed to determine whether the cross-neutralization was due to boosting with mosaic8-mi3.

Although a previous study suggested using gp120 cores as an important intermediate immunization step (*38*), our approach resulted in heterologously-neutralizing antibodies using trimeric SOSIP-based Envs for all immunizations. This is an important distinction, since using trimeric Envs provides the additional benefit of simultaneous targeting of multiple bNAb epitopes. Indeed, a protective HIV-1 vaccine will most likely require the elicitation of bNAbs to multiple epitopes to prevent escape from the host immune response during early infection to enable clearing of the virus. Thus, our immunogens provide a scaffold upon which to engineer other epitopes to initiate germline-targeting of additional bNAb precursors.

IOMA’s relatively lower number of SHMs and normal-length CDRL3 (*34*) suggest that eliciting IOMA-like bNAbs by vaccination might be easier to achieve, compared with eliciting VRC01-class bNAbs. Indeed, the fact our IOMA immunogens elicited CD4bs-specific responses in four animal models suggests that germline-targeting immunogens designed to elicit IOMA-like antibodies are an attractive route to generate an HIV-1 vaccine, which is supported by our engineered immunogens eliciting epitope-specific responses in wt animals and by a commonality of the mutations that were induced across individual transgenic mice. Furthermore, IOMA-like bNAbs have been isolated from multiple patients (*34, 75*), suggesting an immunization regimen targeting this class of bNAbs could be universally effective in a global population. Although IOMA’s neutralization breadth is smaller than that of other bNAbs, the fact that some vaccine-elicited IOMA-like antibodies neutralized strains that IOMA neutralizes less potently or does not neutralize at all suggests that it is possible to create polyclonal serum responses that include individual antibodies with more breadth than IOMA. If elicited at sufficient levels, such antibodies could mediate protection from more strains than predicted by the original IOMA antibody. This is an important property of a potential active vaccine, since clinical trials to evaluate protection from HIV-1 infection by passive administration of VRC01 in humans demonstrated a lack of protection from infection by HIV-1 strains against which the VRC01 exhibited weak *in vitro* potencies (*28*). Although polyclonal antibodies raised against the CD4bs may be more protective than a single administered monoclonal anti-CD4bs antibody, a successful HIV-1 vaccine will likely require broader and more potent responses to the CD4bs and other epitopes on HIV-1 Env. Our results provide new germline-targeting immunogens to build upon, demonstrate that IOMA-like precursors provide a new starting point to elicit CD4bs bNAbs and suggest that eliciting this class of bNAbs should be further pursued as a possible strategy to generate a protective HIV-1 vaccine.

### Data and materials availability

The structure of IOMA iGL Fab is available in the Protein Data Bank under accession code 7TQG. 10x Genomics VDJ sequencing data is available from Gene Expression Omnibus Accession number GSE197951. All other data, mice and reagents used in this study are available from the corresponding authors upon reasonable request. Antibody HC and LC genes were analyzed using our previously described IgPipeline (*76, 77*). The code for the IgPipeline is available at https://github.com/stratust/igpipeline/tree/igpipeline2_timepoint_v2. Further information and reasonable requests for reagents and resources should be directed to Pamela J. Bjorkman.

## Acknowledgements

We thank J. Moore (Weill Cornell Medical College), R.W. Sanders and M.J. van Gils (Amsterdam UMC) for SOSIP expression plasmids; M. Silva, M. B. Melo and D. J. Irvine (MIT) for providing SMNP adjuvant; Lotta von Boehmer (Stanford University) for discussion; J. Vielmetter, P. Hoffman, and the Protein Expression Center in the Beckman Institute at Caltech for expression assistance; T. Eisenreich and S. Tittley for animal husbandry and K. Gordon and K. Chosphel for fluorescence-activated cell sorting at Rockefeller University. Electron microscopy was performed in the Caltech Cryo-EM Center with assistance from S. Chen and A. Malyutin. This work was supported by the National Institute of Allergy and Infectious Diseases (NIAID) Grants HIVRAD P01 AI100148 (to P.J.B. and M.C.N.), HIVRAD P01 AI138212 (to L.S., A.T.M., and M.N.), and R21 AI127249 (to A.T.M.), the Bill and Melinda Gates Foundation Collaboration for AIDS Vaccine Discovery (CAVD) grant INV-002143 (P.J.B., M.C.N., and M.A.M.), a Bill and Melinda Gates Foundation grant # OPP1146996 (to M.S.S.), the Intramural Research Program of the NIAID (to M.A.M. and Y.N.), and NIH P50 AI150464 (P.J.B.). A.T.D. and M.E.A. were supported by NSF Graduate Research Fellowships. M.C.N. is an HHMI investigator. Under the grant conditions of the Bill and Melinda Gates Foundation Collaboration, a Creative Commons Attribution 4.0 Generic License has already been assigned to the Author Accepted Manuscript version that might arise from this submission.

## METHODS

### Antibody, gp120, and Env trimer expression and purification

Env immunogens were expressed as soluble SOSIP.664 native-like gp140 trimers (*33*) as described (*66*). For SpyTagged trimers, either SpyTag (13 residues) (*78*) or SpyTag003 (16 residues) (*79*) was added to the C-terminus to allow formation of an irreversible isopeptide bond to SpyCatcher003 moieties. All soluble SOSIP Envs were expressed by transient transfection in HEK293-6E cells (National Research Council of Canada) or Expi293 cells (Life Technologies) and purified from transfected cell supernatants by 2G12 affinity chromatography. Soluble Envs were stored at 4°C in 20 mM Tris pH 8.0, 150 mM sodium chloride (TBS) (untagged and AviTagged versions) or 20 mM sodium phosphate pH 7.5, 150 mM NaCl (PBS) (SpyTagged versions). We also expressed untagged gp120 proteins as cores with N/C termini and V1/V2/V3 loop truncations as described (*63*) by transient transfection of suspension-adapted HEK293-S cells. gp120s were purified using Ni-NTA affinity chromatography and Superdex 200 16/60 SEC. Proteins were stored in 20 mM Tris, pH 8.0, 150 mM sodium chloride.

The iGL sequences of IOMA was derived as described in the main text. The iGL sequences of VRC01 and 3BNC60 were derived as described (*27, 80*). The iGL of BG24, a VRC01-class bNAb with relatively few SHMs (*81*), was derived as described (*82*). IgGs were expressed by transient transfection in Expi293 cells or HEK293-6E cells and purified from cell supernatants using MabSelect SURE (Cytiva) columns followed by SEC purification using a 10/300 or 16/600 Superdex 200 (GE Healthcare) column equilibrated with PBS (20 mM sodium phosphate pH 7.4, 150 mM NaCl). His-tagged Fabs were prepared by transient transfection of truncated heavy chain genes encoding a C-terminal 6x-His tag with a light chain expression vector and purified from supernatants using a 5 mL HisTrap colum (GE Healthcare) followed by SEC as described above.

### Generation of anti-idiotypic monoclonal antibodies

Mice were injected three times with purified IOMA iGL. 3 days after the final injection spleens were harvested and used to generate hybridomas at the Fred Hutchinson Antibody Technology Center. Hybridoma supernatants were initially screened against IOMA iGL to identify antigen-specific hybridomas. Supernatants from positive wells were then screened against a panel of monoclonal antibodies that included IOMA, IOMA iGL, and inferred germlines of other anti-HIV-1 antibodies that served as isotype controls using a high throughput bead array. We identified two hybridomas of interest; 3D3, which bound specifically to IOMA iGL, and 3D7, which bound to IOMA and IOMA iGL, which were subcloned from single cells. To produce recombinant anti-idiotypes, RNA was extracted from 1 × 10^6^ cells using the RNeasy kit (Qiagen), and the heavy and light chain sequences of the murine hybridomas were by obtained using the mouse Ig-primer set (69831; EMD Millipore) as described (*83*). Sequences were codon optimized, cloned into pTT3-based IgG expression vectors with human constant regions (*84*) using In-Fusion cloning (Clontech), expressed in 293 cells, and purified using Protein A chromatography.

### X-ray crystallography

Crystallization screens for IOMA iGL Fab were performed using the sitting drop vapor diffusion method at room temperature (RT) by mixing 0.2 µL Fabs with 0.2 µL of reservoir solution (Hampton Research) using a TTP Labtech Mosquito automatic microliter pipetting robot. IOMA iGL Fab crystals were obtained in 20% (v/v) PEG 2000, 0.1 M Sodium Acetate (pH 4.6). Crystals were looped and cryopreserved in reservoir solution supplemented with 20% glycerol and flash frozen in liquid nitrogen.

The crystal structure of IOMA iGL Fab was solved with data sets. A 1.9 Å-resolution structure of IOMA – 10-1074 – BG505 was solved with a single data set collected at 100 K and 1 Å resolution on Beamline 12-2 at the Stanford Synchrotron Radiation Lightsource (SSRL) with a Pilatus 6M pixel detector (Dectris) that was indexed and integrated with iMosflm v7.4, and then merged with AIMLESS in the CCP4 software package v7.1.018. The structure was determined by molecular replacement using Phaser with one copy of IOMA Fab (PDB 5T3Z). Coordinates were refined with PHENIX v1.19.2-4158 (*85*) with group B factor and TLS restraints. Manual rebuilding was performed iteratively with Coot v1.0.0 (*86*). Data refinement statistics are shown in Table S2, with > 98% of the residues in the favored region of the Ramachandran plot and < 1% in the disallowed regions.

### Cloning yeast libraries

Crystal structures of IOMA in complex with BG505 SOSIP.664 (PDB ID 5T3X and 5T3Z) were analyzed to determine mutations on gp120 that potentially could be beneficial for IOMA iGL binding. In addition, we modeled the crystal structure of IOMA iGL (PDB ID 7TQG) onto 426c.TM4ΔV1-3 (426c.TM4) gp120 (PDB ID 5FEC) and selected positions within gp120 that we predicted to be favorable for IOMA iGL binding. We chose 426c.TM4ΔV1-3 (426c TM4), an engineered clade C Env previously shown to activate B cell precursors of HIV-1 bNAbs targeting the CD4bs (*25*) as the starting point for our library design.

Yeast libraries were generated as described (*87*). Specifically, to generate the libraries of 426c gp120 variants we used degenerate oligos in conjunction with an overlap assembly polymerase chain reaction (PCR) method. Overlapping primers for the PCR assembly reactions were designed using Primerize (*88*) and shown in Table S6. NNK codons (where N = A/C/G/T and K = G/T) were utilized that encode for all 20 amino acids but decrease the chances of introducing a premature stop codon. Two different DNA fragments (426c library fragment 1 and 2) were synthesized first and then linearized in a final PCR step to generate the full-length 426c gp120 library used in yeast transformation. To obtain the full-length 426c gp120, a final PCR reaction was performed in which the PCR products of the 426c Library Fragment 1 and 2 were used as a template. Primers were used with overhangs complementary to the yeast display vector pCTCON-2 necessary for the homologous recombination in yeast. Library 2 was cloned in a similar manner as Library 1, but using a different set of primers as shown in Table S6 based on results from Library 1.

### Yeast transformation

The yeast display vector pCTCON-2 was used for cell surface display of the 426c gp120 proteins in *Saccharomyces cerevisiae* (*S. cerevisiae*) strain EBY100. A primary culture of 5 mL 2x YPD (40 g/L glucose, 20 g/L peptone, 20 g/L yeast extract) media was inoculated with a single *S. cerevisiae* EBY100 colony (freshly streaked on a YPD plate) and incubated overnight in a shaker at 30 °C and 250 rpm. 100 μL of the overnight yeast *S. cerevisiae* EBY100 cultures was transferred into 5 mL 2x YPD media and incubated overnight at 30 °C, 250 rpm. The following day, 300 mL 2x YPD media was inoculated with the overnight precultures to an OD_600_ ∼0.3 and was grown until an OD_600_ ∼1.6. 3 mL of sterile filtered Tris/DTT (0.462 g 1,4-dithiothreitol in 3 mL 1 M Tris, pH 8.0) and 15 mL sterile filtered 2 M LiAc/TE (1.98 g LiAc in 10 mL of TE (10 mM Tris, 1 mM EDTA) was added and the culture incubated for 15 min at 30 °C and 250 rpm. Yeast cells were then pelleted at 3,500 g for 3 min and washed with 50 mL ice-cold sterile filtered NewE buffer (0.6 g Tris base, 91.09 g Sorbitol (1 M), 73.50 mg CaCl_2_ in ddH_2_O to a final volume of 500 mL, pH 7.5). After two additional wash steps, the pellet was re-suspended in 3 mL NewE buffer and 50 μg 426c library DNA insert and 10 μg pCTCON-2 vector (digested with NheI and BamHI) was added. 200 μL of this transformation mix was then aliquoted into pre-chilled 2 mm electroporation cuvettes (Bio-Rad) and electroporated at 1500 V with an average time constant of ∼4.5 ms using a Gene Pulser Xcell Electroporation System (Bio-Rad), which was repeated for the entire transformation mix. After electroporation, yeast cells were directly recovered with 2 mL 2x YPD media and transferred into 50 mL cold 2x YPD media (final volume up to 200 mL 2x YPD media) and grown for 1 h at 30 °C and 250 rpm. Serial dilutions of the freshly transformed yeast culture were plated on SDCAA (20 g/L glucose, 6.7 g/L Difco yeast nitrogen base, 1.4 g/L Yeast Synthetic Drop-out Medium Supplements without histidine, leucine, tryptophan and uracil, 20 mg/L uracil, 50 mg/L histidine, 100 mg/L leucine) agarose plates to test the viability and size of the library. After 1 h, the culture was removed and the cells were pelleted and resuspended in 500 mL SDCAA media + carbenicillin (100 μg/mL final concentration) and grown for two days at 30 °C and 250 rpm. To confirm the genetic diversity of the library, a yeast colony PCR was performed on the liquid culture and the PCR product was sequenced. Sequencing reactions were performed at Laragen Inc (Culver City, CA). The sequence data was analyzed using SeqMan Pro (DNASTAR, v13.02). After two days, cells were pelleted and glycerol stocks were made by suspending ∼10^9^ yeast cells in 1 mL of freezing buffer (0.335 g Yeast Nitrogen Base, 1 mL glycerol in 50 mL H_2_O, sterilized by filtration). Aliquots were flash frozen in liquid nitrogen and stored at −80 °C.

### Magnetic-activated cell sorting

Magnetic-activated cell sorting (MACS) was used to remove transformants containing stop codons. After growing up the freshly transformed cells for two days in SDCAA, cells were pelleted and induced at an OD_600_ ∼1.0 in 100 mL SGCAA-carb (SDCAA prepared with 20 g/L galactose instead of glucose and supplemented with 100 µg/mL carbenicillin final concentration) for 20 h at 20 °C and 250 rpm. Yeast cells were washed 5 times with PBSF (PBS + 0.1% bovine serum albumin (BSA)) and 10^8^ cells were incubated with 400 μL PBSF and 100 μL μMACS™ anti-c-Myc MicroBeads (Miltenyi Biotec) for 45 min on a rotator at 4 °C. Cells were then pelleted and resuspended in 5 mL PBSF and sorted using a MidiMACS Separator magnet (Miltenyi Biotec) in combination with an LS column (Miltenyi Biotec) equilibrated in PBSF. Isolated cells were then grown for 2 days in 100 mL SDCAA-carb at 30 °C and 250 rpm and then induced again with SGCAA-carb for 20 h at 20 °C and 250 rpm.

### Yeast flow cytometry and cell sorting

To prepare the yeast library for FACS analysis, cells were pelleted at 3000 rpm for 2 min and washed 5 times with PBSF. Cells were then stained at a density of 10^7^ cells/mL with 1:500 anti-c-Myc antibody conjugated to AlexaFluor488 (Abcam, ab190026) and 1 µM IOMA iGL and incubated for 1 – 2 h on a rotator at 4 °C. Cells were then washed twice with PBSF and resuspended in 200 μL PBSF with 1:1000 goat anti-human antibody conjugated to AlexaFluor647 (Abcam, ab190560) and incubated for 30 min at 4 °C. Cells were then analyzed on a MACSQuant Analyzer (Miltenyi Biotec) or sorted using an SY3200 cell sorter system (Sony). In either case, non-transformed yeast cells and single-stained transformed samples stained with either anti-cMyc or IOMA iGL IgG were used to set the gates for analysis and collection. Cells that stained double-positive for both c-Myc and IOMA iGL were collected and grown in 5 mL SDCAA-carb for 1 - 2 days at 30 °C and 250 rpm and then transferred to 100 mL SDCAA-carb for an additional 1 - 2 days at 30 °C and 250 rpm. Cells were then pelleted and resuspended in H_2_O and plated onto SDCAA-carb for 2 - 3 days at 30 °C. After multiple iterative rounds of sorting (three rounds for Library 1 and seven rounds for Library 2), sequences were recovered by colony PCR and sequence confirmed (Laragen). Primers were used with specific complementary regions to enable ligation of the linear product into the expression vector pTT5 using the Gibson assembly method for protein production. After construction, plasmids were isolated from *E.coli* using the QIAprep Miniprep kit (Qiagen) and confirmed by Sanger sequencing (Laragen).

### ELISAs

Serum ELISAs were performed using randomly biotinylated SOSIP trimers using the EZ-Link NHS-PEG_4_-Biotin kit (Thermo Fisher Scientific) according to the manufacturer’s guidelines. Based on the Pierce Biotin Quantitation kit (Thermo Fisher Scientific), the number of biotin molecules per protomer was estimated to be ∼1 - 4. Biotinylated SOSIP timers were immobilized on Streptavidin-coated 96-well plates (Thermo Fisher Scientific) at a concentration of 2 - 5 µg/mL in blocking buffer (1% BSA in TBS-T: 20 mM Tris pH 8.0, 150 mM NaCl, 0.1% Tween 20) for 1 h at RT. After washing plates in TBS-T, plates were incubated with a 3-fold concentration series of mouse, rabbit, or rhesus macaque serum at a top dilution of 1:100 in blocking buffer for 2-3 h at RT. After washing plates with TBS-T, HRP-conjugated goat anti-mouse Fc antibody (Southern Biotech, #1033-05) or HRP-conjugated goat anti-rabbit IgG Fc antibody (Abcam, ab98467) or HRP-conjugated goat anti-human multi-species IgG antibody (Southern Biotech, #2014-05) was added at a dilution of 1:8,000 in blocking buffer for 1 h at RT. After washing plates with TBS-T, 1-Step Ultra TMB substrate (Thermo Fisher Scientific) was added for ∼3 min. Reactions were quenched by addition of 1 N HCl and absorbance at 450 nm were analyzed using a plate reader (BioTek). ELISAs with gp120s and anti-idiotype monoclonal antibodies were performed as above except these proteins were immobilized directly onto high-binding 96-well assay plates (Costar) in 0.1 M sodium bicarbonate buffer (pH 9.8) at a concentration of 2 – 5 µg/mL in blocking buffer (1% BSA in TBS-T) for 2 h at RT. ELISAs with IgGs instead of serum were performed as above with a top IgG concentration of 100 µg/mL. All reported values represent the average of at least two independent experiments.

### SPR binding studies

All SPR measurements were performed on a Biacore T200 (GE Healthcare) at 20 °C in HBS-EP+ (GE Healthcare) running buffer. IgGs were directly immobilized onto a CM5 chip (GE Healthcare) to ∼3000 resonance units (RUs) using primary amine chemistry. A concentration series of monomeric gp120 core constructs (IGT2, IGT1, 426c TM4) were injected over the flow cells at increasing concentrations (top concentrations ranging from 600 µM to 10 µM) at a flow rate of 60 µL/min for 60 s and allowed to dissociate for 300 s. Regeneration of flow cells was achieved by injecting one pulse each of 10 mM glycine pH 2.0 at a flow rate of 90 µL/min. Kinetic analyses were used after subtraction of reference curves to derive on/off rates (*k*_a_/*k*_d_) and binding constants (*K*_D_s) using a 1:1 binding model with or without bulk refractive index change (RI) correction as appropriate (Biacore T200 Evaluation software v3.0). Reported affinities represent the average of two independent experiments. SPR experiments that were not used to derive binding affinities or kinetic constants were done using a single high concentration (1 µM) to qualitatively determine binding versus no binding.

### Preparation of SOSIP-mi3 nanoparticles

SpyCatcher003-mi3 particles were prepared by purification from BL21 (DE3)-RIPL *E. coli* (Agilent) transformed with a pET28a SpyCatcher003-mi3 gene (*89*) (including an N-terminal 6x-His tag) as described (*74, 90*). Briefly, cell pellets from transformed bacterial were lysed with a cell disruptor in the presence of 2.0 mM PMSF (Sigma). Lysates were spun at 21,000 g for 30 min, filtered with a 0.2 µm filter, and mi3 particles were isolated by ammonium sulfate precipitation followed by SEC purification using a HiLoad 16/600 Superdex 200 (GE Healthcare) column equilibrated with 25 mM Tris-HCl pH 8.0, 150 mM NaCl, 0.02% NaN_3_ (TBS). SpyCatcher003-mi3 particles were stored at 4 °C and used for conjugations for up to 1 month after filtering with a 0.2 µm filter and spinning for 30 min at 4 °C and 14,000 g.

Purified SpyCatcher003-mi3 was incubated with a 2-fold molar excess (SOSIP to mi3 subunit) of purified SpyTagged SOSIP (either a single SOSIP or an equimolar mixture of eight SOSIPs for making mosaic8 particles) overnight at RT in PBS. Conjugated SOSIP-mi3 particles were separated from free SOSIPs by SEC on a Superose 6 10/300 column (GE Healthcare) equilibrated with PBS. Fractions corresponding to conjugated mi3 particles were collected and analyzed by SDS-PAGE. Concentrations of conjugated mi3 particles were determined using the absorbance at 280 nm as measured on a Nanodrop spectrophotometer (Thermo Scientific).

### Electron microscopy of SOSIP-mi3 nanoparticles

SOSIP-mi3 particles were characterized using negative stain electron microscopy (EM) to confirm stability and the presence of conjugated SOSIPs on the mi3 surface. Briefly, SOSIP-mi3 particles were diluted to 20 µg/mL in 20 mM Tris (pH 8.0), 150 mM NaCl and 3 µL of sample was applied onto freshly glow-discharged 300-mesh copper grids. Sample was incubated on the grid for 40 s and excess sample was then blotted away with filter paper (Whatman). 3 µL uranyl acetate was added for 40 s and excess stain was then blotted off with filter paper. Prepared grids were imaged on a Talos Arctica (ThermoFisher Scientific) transmission electron microscope at 200 keV using a Falcon III 4k × 4k (ThermoFisher Scientific) direct electron detector at 13,500x magnification.

### Generation of IOMA-expressing RAMOS cells by CRISPR/Cas9 gene editing

A targeting vector was constructed using the NEB Hifi DNA assembly kit to clone a gBlock (IDT) into pUCmu (*91*). The gBlock (IDT) contained ∼0.5 kb homology arms to the human IgH locus which flanked an expression cassette consisting of the C_µ_ splice acceptor, the entire IOMA LC gene, a furin-GSG-P2A sequence (*92*) followed by the IOMA HC Leader-VDJ and the J_H_4 splice donor based on previously-described designs (*93*) (Figure S5C). Vectors were maxi-prepped (Machery-Nagle) for transfection.

RAMOS (RA 1) cells were purchased from ATCC (CRL-1596) and maintained in RPMI-1640 supplemented with 10 FCS, 1x antibiotic/antimycotic, 2 mM glutamine, 1 mM sodium pyruvate, 10 mM HEPES and 55 µM β-mercaptoethanol. Before transfection, cells were harvested, washed once in PBS and resuspended at 6×10^7^ cells/mL in Neon kit buffer T (ThermoFisher). Three ribonucleoprotein complexes (RNPs) were prepared using 3 different sgRNAs. AGGCATCGGAAAATCCACAG was used to target the IgH locus in the intron 3’ of *IGHJ6* to integrate the sequence flanked by the appropriate homology arms from the targeting vector; CTGGGAGTTACCCGATTGGA was used to ablate the human *IGKC* exon and CACGCATGAAGGGAGCACCG was used to ablate all functional *IGLC* genes (*IgLC1, IGL2, IGLC3* and *IGLC7*). Complexes were prepared by mixing 1.875 µL of 100 µM sgRNA with 1 µL of 61 µM Cas9 (all IDT) for a molar ratio of ∼3:1 followed by incubation for 20 min at RT. *IGH:IGK:IGL* RNPs were then mixed at a 2:1:1 v/v/v ratio. 2.6 µg targeting vector (at 4 mg/mL) were mixed with 1.5 µL *IGH:IGK:IGL* RNP mix and 11 µL RAMOS cells in buffer T. 10 µL of the final mix were transfected in a 10 µL Neon tip in a Neon device at 1350 V 30 ms 1 pulse. Cells were immediately transferred into 50 µL RAMOS medium without 1x antibiotic/antimycotic in a 48-well plate and 2 h later 450 µL full RAMOS medium was added. Cells were then cultured as before. Edited IOMA-expressing cells were bulk sorted by flow cytometry as live, singlet, CD19^+^, RC1 antigen^hi^, IgL^+^, IgK^+^ IgM^+^ (Table S7) and cultured as before. IOMA-expression was further verified by staining with 426c-, CNE8- and CNE20-derived SOSIPs and 426c-CD4bs-KO proteins to show specificity.

### Mice

C57BL/6J and B6(Cg)-Tyrc-2J/J (B6 albino) mice were purchased from Jackson Laboratories. *Igh^IOMAiGL^* and *Igk^IOMAiGL^*mice were generated with the Rockefeller University CRISPR and Genome Editing Center and Transgenic and Reproductive Technology Center in CY2.4 albino C57BL/6J-Tyrc-2J-derived embryonic stem cells. Chimeras were crossed to B6(Cg)-Tyrc-2J/J for germline transmission. *Igh^IOMAiGL^* and *Igk^IOMAiGL^* mice carry the IG*V(D)J* genes encoding the IOMA iGL HC and LC respectively. IOMA iGL LC was targeted into the *Igk* locus deleting the endogenous mouse *Igkj1* to *Igkj5* gene segments. IOMA iGL HC was targeted into the *Igh* locus and deleting the endogenous mouse *Ighd4-1* to *Ighj4* gene segments thereby minimizing rearrangement of the locus (Figure S2A-B) (*94, 95*). The constant regions of *Igh* and *Igk* remain of mouse origin. Mice were only crossed to C57BL/6J or B6(Cg)-Tyrc-2J/J or themselves and maintained at Rockefeller University and all experiments shown used double homozygous animals for *Igh^IOMAiGL^* and *Igk^IOMAiGL^* abbreviated IOMAgl mice. These mice are available upon request. Mice were housed at a temperature of 22 °C and humidity of 30 – 70% in a 12 h light/dark cycle with ad libitum access to food and water. Male and female mice aged 6 – 12 weeks at the start of the experiment were used throughout. All experiments were conducted with approval from the institutional review board and the institutional animal care and use committee at the Rockefeller University. Sample sizes were not calculated a priori. Given the nature of the comparisons, mice were not randomized into each experimental group and investigators were not blinded to group allocation. Instead, experimental groups were age- and sex-matched.

### Animal immunizations and sampling

Mice were immunized intraperitoneally with 10 µg conjugated mi3-SOSIP in 100 µL PBS with 1 U SMNP adjuvant (*59*) (kindly provided by Murillo Silva, Mariane B. Melo and Darrell J. Irvine, MIT). Serum samples were collected throughout the experiment by submandibular bleeding and animals were terminally bled under isoflurane anesthesia first submandibularly followed by cardiac puncture. Spleen and mesenteric lymph nodes were dissected, mashed though a 70 µm cell strainer and frozen in FCS with 10% DMSO in a gradual freezing (∼1 °C/min) container, followed by transfer to liquid N_2_ for long-term storage.

Eight six-month-old New Zealand White rabbits (LabCorp) were used for immunizations. Rabbits were immunized subcutaneously with 50 µg of a SOSIP-mi3 in SMNP adjuvant (375 U/animal) as described (*66, 96*). Serum samples were collected from rabbits at the time points indicated in Figure 5A. Procedures in rabbits were approved by the Denver PA IACUC Committee.

Five rhesus macaques (Macaca mulatta) of Indian genetic origin were housed in a biosafety level 2 NIAID facility and cared for in accordance with Guide for Care and Use of Laboratory Animals Report number NIH 82-53 (Department of Health and Human Services, Bethesda, 1985). All animal procedures and experiments were performed according to protocols approved by the IACUC of NIAID, NIH. The NHPs used in this study did not express the MHC class I Mamu-A*01, Mamu-B*08 and Mamu-B*17 alleles. NHPs were immunized subcutaneously in the medial inner forelegs and hind legs (total of 4 sites per animal) with 200 μg of the indicated SOSIP-mi3 adjuvated in SMNP (375 U/animal) as described (*66*). Immunizations and blood samples were obtained from naïve and immunized macaques at the time points indicated in Figure 5A.

### Flow Cytometry and cell sorting

Fresh bone marrow was flushed out of 1 femur and 1 tibia per mouse. Fresh mouse spleens were forced through a 70 µm mesh into FACS buffer (PBS containing 2% heat-inactivated FBS and 2 mM EDTA), and red blood cells of fresh spleens or bone marrow were lysed in ammonium-chloride-potassium buffer lysing buffer (Gibco) for 3 min. Frozen cells were thawed in a 37 °C water bath and immediately transferred to prewarmed mouse B cell medium consisting of RPMI-1640, supplemented with 10% heat-inactivated FBS, 10 mM HEPES, 1× antibiotic-antimycotic, 1 mM sodium pyruvate, 2 mM L-glutamine, and 53 µM 2-mercaptoethanol (all from Gibco).

Bait proteins were randomly conjugated to biotin and free biotin removed using EZ-Link Micro NHS-PEG_4_-Biotinylation Kit (ThermoFisher # 21955) according to the manufacturer’s instructions. Fluorophore conjugated bait and bait-KO antigen tetramers were prepared by mixing a 5 µg/mL solution of a single randomly-biotinylated bait protein with fluorophore-conjugated streptavidin (Table S7) at a 1:200 to 1:600 dilution in PBS for 30 min on ice. Conjugates were then mixed equivolumetrically.

RAMOS cells were harvested, washed in FACS buffer and stained with human FC-blocking reagent, biotinylated bait antigen-streptavidin tetramers (PE, AF647 and sometimes PECy7) and Zombie-NIR Live/Dead cell marker for 15 min before addition of anti-human antibodies to IgL-APC, IgK-BV421, IgM-FITC, and for some experiments, CD19-PECy7 (Table S7).

Mouse cells and controls (see below) were washed and resuspended in a solution of mouse Fc-receptor blocking antibody, fluorophore-conjugated antigen tetramers and Zombie-NIR Live/Dead cell marker for 15 min on ice. A mastermix of other antibodies was then added and cells stained for another 20 min on ice. Antibodies and reagents are listed in Table S7. All cells were analyzed on an LSRFortessa or cells were sorted on a FACS Aria III (both Becton Dickinson) using IOMA-expressing RAMOS cells as an antigen-binding positive control and splenocytes from naïve IOMAgl mice as negative controls (Figure S5). To derive absolute cell numbers, a master mix of AccuCheck counting beads (ThermoFisher #PCB100) in FACS buffer was prepared and 10^4^ beads/sample were added before acquisition. Absolute numbers of cells were calculated as:

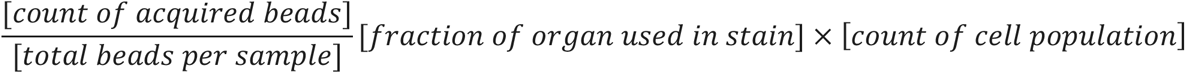

IOMA-expressing RAMOS cells were separated from unedited cells by sorting into RAMOS medium, and then washed and cultured as described above.

1838 single, mouse B cells from spleen and mesenteric lymph nodes of 3 IOMA iGL knock-in mice (ES30, HP1, and HP3) following the final boost (Week 18 or 23) were sorted into individual wells of a 96-well plate containing 5 μL of lysis buffer (TCL buffer (Qiagen, 1031576) with 1% of 2-β-mercaptoethanol). Plates were immediately frozen on dry ice and stored at −80 °C. Singlet, live Zombie-NIR^−^ CD4^−^ CD8^−^ F4/80^−^ NK1.1^−^ CD11b^−^ CD11c^−^ B220^+^ double Bait^+^ BaitKO^−^ lymphocytes were sorted unless GC B cells were sorted, which were gated as single, live Zombie-NIR^−^ CD4^−^ CD8^−^ F4/80^−^ NK1.1^−^ CD11b^−^ CD11c^−^ B220^+^CD38^−^ FAS^+^ lymphocytes (see Figure S5).

Mouse GC B cells for 10x Genomics single cell analysis were processed in PBS with 0.5% BSA instead of FACS buffer and 31,450 cells sorted into 5 µL of 0.05% BSA in PBS. Cells were spun down 400 g 6 min at 4 °C and volume adjusted to 22 µL before further processing.

### 10x Genomics single cell processing and next generation V(D)J sequencing

Cells were counted in the final injection volume, and 18,000 cells loaded onto a Chromium Controller (10x Genomics). Single-cell RNA-seq libraries were prepared using the Chromium Single Cell 5 v2 Reagent Kit (PN-1000265) according to manufacturer’s protocol. Chromium Single Cell Mouse BCR Amplification Kit (PN-1000255) was used for VDJ cDNA amplification. After QC, 5’ expression and VDJ Libraries were pooled 1:1 and sequenced on an Illumina NOVAseq S1 flowcell at the Rockefeller University Genomics Core.

### Computational Analyses of V(D)J sequences derived from IOMAgl mice by next generation sequencing

The single-cell V(D)J assembly was carried out by Cell Ranger 6.0.1. A customized reference was created by adding the knocked-in IOMA iGL V(D)J genes to the mouse GRCm38 V(D)J reference so Cell Ranger could recognize and assemble the human/mouse chimera transcripts. Contigs associated with a valid cell barcode according to Cell Ranger were selected for downstream processing using seqtk version 1.3-r106 (https://github.com/lh3/seqtk).

IgBlast standalone version 1.14 (*97*) was used to annotate the immunoglobulin sequences based on a custom database with mouse and human V(D)J genes. Productive IG sequences with more than 20 reads of coverage and with any identified isotype were selected for downstream processing. Unexpectedly, although the IgBlast algorithm identified the V and J genes for 8010 LC sequences, it failed to annotate the CDR3, and consequently, the information regarding their functionality was missing. We extracted and submitted 7782 (97.15%) sequences corresponding to the knock-in LC to IMGT/V-QUEST (*98*), which successfully identified the CDR3 and provided the productivity information.

Cell barcodes associated with sequences coded by different V genes for either HC or LC were considered doublets and were subsequently removed from downstream analysis. HCs and LCs derived from the same cell were paired, and clones were assigned using our previously-described IgPipeline (*76, 77*) (https://github.com/stratust/igpipeline/tree/igpipeline2_timepoint_v2).

### Single cell antibody cloning

Sequencing and cloning of mouse monoclonal antibodies from single cell-sorted B cells were performed as described (*99*) with the following modifications. Briefly, single cell RNA in 96-well plates was purified using magnetic beads (RNAClean XP, Beckman Coulter, Cat # A63987). RNA was eluted from the magnetic beads with 11 μL of a solution containing 14.5 ng/μL of random primers (Invitrogen, Cat # 48190011), 0.5% of Igepal Ca-630 (type NP-40, 10% in dH_2_O, MP Biomedicals, Cat # 198596) and 0.6 U/μL of RNase inhibitor (Promega, Cat# N2615) in nuclease-free water (Qiagen, Cat # 129117), and incubated at 65 °C for 3 min. cDNA was synthesized by reverse transcription (SuperScript™ III Reverse Transcriptase 10,000 U, Invitrogen, Cat# 18080-044). cDNA was stored at −80 °C or used for antibody gene amplification by nested polymerase chain reaction (PCR) after addition of 10 μL of nuclease-free water.

Mouse antibody genes were amplified using HotstarTaq DNA polymerase (Qiagen Cat # 203209) with the primer sets specific for the *Igh^IOMAiGL^* and *Igk^IOMAiGL^* transgenes. Primer sequences and reaction mixes are provided in Table S8. Thermocycler conditions were as follows for annealing (°C)/elongation (s)/number of cycles: PCR1 (IgG, IgM and IgK): 51/55/50; PCR2 (IgG and IgM): 54/55/50; PCR2 (IgK): 50/55/50.

PCR products of antibody HC and LC genes were purified and Sanger-sequenced (Genewiz) and *ab1 files analyzed using our previously described IgPipeline (https://github.com/stratust/igpipeline/tree/igpipeline2_timepoint_v2) (*76, 77*). V(D)J sequences were ordered as eBlocks (IDT) with short homologies for Gibson assembly and cloned into human IgG1 or human IgL2 expression vectors using the NEB Hifi DNA Assembly mix (NEB, Cat#E2621L). Plasmid sequences were verified by Sanger sequencing (Genewiz).

### Mutation analysis

All HC and LC V(D)J sequences were translated and the CDR3 region was trimmed. The resulting V region was aligned against the IOMA iGL and IOMA using MAFFT (*100*). Indels were ignored for downstream analysis. All mismatches to IOMA iGL were counted as total mismatches (Figure 3A). Only mismatches shared with IOMA mature when compared to IOMA iGL were used to assess chemical equivalence and calculate IOMA-like mutations (Figure 3A). Chemical equivalence was as follows: Group 1: G/A/V/L/I; Group 2: S/T; Group 3: C/M; Group 4: D/N/E/Q; Group 5: R/K/H; Group 6: F/Y/W; Group 7: P. The baseline was calculated using extracted IGHV1-2, IGHJ5 (318, 769) and IGLV2-23, IGLJ2 (1,790,961) sequences from healthy, HIV-negative donors generated by Soto et al. (*61*) and downloaded from cAb-Rep (*64*), a database of human shared BCR clonotypes available at https://cab-rep.c2b2.columbia.edu/.

3D neutralization plot shows the total number of V(D)J amino acid mutations (untrimmed) of each antibody vs the number of these mutations that are chemically equivalent to IOMA (Figure 3C). Chemical equivalence defined as above.

### *In vitro* neutralization assays

Pseudovirus neutralization assays were conducted as described (*62, 101*), either in house (Figure 2G, Figure 3A, Figure 4F, Figure 5B) or at the Collaboration for AIDS Vaccine Discovery (CAVD) core neutralization facility (Figure S1C). Monoclonal antibody IgGs were evaluated in duplicate with an 8-point, 3-fold dilution series starting at a top concentration of ∼100 µg/mL. All pseudovirus assays using monoclonal antibody IgGs were repeated at least twice for each value reported here. For polyclonal neutralizations, serum samples were heat inactivated at 56 °C for 30 min before being added to the neutralization assays, and then neutralization was evaluated in duplicate with an 8-point, 4-fold dilution series starting at a dilution of 1:60. The percent of neutralization at a 1:100 dilution (% 1:100) are reported for all serum samples. Tiers for viral strains were obtained from ref. (*102*) Antibody neutralization score was calculated as

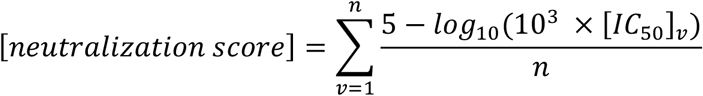

were *n* is number of different HIV pseudoviruses *v* tested for that antibody and [*IC*_50_]*_v_* is the IC_50_ of pseudovirus *v* in µg/mL.

### Statistical analysis

Comparisons between groups for ELISAs and neutralization assays were calculated using an unpaired or paired t-test in Prism 9.0 (Graphpad). Differences were considered significant when p values were less than 0.05. Exact p values are in the relevant figure at the top of the plot, with asterisks denoting level of significance (* denotes 0.01 < p ≤ 0.05, ** denotes 0.001 < p ≤ 0.01, *** denotes 0.0001 < p ≤ 0.001, and **** denotes p ≤ 0.0001). Comparisons between total amino acid mutations and IOMA-like mutations in antibodies cloned from IOMA iGL mice (Figure 3) were performed using a Pearson correlation and R^2^ values are presented.

### Analysis Software

Unless stated otherwise, Geneious Prime 2021.2.2, MacVector 18.2.0 and DNAStar SeqMan Pro 17.1.1 were used for sequence analysis and graphs were created using R language. Flow cytometry data were processed using Mac versions of FlowJo 10.7.2. and GraphPad Prism 9.3 and Microsoft Excel for Mac 16.54 were used for data analysis. Structural figures were made using PyMOL (Schrödinger, LLC) or ChimeraX (*103*). V(D)J gene assignments of NHP and murine antibodies were done using IMGT/V-QUEST (*98*). Sequence alignments were done using Clustal Omega (*104*).

---

**Figure S1.**
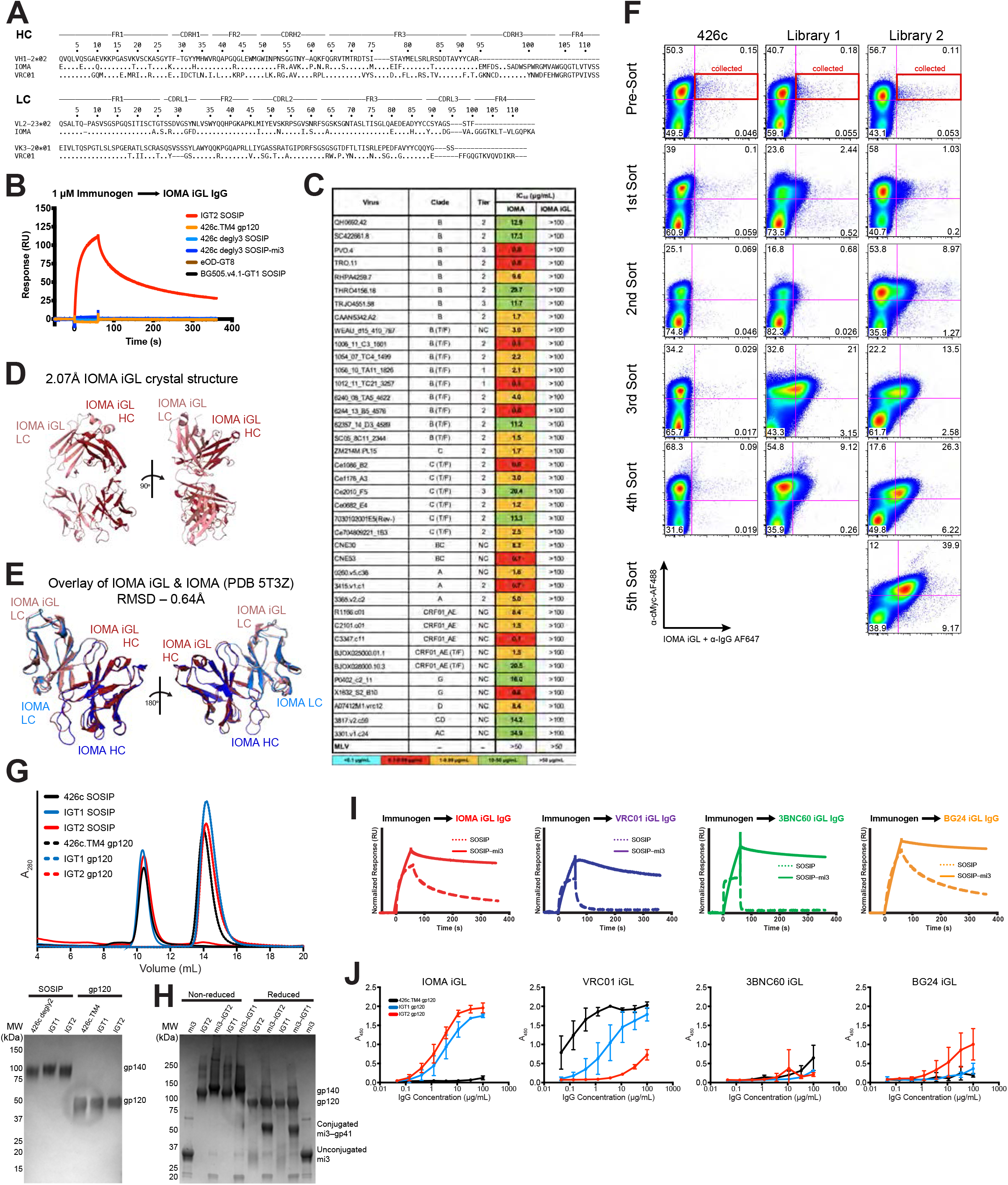
Development and characterization of IGT1 and IGT2 immunogens. **(A)** Amino acid alignment of IOMA and VRC01 to their respective germline V genes. **(B)** Representative SPR sensorgrams demonstrating no detectable binding of IOMA iGL to previously described immunogens (eOD-GT8, 426c.TM4, BG505.v4.1-GT1). This experiment was performed to qualitatively evaluate binding of IGT2 and previously described CD4bs immunogens to IOMA iGL rather than to derive affinity or kinetic constants. **(C)** Neutralization titers (IC_50_s) of IOMA and IOMA iGL against a panel of 38 viruses and an MLV control. **(D)** 2.07 Å crystal structure of IOMA iGL Fab shown in two views. **(E)** Structural overlay of IOMA iGL Fab and IOMA Fab from BG505-bound structure (PDB 5T3Z).**(F)** Flow cytometric analysis of yeast cells expressing 426c.TM4 starting protein (left), Library 1 (middle), or Library 2 (right) stained with IOMA iGL IgG/anti-IgG AF647 (x-axis) and anti-cMyc AF488 (y-axis). **(G)** Representative size exclusion chromatography profiles and Coomassie-stained SDS-PAGE analysis for 426c.TM4 gp120, IGT1 gp120, and IGT2 gp120, 426c SOSIP, IGT1 SOSIP, and IGT2 SOSIP demonstrating that all of these proteins are monodispersed samples and that the selected mutations do not alter the stability or behavior of the immunogens compared to the starting proteins. **(H)** Coomassie-stained SDS–PAGE analysis for mi3, IGT2, IGT2-mi3, IGT1, and IGT1-mi3 under non-reducing and reducing conditions. **(I)** SPR sensorgrams demonstrating binding of IGT2 (dashed line) and IGT2-mi3 (solid line) to IOMA iGL IgG (red), VRC01 iGL IgG (purple), 3BNC60 iGL IgG (green), and BG24 iGL IgG (orange). IgG was immobilized to the CM5 chip and 1 µM SOSIP or 1 µM SOSIP-mi3 was flowed over the chip surface. **(J)** Representative ELISA binding curves measuring binding of 426c.TM4 gp120, IGT1 gp120, and IGT2 gp120 to the same iGL IgGs as in (I). Dots indicate mean and error bars indicate 95% confidence interval.

**Figure S2.**
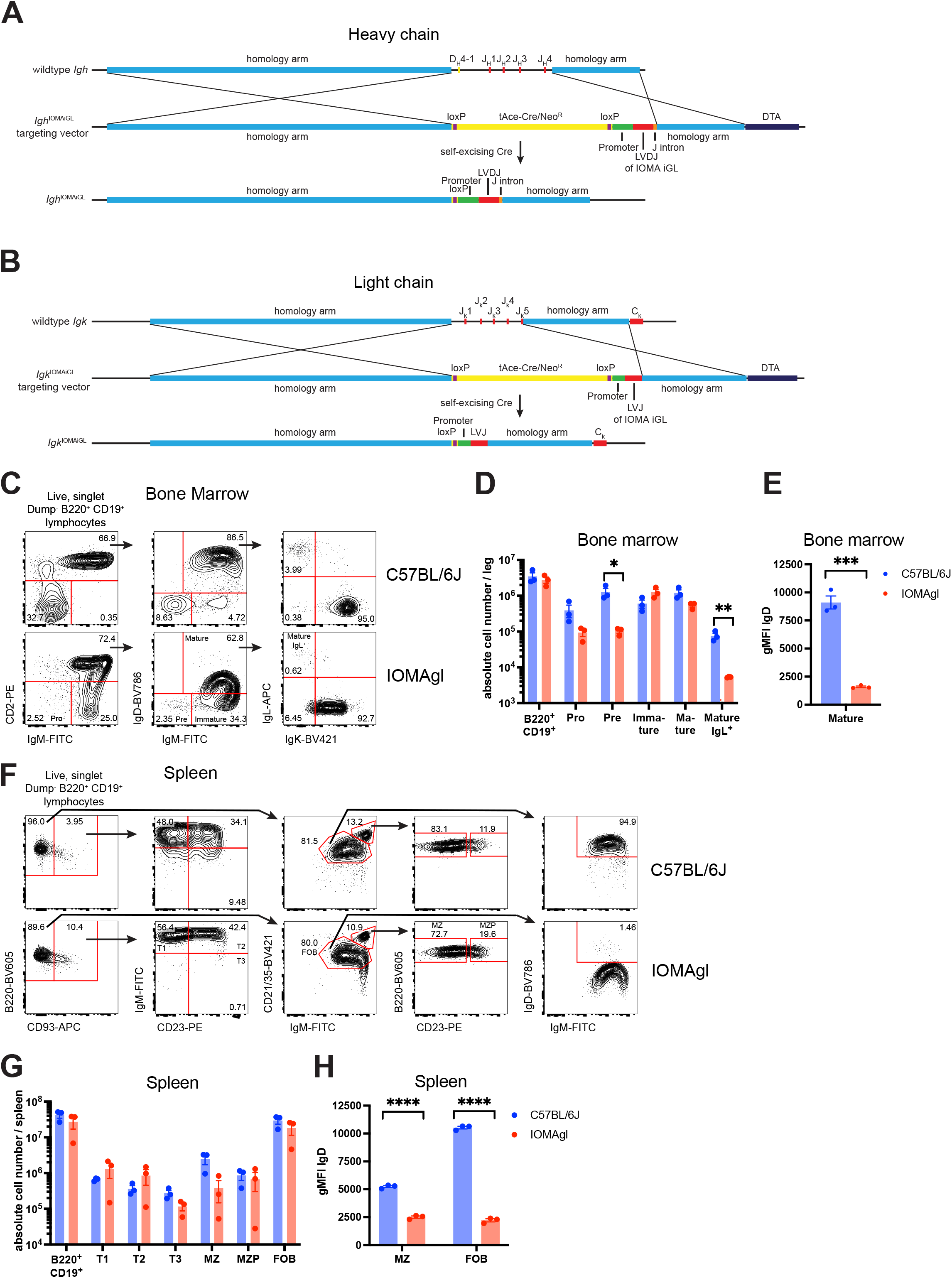
Targeting strategy and characterization of IOMAgl mice. **(A)** In *Igh^IOMAiGL^* mice *Ighd4-1* to *Ighj4* are replaced by a self-excising Neomycin cassette followed by the mouse *Ighv9-4* promoter, a leader sequence (L) followed by the iGL version of the IOMA HC VDJ sequence and a *Ighj1* splice donor sequence. **(B)** In *Igk^IOMAiGL^*mice *Igkj1* to *Igkj5* are replaced by a self-excising Neomycin cassette followed by a mouse *Igkv3-12* promoter, a leader sequence followed by the iGL version of the IOMA lambda LC VDJ sequence and a *Igkj5* splice donor sequence. DTA, diphtheria toxin A **(C)** Flow cytometric analysis of B cell development in the bone marrow of control (C57BL/6J) or IOMAgl (*Igh^IOMAiLG/IOMiGL^ Igk^IOMAiG/IOMAiGLL^*) mice. **(D)** Absolute cell number quantification from (C). **(E)** Geometric mean fluorescence intensity (gMFI) of IgD in mature recirculating B cells from the bone marrow. **(F)** Flow cytometric analysis of peripheral B cell development in the spleens of control (C57BL/6J) or IOMAgl mice. **(G)** Absolute cell number quantification from (F). **(H)** gMFI of IgD in marginal zone and follicular B cell. MZ, marginal zone B cells; MZP, marginal zone precursors; FOB, follicular B cells. Data from 1 of 2 independent experiments, each dot represents a data from 1 mouse. Bars represent mean ± SEM. Statistical analysis used unpaired t test.

**Figure S3.**
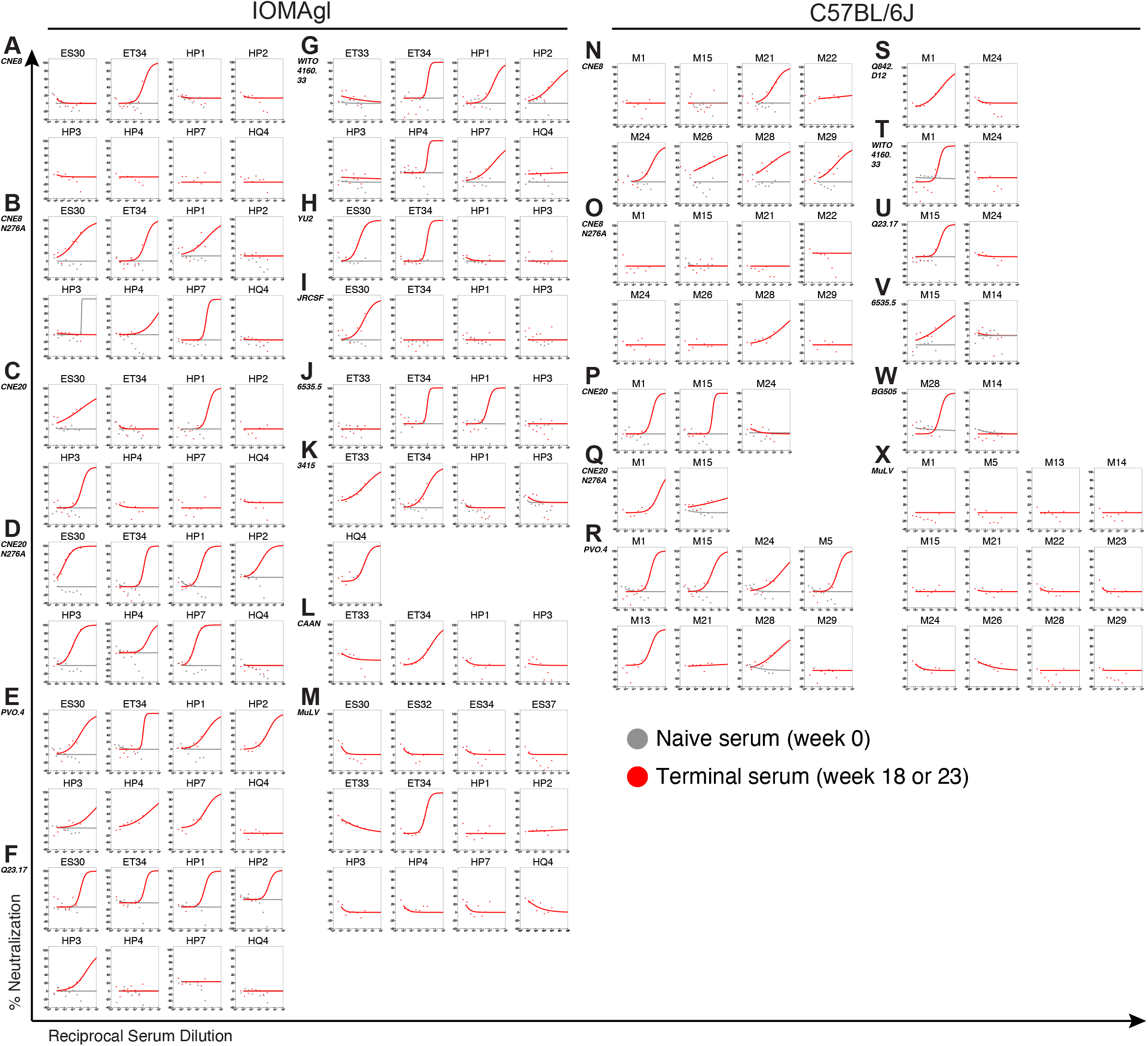
Serum neutralization from immunized mice. Neutralization curves of serum isolated from IOMA iGL transgenic mice (**A-M**) or C57BL/6J wildtype mice (**N-X**) against the following HIV strains or control MuLV: **(A,N)** CNE8, **(B,O)** CNE8 N276A, **(C,P)** CNE20, **(D,Q)** CNE20 N276A, **(E,R)** PVO.4, **(F,U)** Q23.17, **(G,T)** WITO4160.33, **(H)** YU2, **(I)** JRCSF, **(J, V)** 6535.5, **(K)** 3415_V1_C1, **(L)** CAAN5342.A2, **(M,X)** MuLV, (**S**) Q842.D12 and (**W**) BG505. Naïve serum was also tested against the same strains when available. Note that sera which showed neutralization activity of < 40% as listed in Table S3 are presented in Figure 2G as white rectangles; several of these sera neutralized strains above background including ET33 against PVO.4; ET34 against CNE20 N276A and Q23.17; HP1 against CNE8 N276A, CNE20, and WITO4160.33; HP2 against Q23.17; HP3 against Q23.17 and PVO.4; HP4 against CNE8 N276A, CNE20 N276A, and PVO.4.

**Figure S4.**
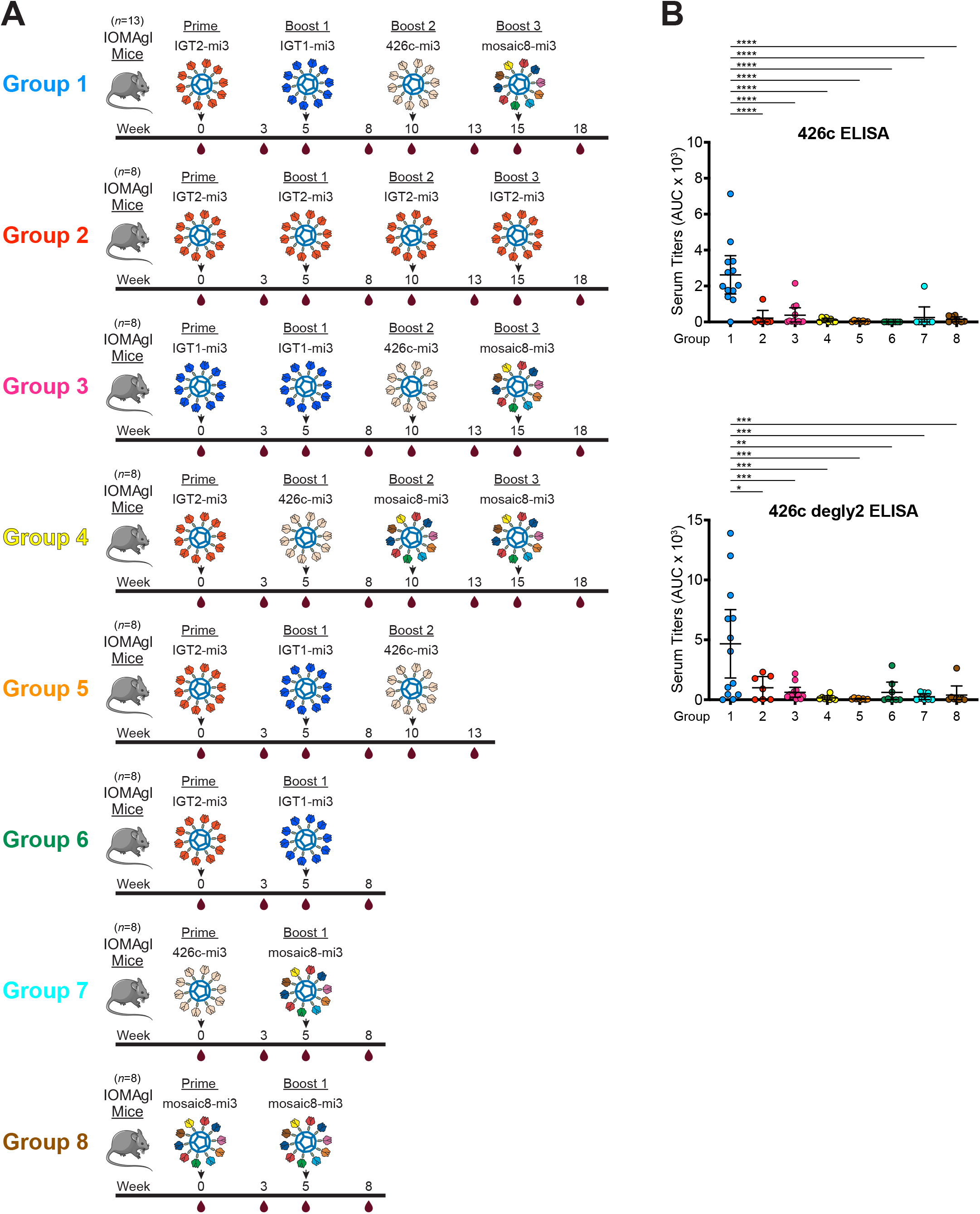
Screening immunization regimens to determine the optimal boosting strategy. **(A)** Schematic and timeline of immunization strategies to determine the optimal regimen to elicit IOMA-like bNAbs. **(B)** Serum ELISA binding to 426c and 426c degly2 represented as AUC using serum samples isolated from mice at the end of the regimen.

**Figure S5.**
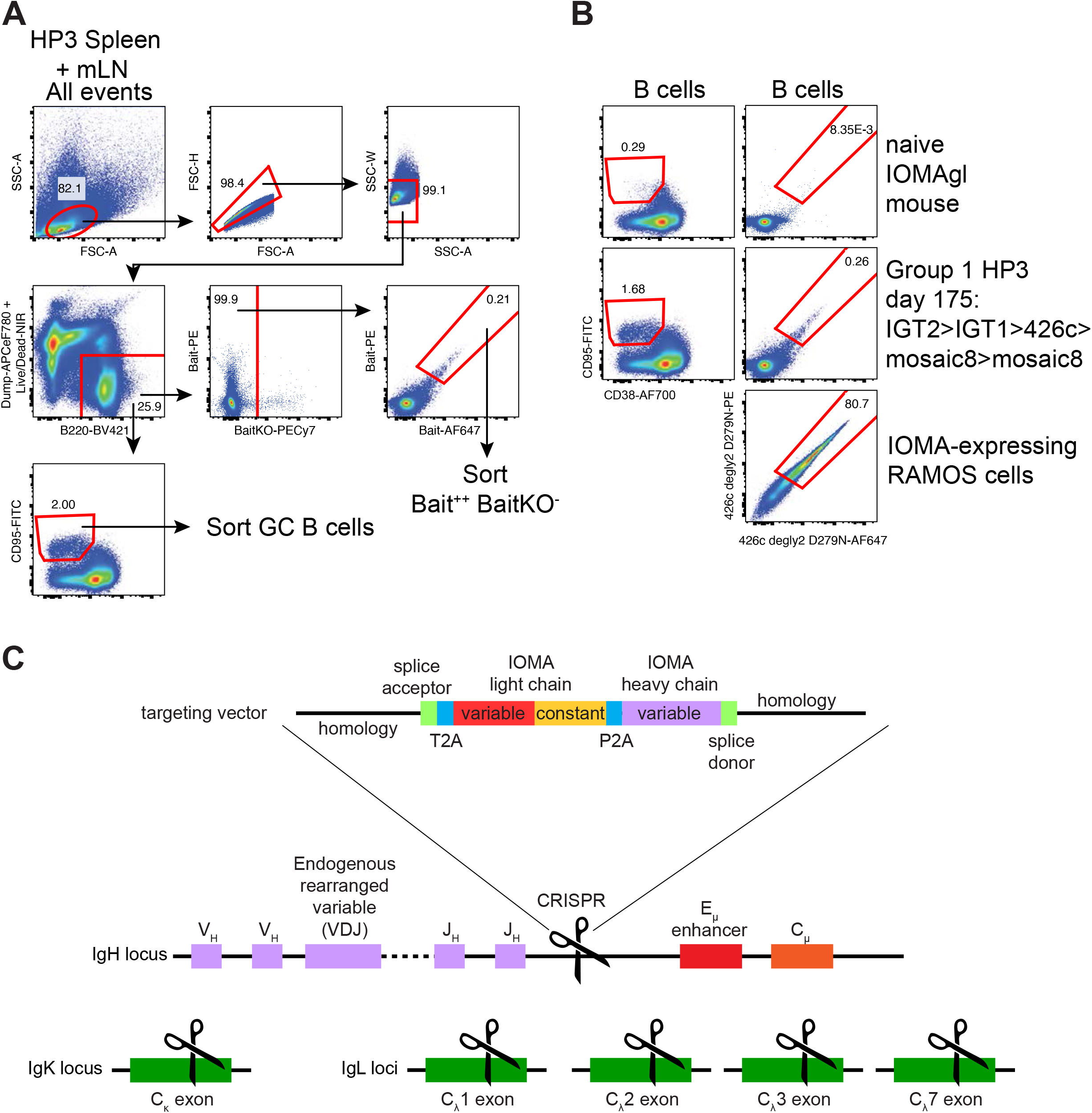
Cell sorting strategies and sorting controls. **(A)** Representative full gating of cell sorts for single cell Bait++ BaitKO^-^ B cell cloning and 10x Genomics next generation VDJ sequencing of bulk-sorted GC B cells from splenic and mesenteric lymph nodes. Baits used were 426c degly2 D279N or CNE8 N276A with 426c degly2 D279N-CD4bs KO, the former is shown. **(B)** Induction of germinal center response and wt SOSIP-binding cells by immunization regimen (group 1). Naïve IOMAgl mouse splenocytes and IOMA-expressing RAMOS cells served as negative and positive control, respectively. (**C**) Gene editing strategy to generate IOMA-expressing RAMOS cells. Simultaneous targeting of IgH, IgK and IgL loci with CRISPR/Cas9 to delete endogenous LCs and edit a promoterless tricistronic expression cassette into the IgH locus to express IOMA on the surface of RAMOS cells. A polycistronic mRNA was created using T2A and P2A sequences to induce ribosomal skipping (*92*).

**Figure S6.**
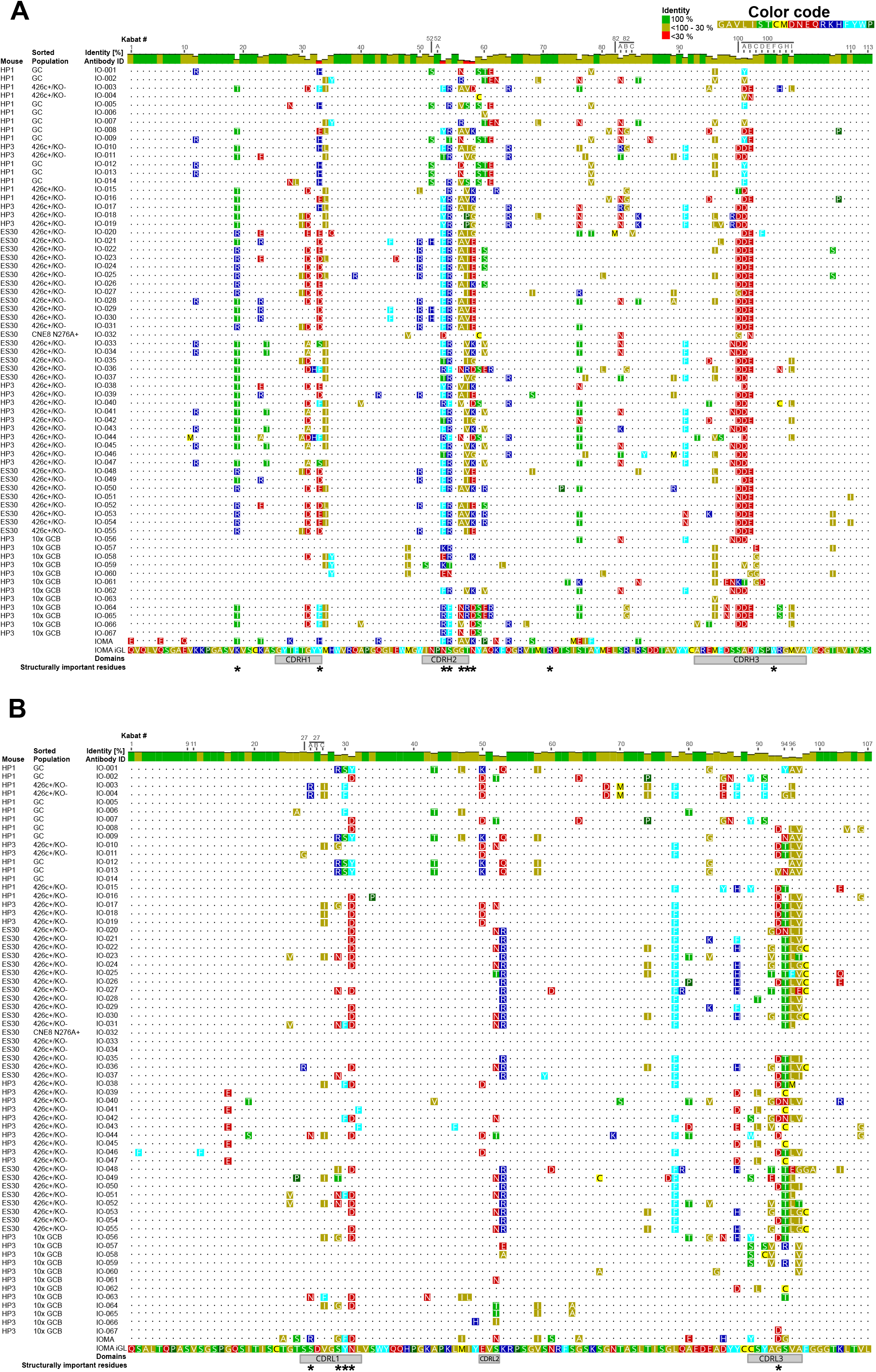
Amino acid alignments of selected IOMAgl mouse-derived antibodies. **(A)** V_H_ alignment of cloned antibodies IO-001 to IO-067 that were expressed and tested for Env binding. IOMA iGL and IOMA sequence at the bottom as reference. Mouse ID and population sorted are indicated. Differences to IOMA iGL are highlighted using chemically similar color coding; dots indicate identical residues to IOMA iGL. Kabat numbering and percent identity of residues are indicated on top. Domains and residues of structural importance are annotated below. **(B)** as above but corresponding V_L_ alignment.

**Figure S7.**
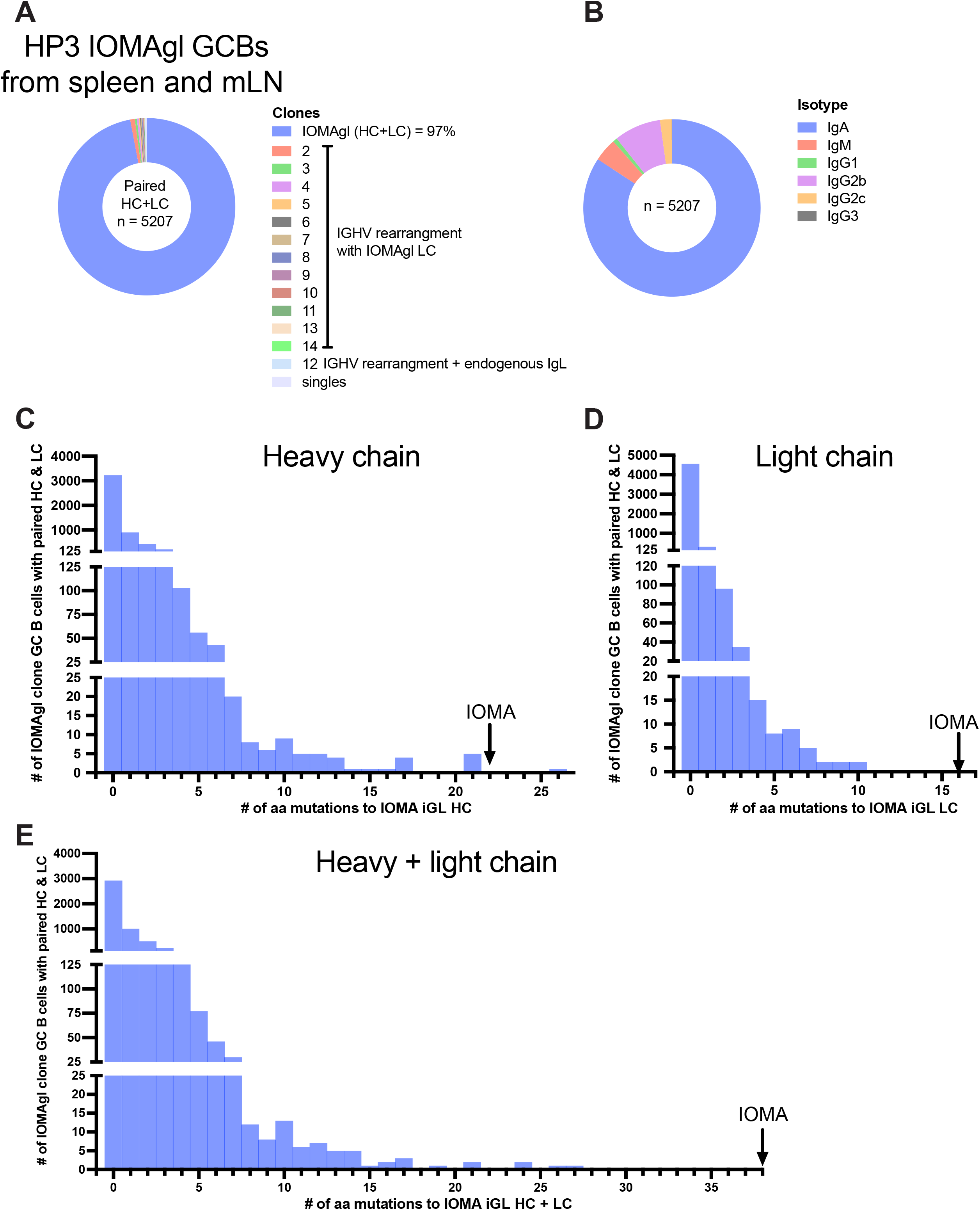
Next generation single cell VDJ analysis determines the extent of mutations in germinal centers of IOMAgl mice. **(A)** Clonal analysis of paired HC and LC sequences from splenic and mesenteric lymph node germinal center B cells of IOMAgl mouse HP3. **(B)** Isotype distribution among these cells. **(C)** Frequency distribution of the number of amino acid mutations to IOMA iGL in the HC sequences of these cells. **(D)** Frequency distribution of the number of amino acid mutations to IOMA iGL in the LC sequences of these cells. **(E)** Frequency distribution of the number of amino acid mutations to IOMA iGL in the paired HC and LC sequences of these cells.

**Figure S8.**
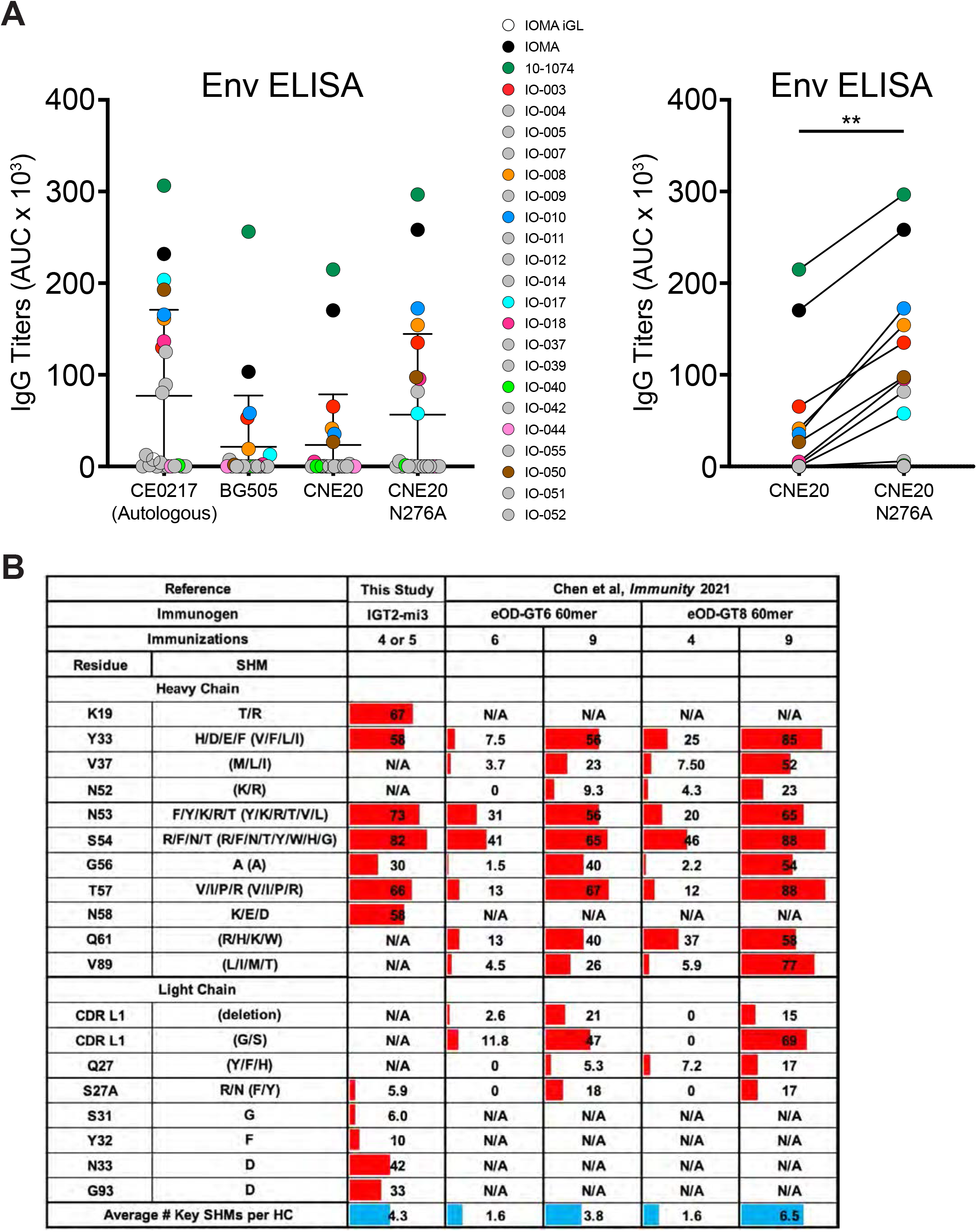
Monoclonal antibodies cloned from IOMA iGL transgenic mice bind to heterologous Envs. (A) AUC of. ELISA binding curves of selected monoclonal antibodies isolated from IOMA iGL knock-in mice to BG505, CE0217, CNE20 and CNE20 N276A SOSIPs. **(B)** Comparison of the occurrence frequency of key mutations among IOMA-like antibody sequences selected for cloning and VRC01-class antibody sequences from reference 37 at different time points throughout the respective sequential immunization regimen. Mutations essential for IOMA-class antibody binding to gp120 are listed first, while mutations essential for VRC01-class antibody binding to gp120 are listed second in brackets. Values for each residue represent the percentage of antibodies containing one of the essential mutations at that position.

**Table S1:**
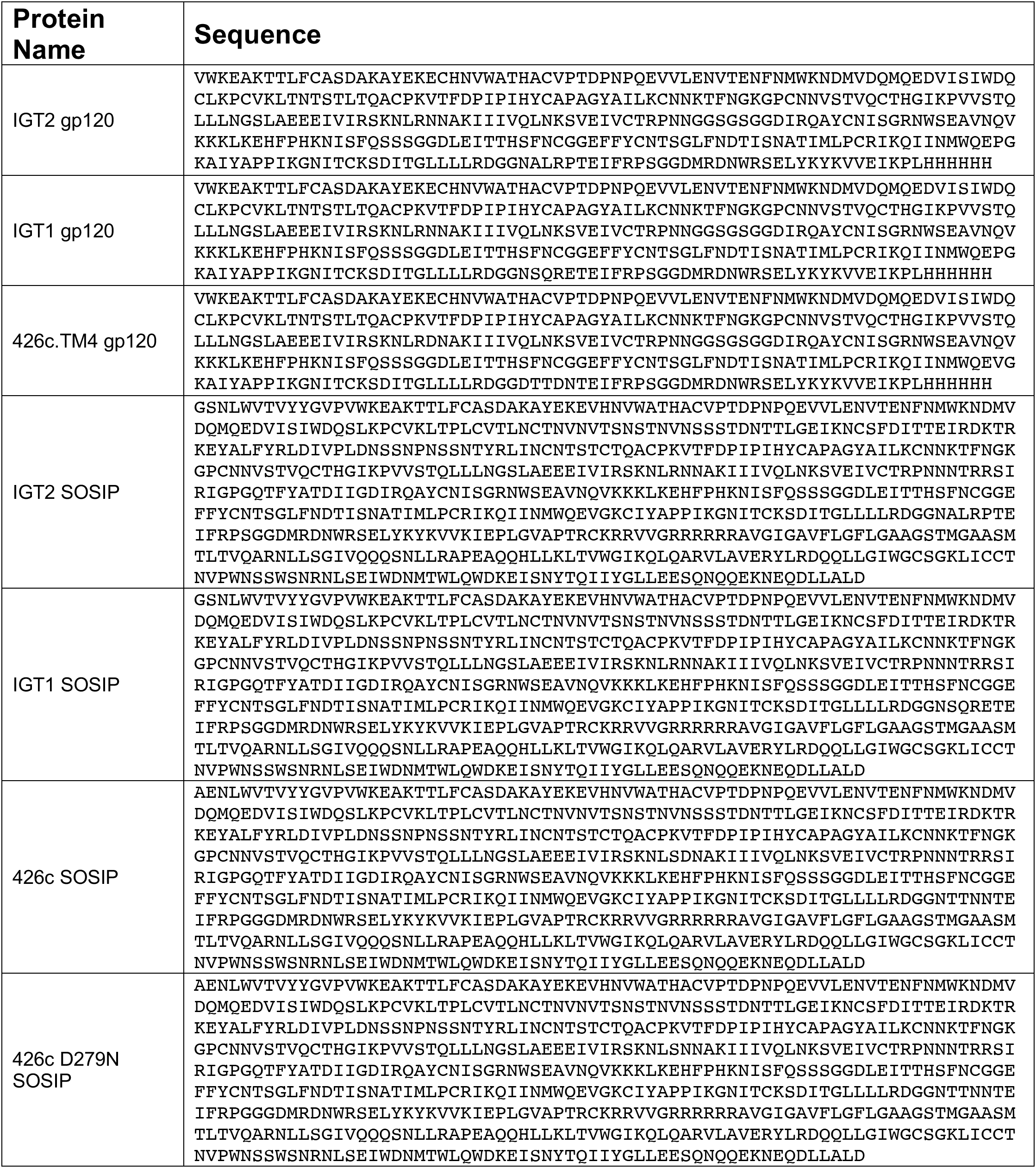
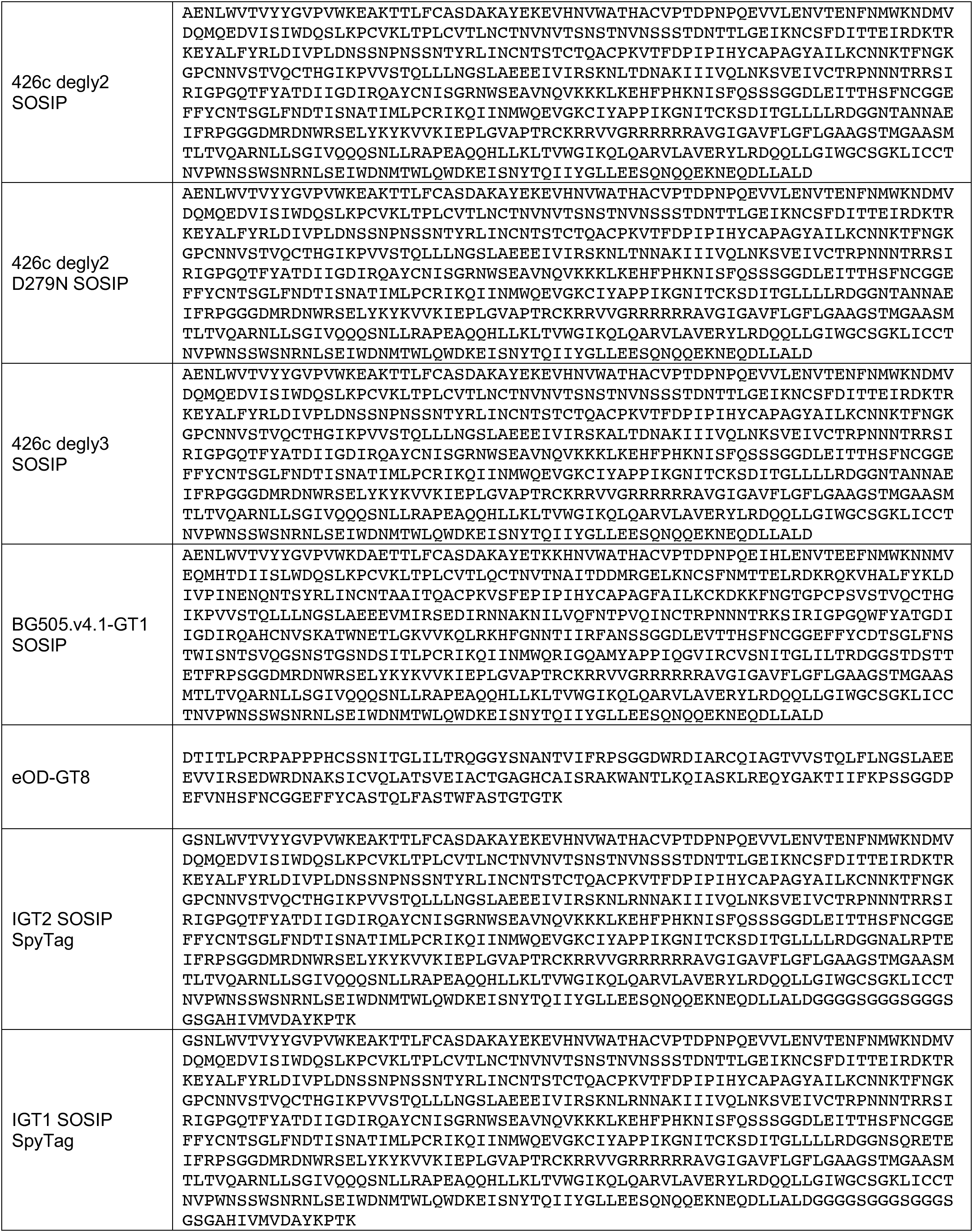
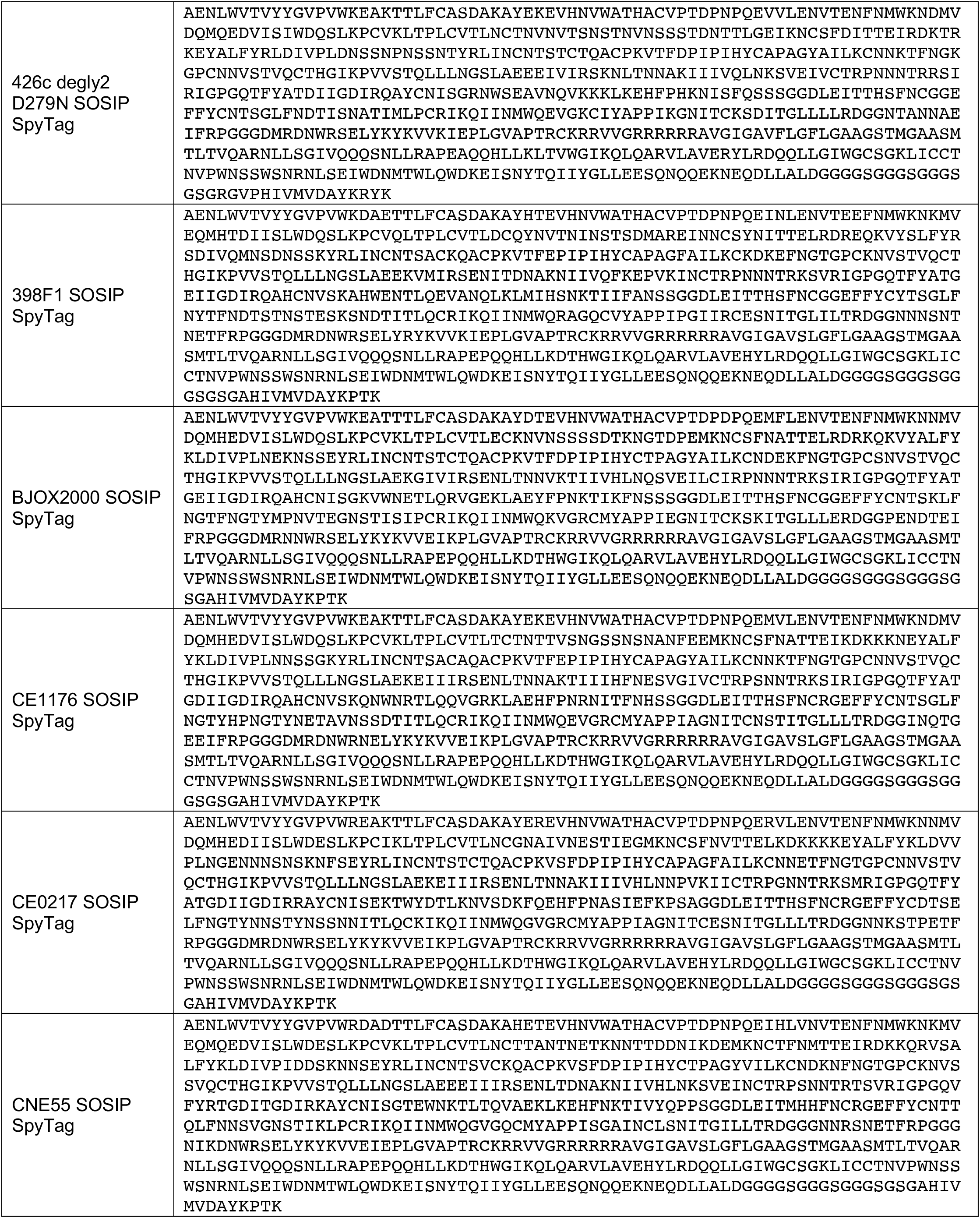
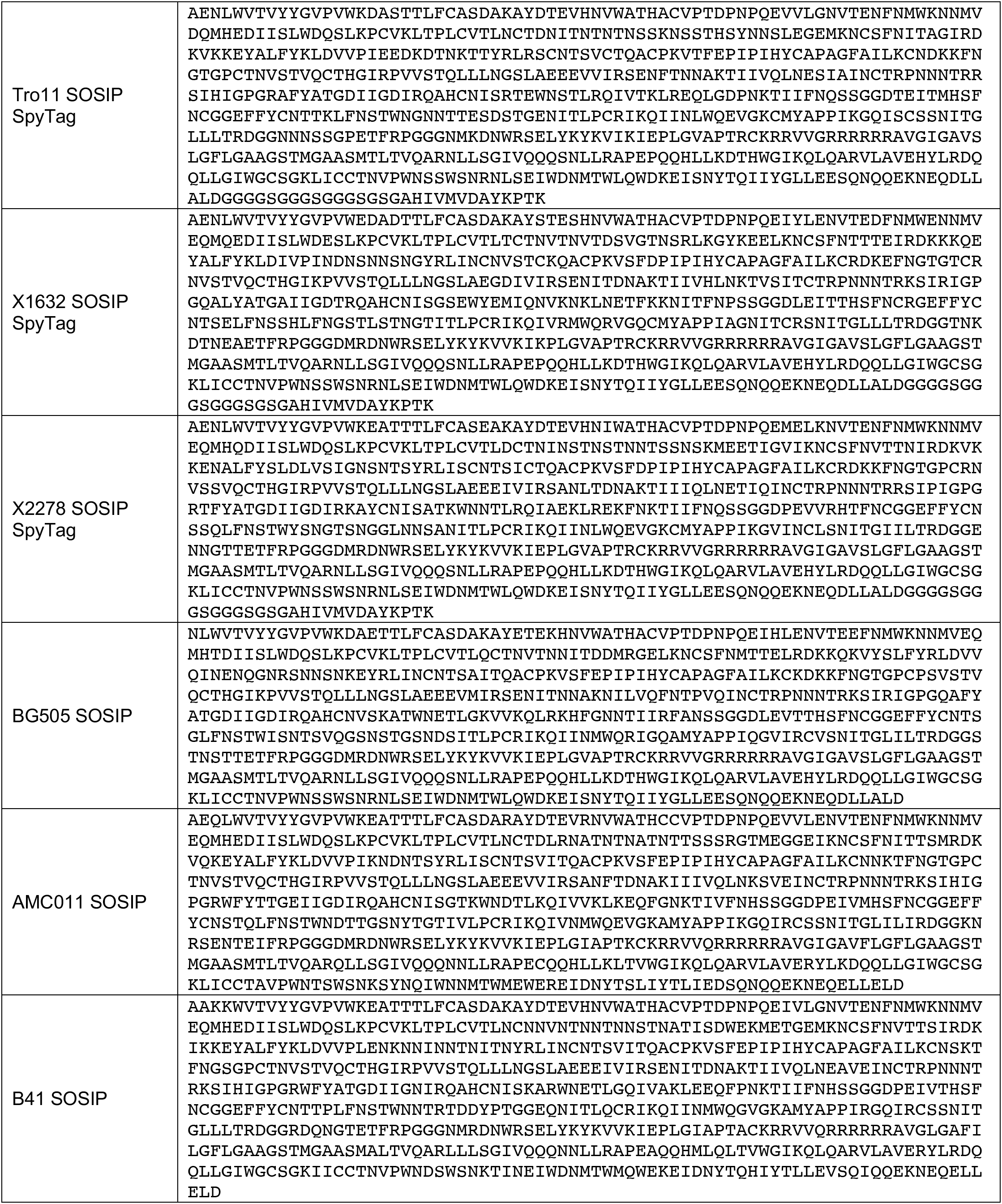
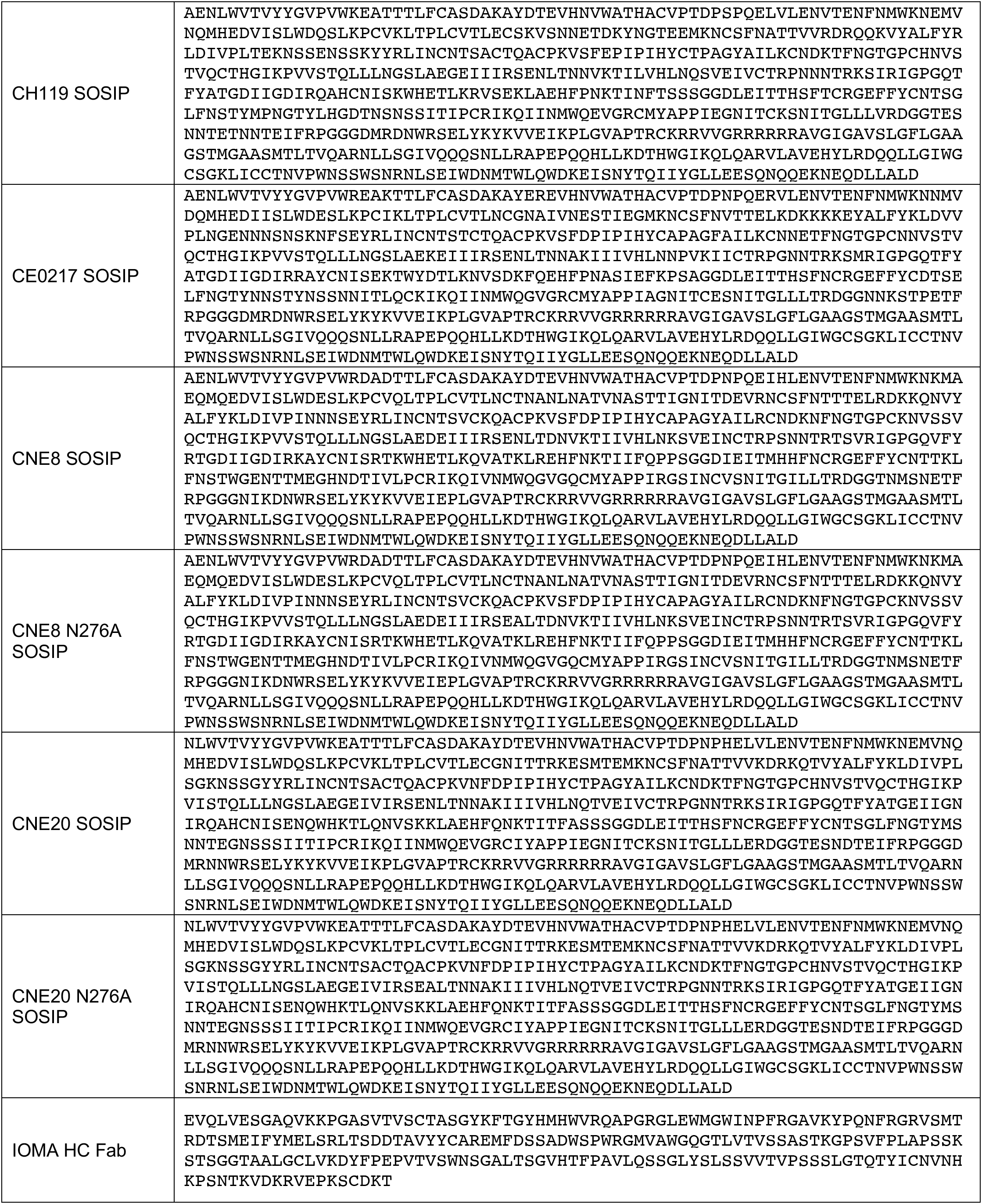
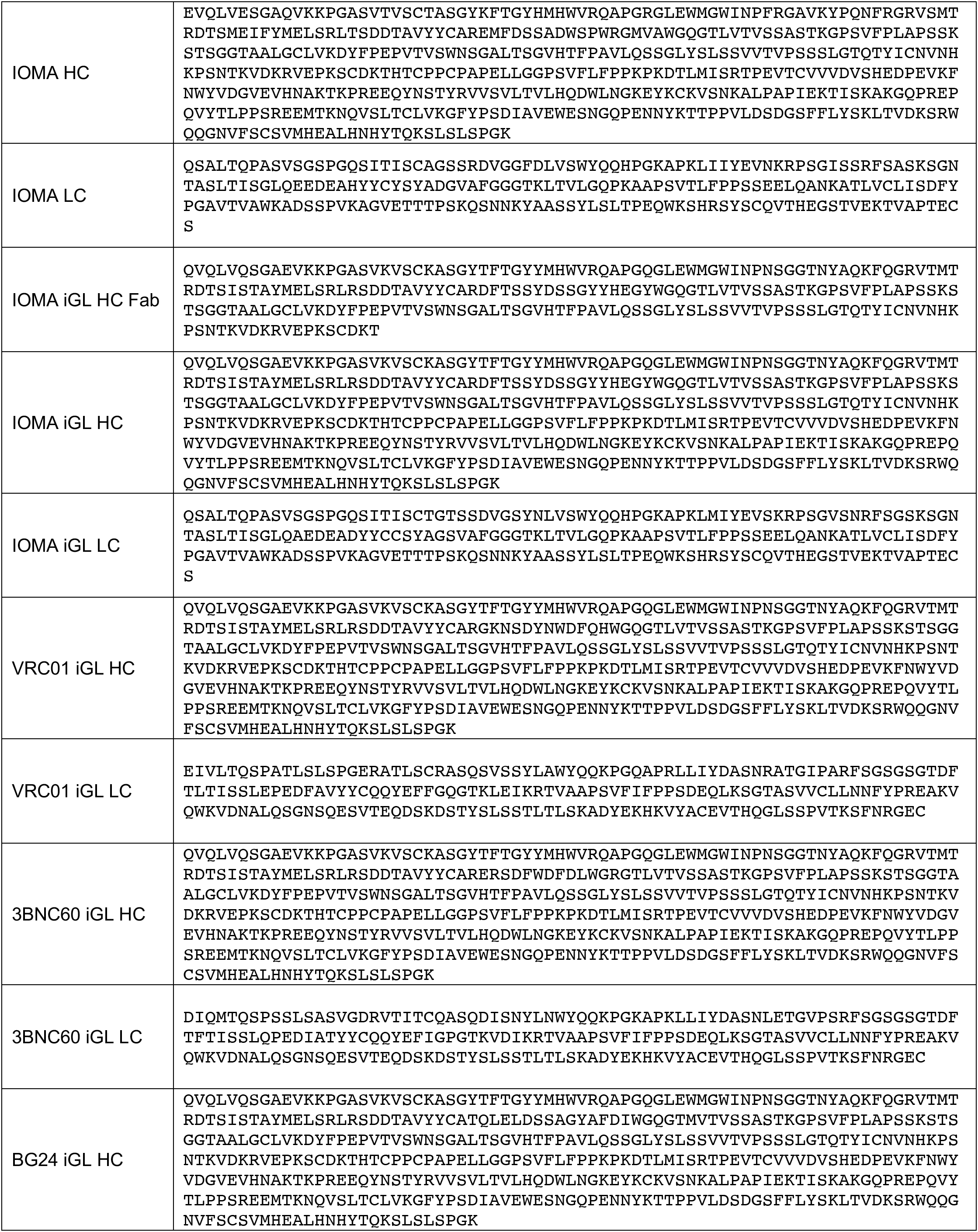
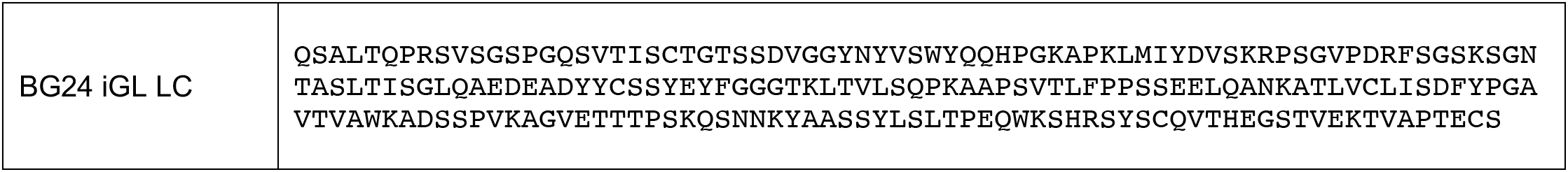
Amino acid sequences for HIV Envs and antibodies used in this study.

**Table S2:**
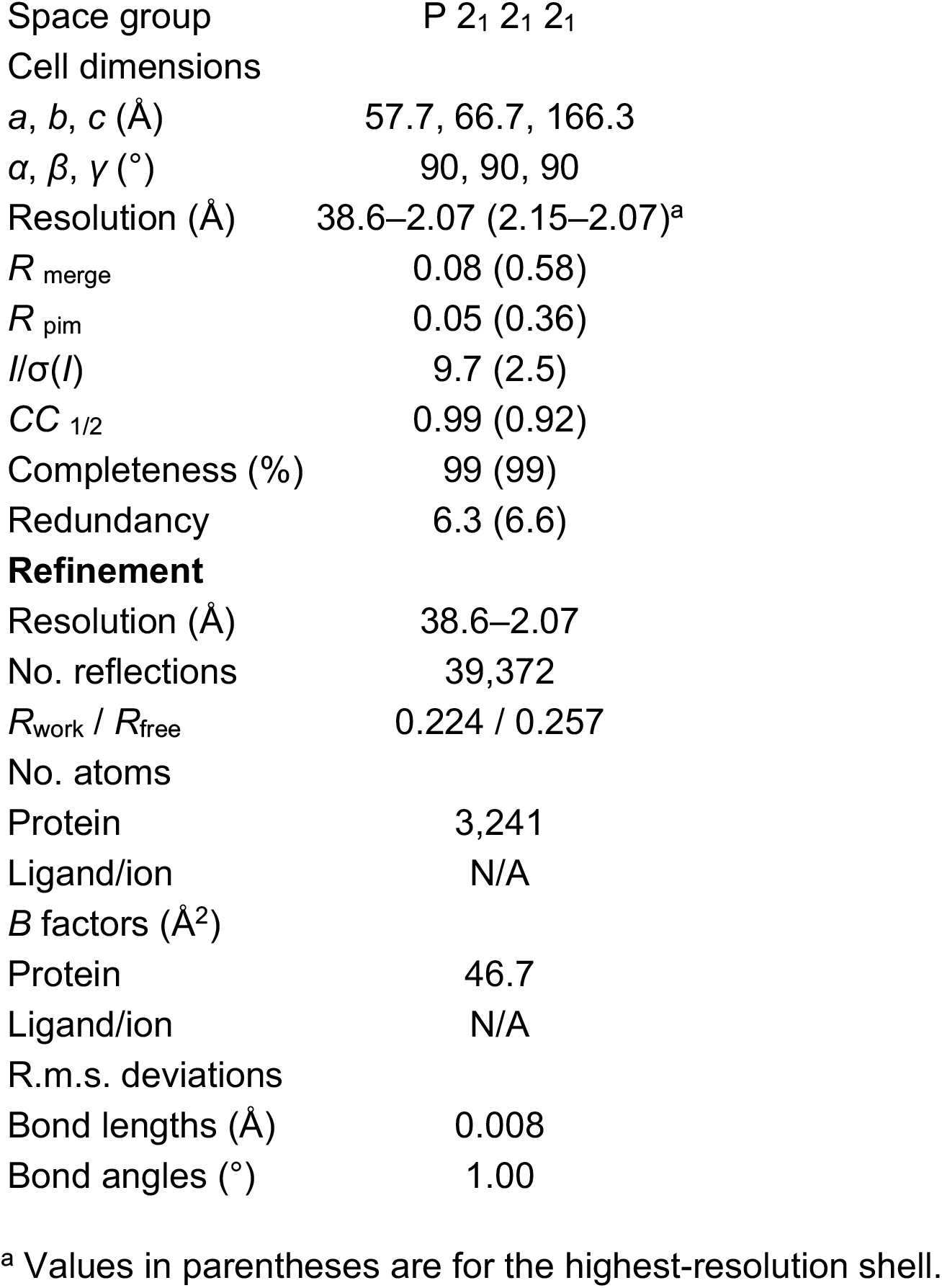
X-ray data collection for IOMA iGL Fab crystals.

**Table S3:**
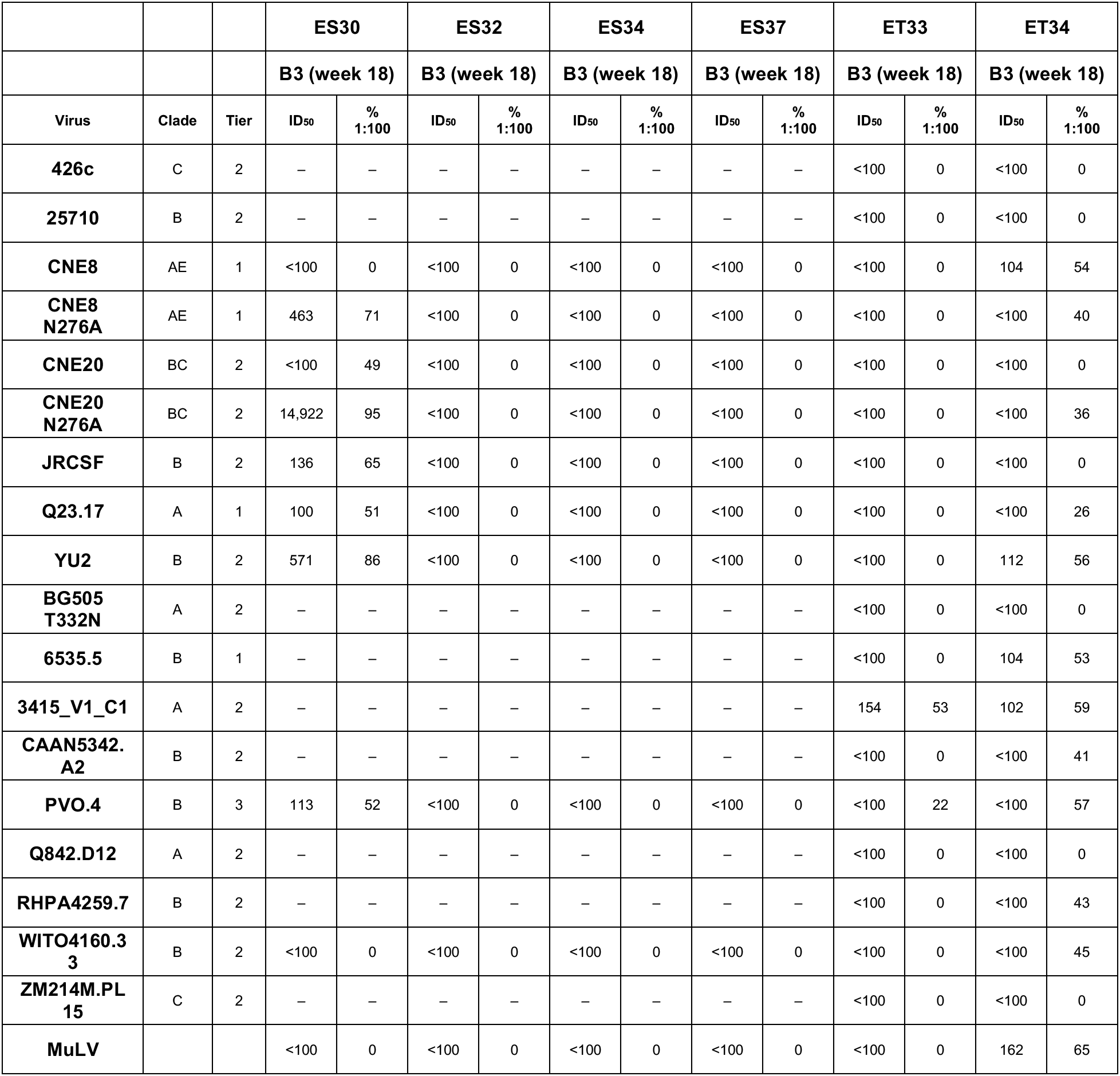
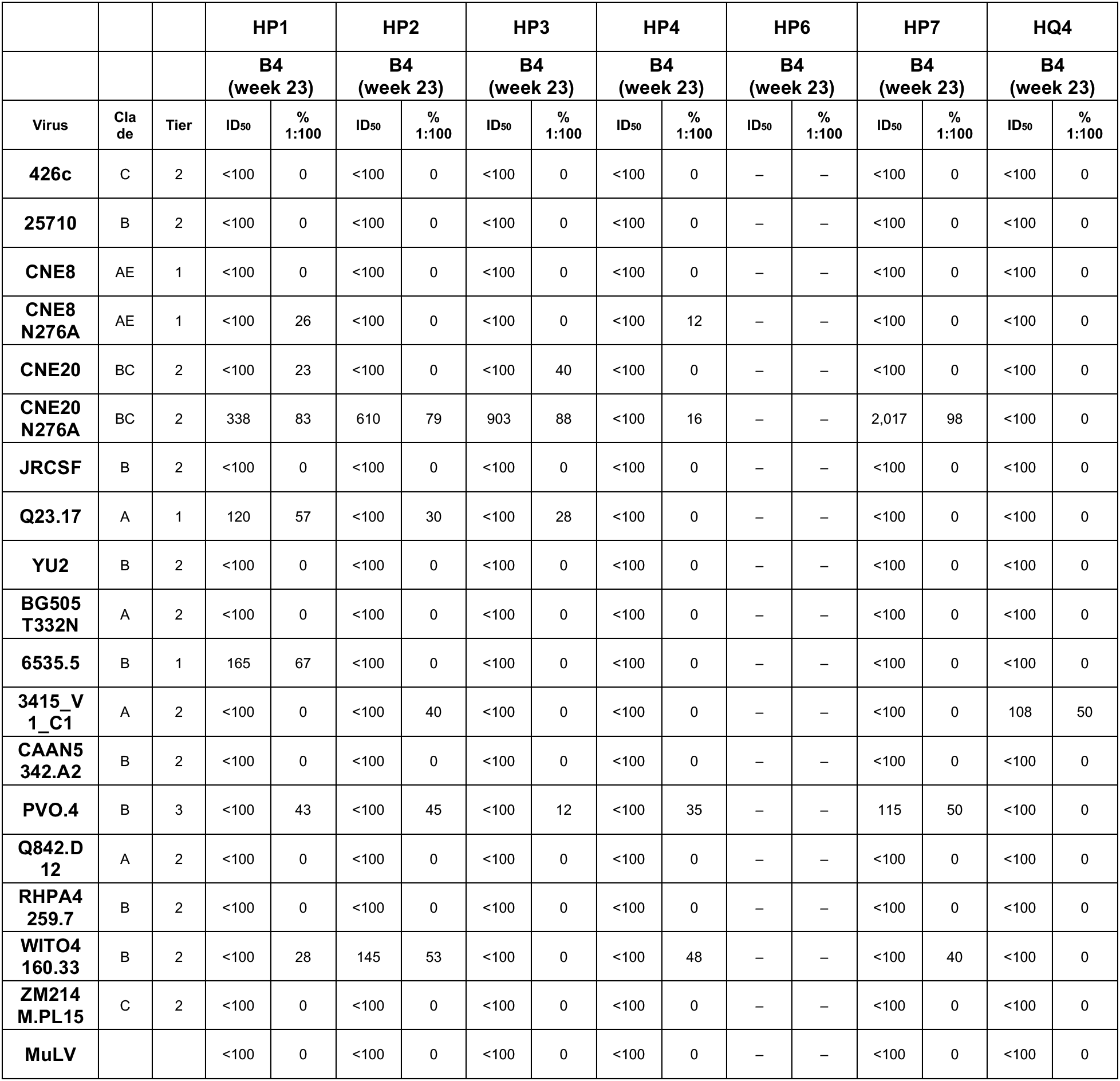
Serum neutralization data for IOMA iGL transgenic mice.

**Table S4:**
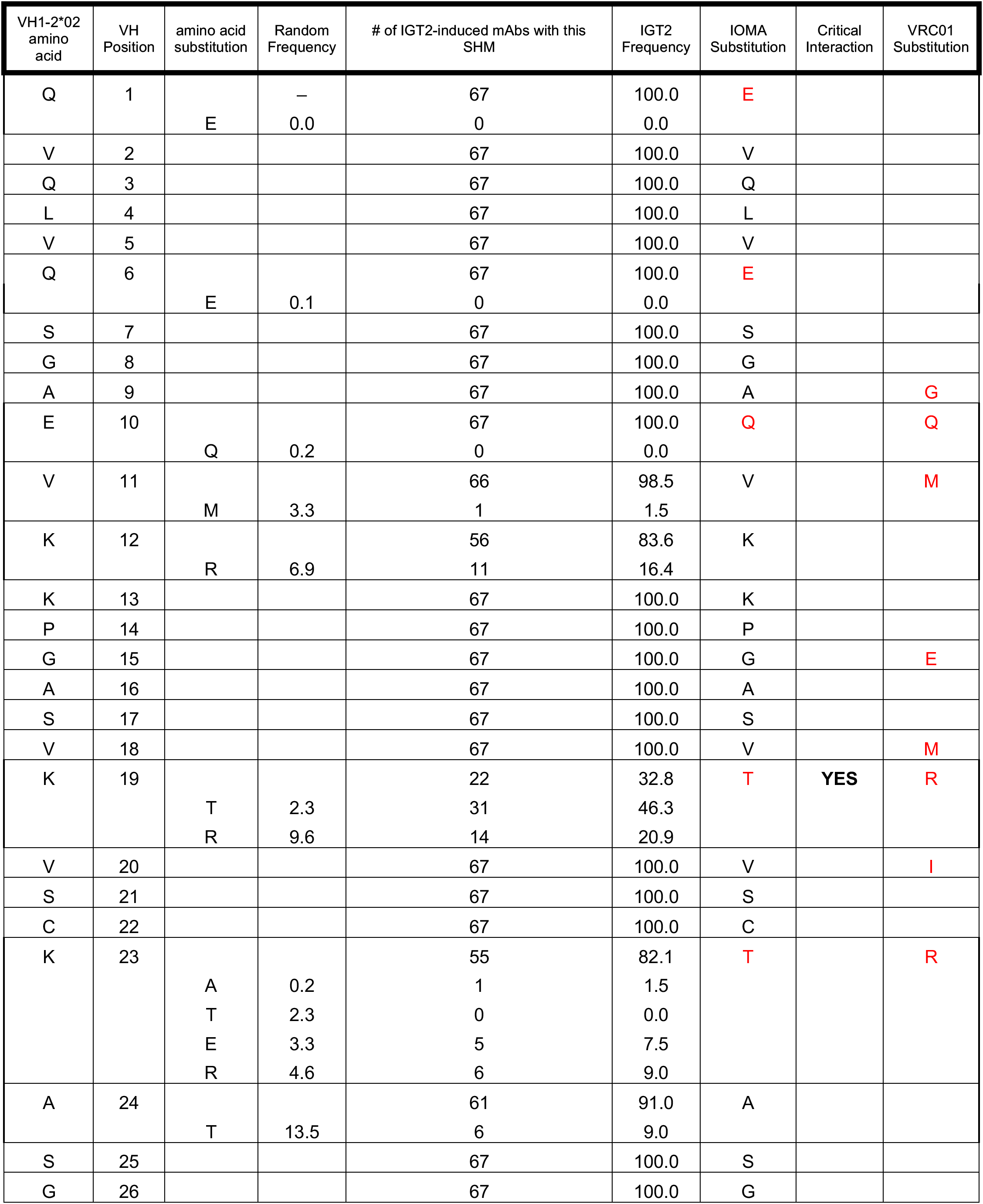
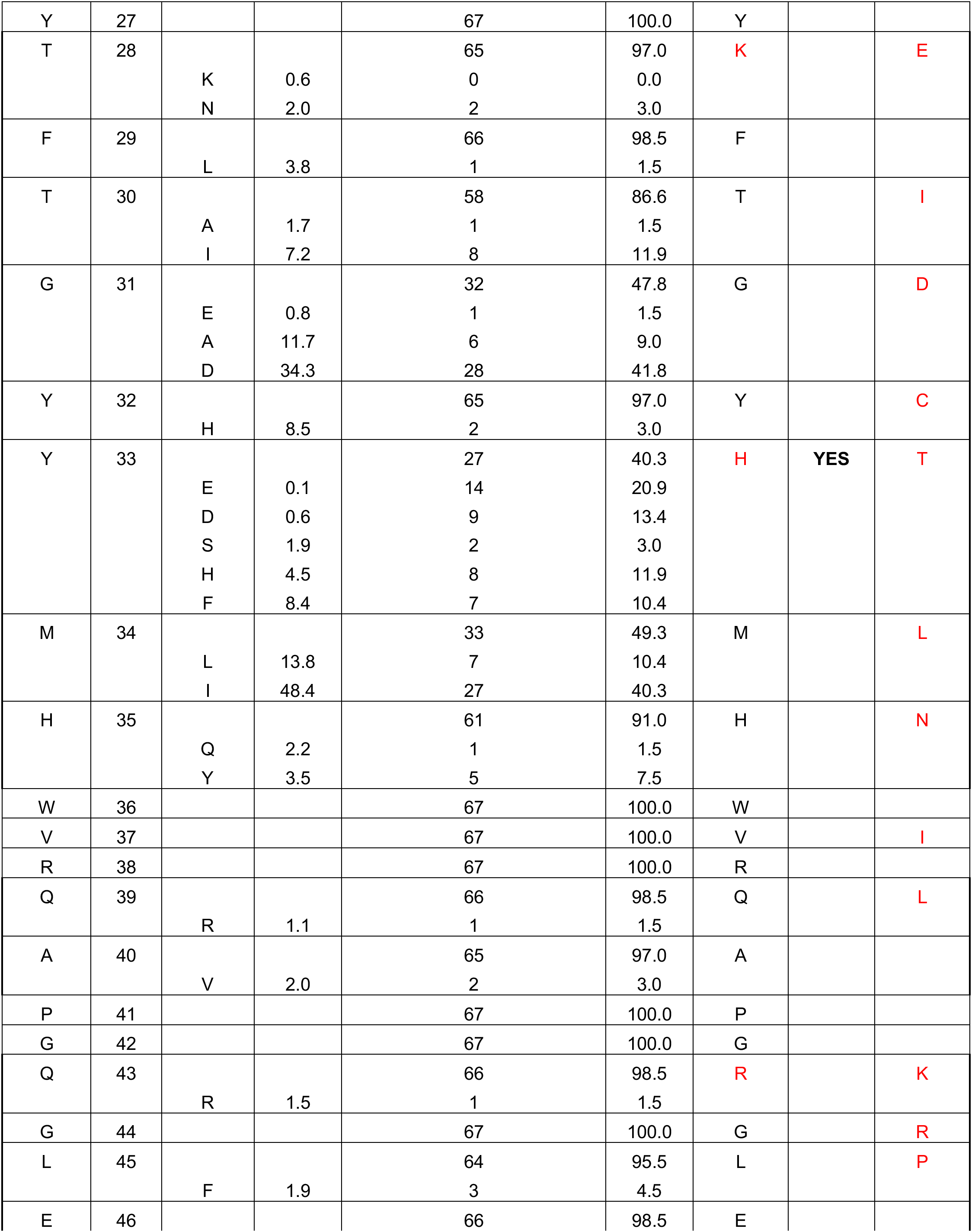
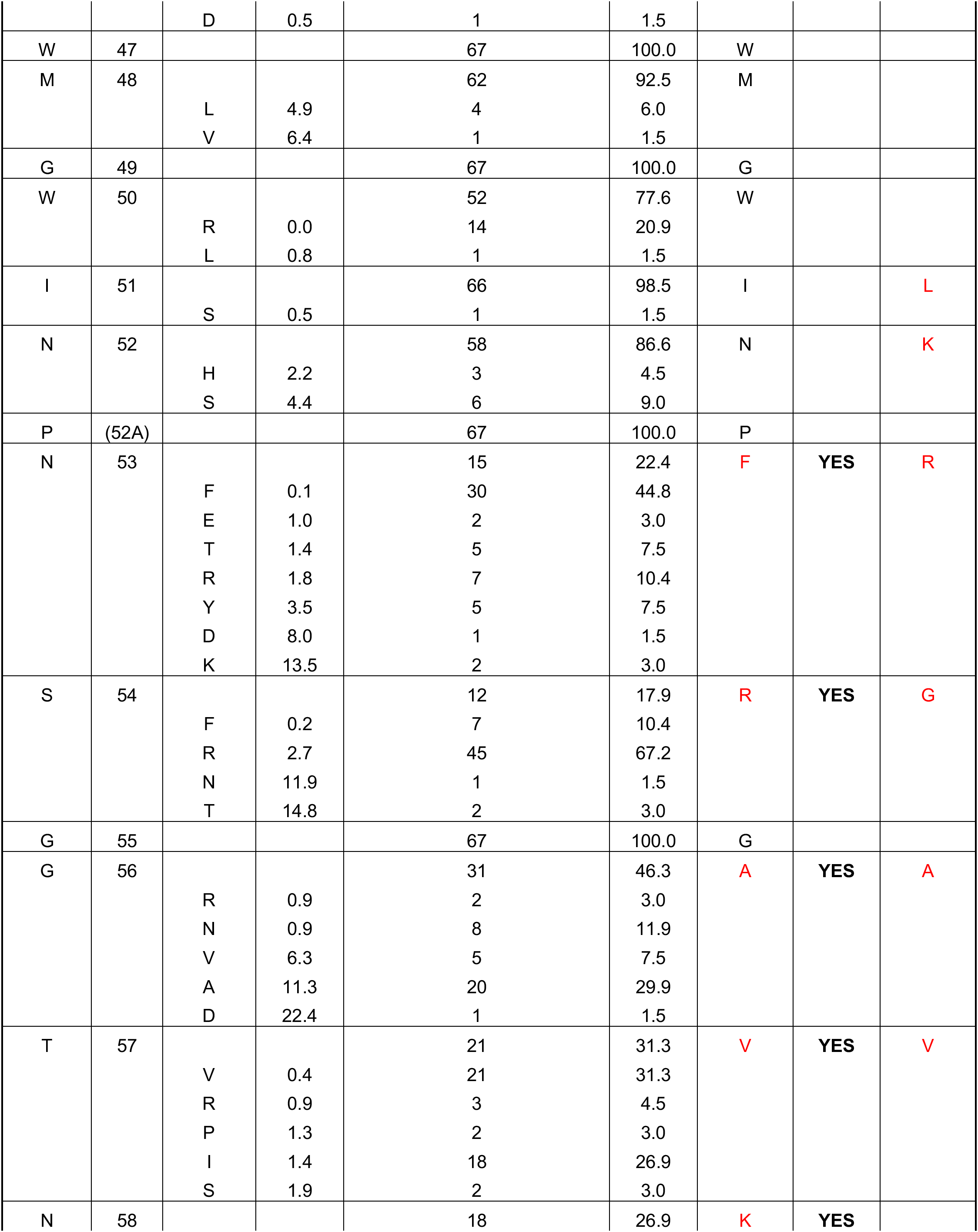
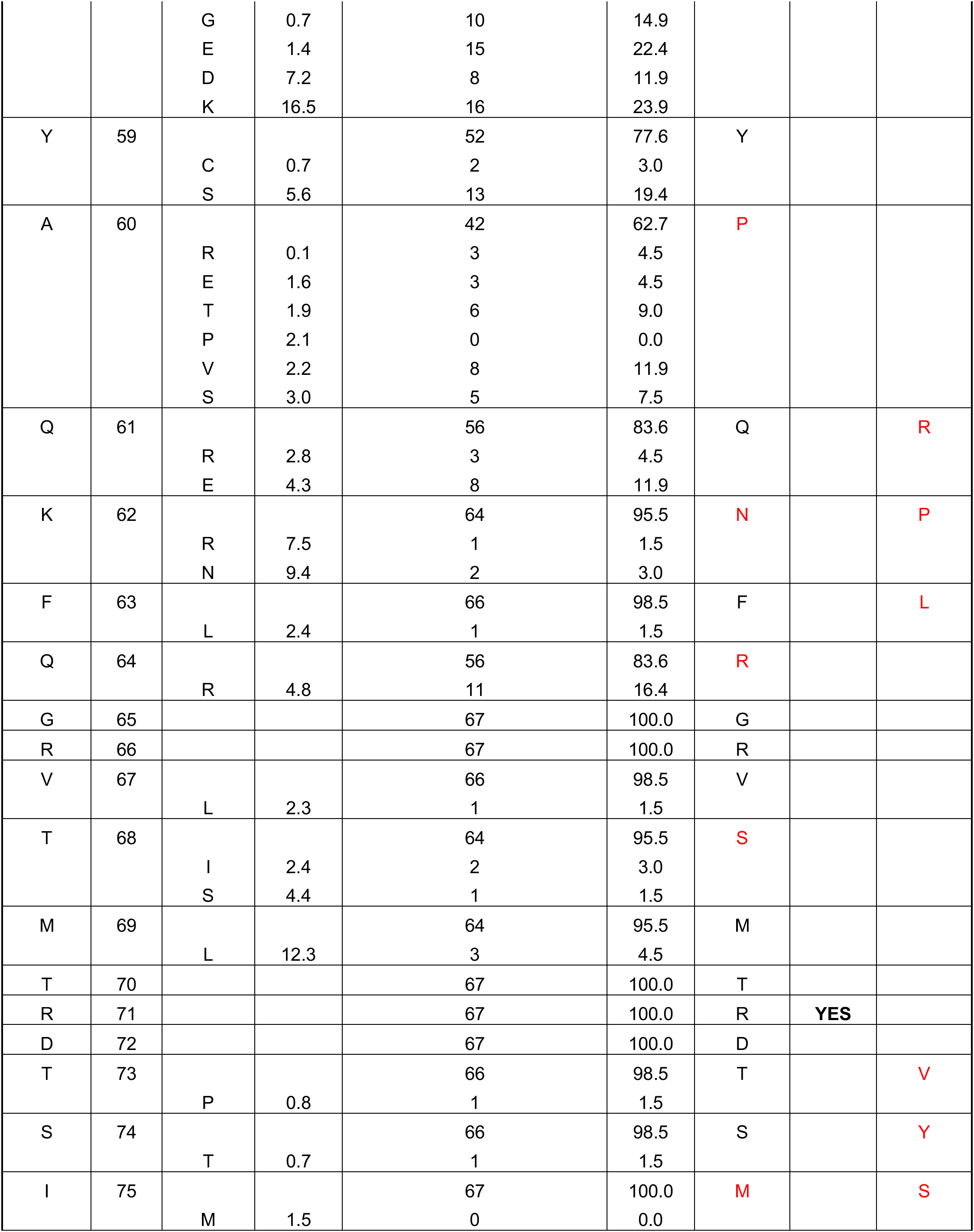
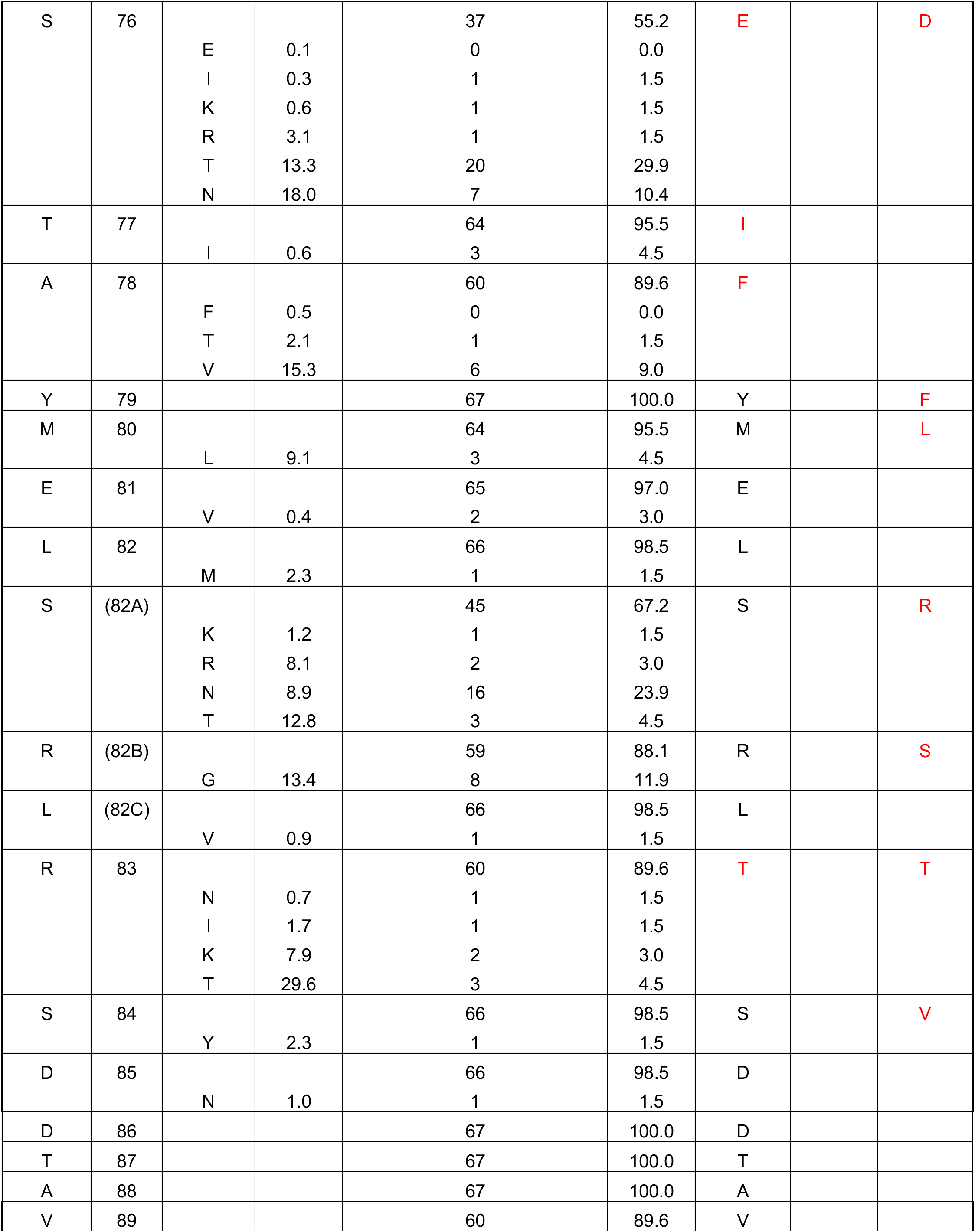
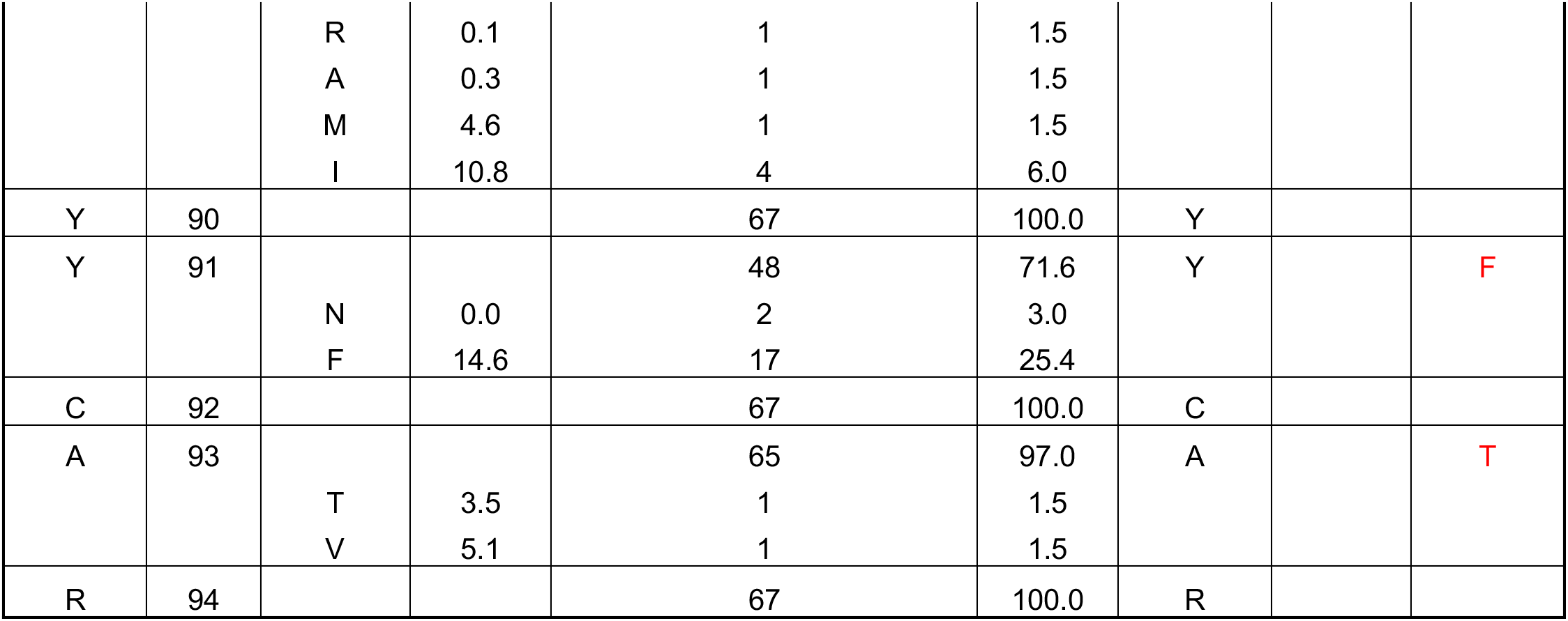
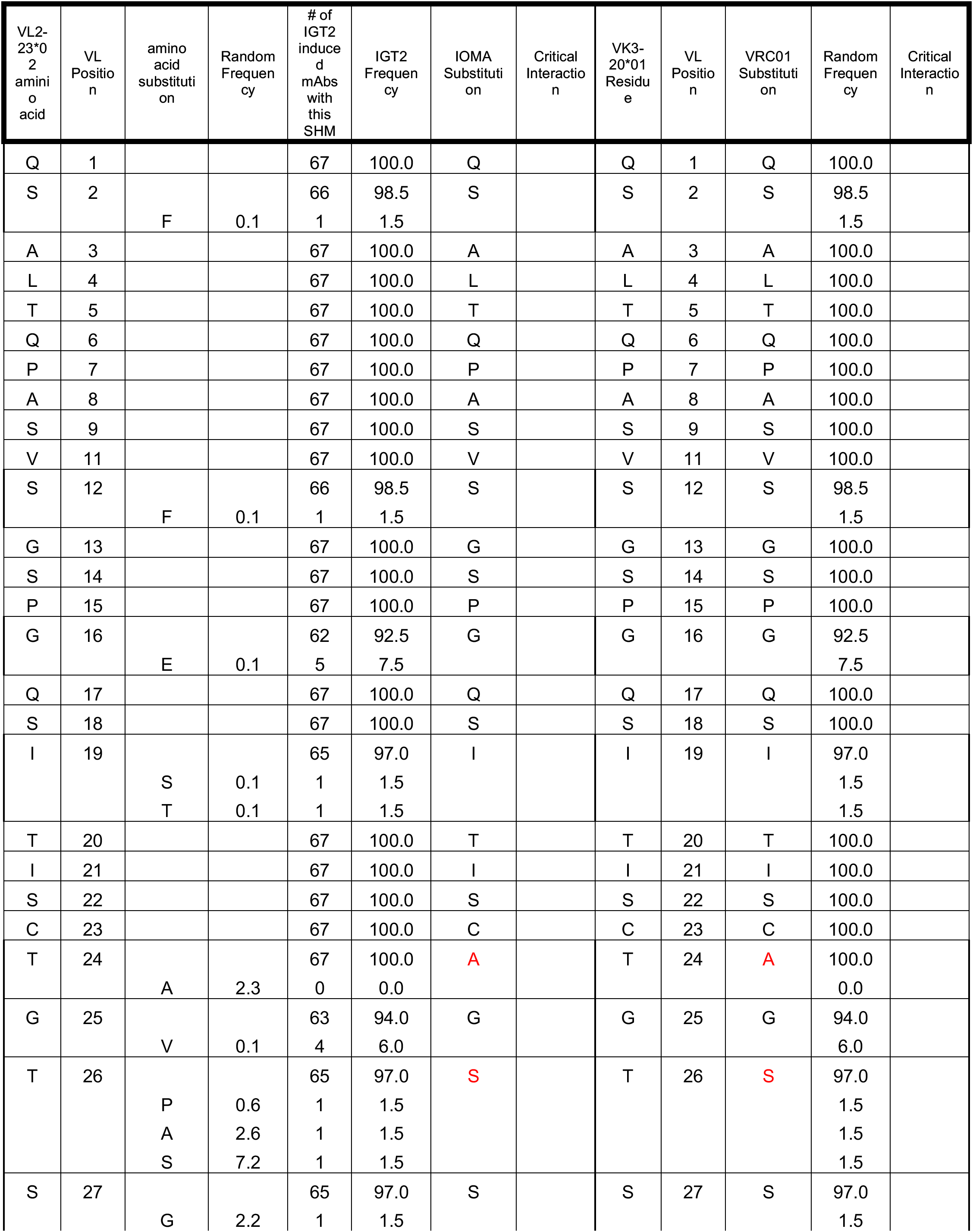
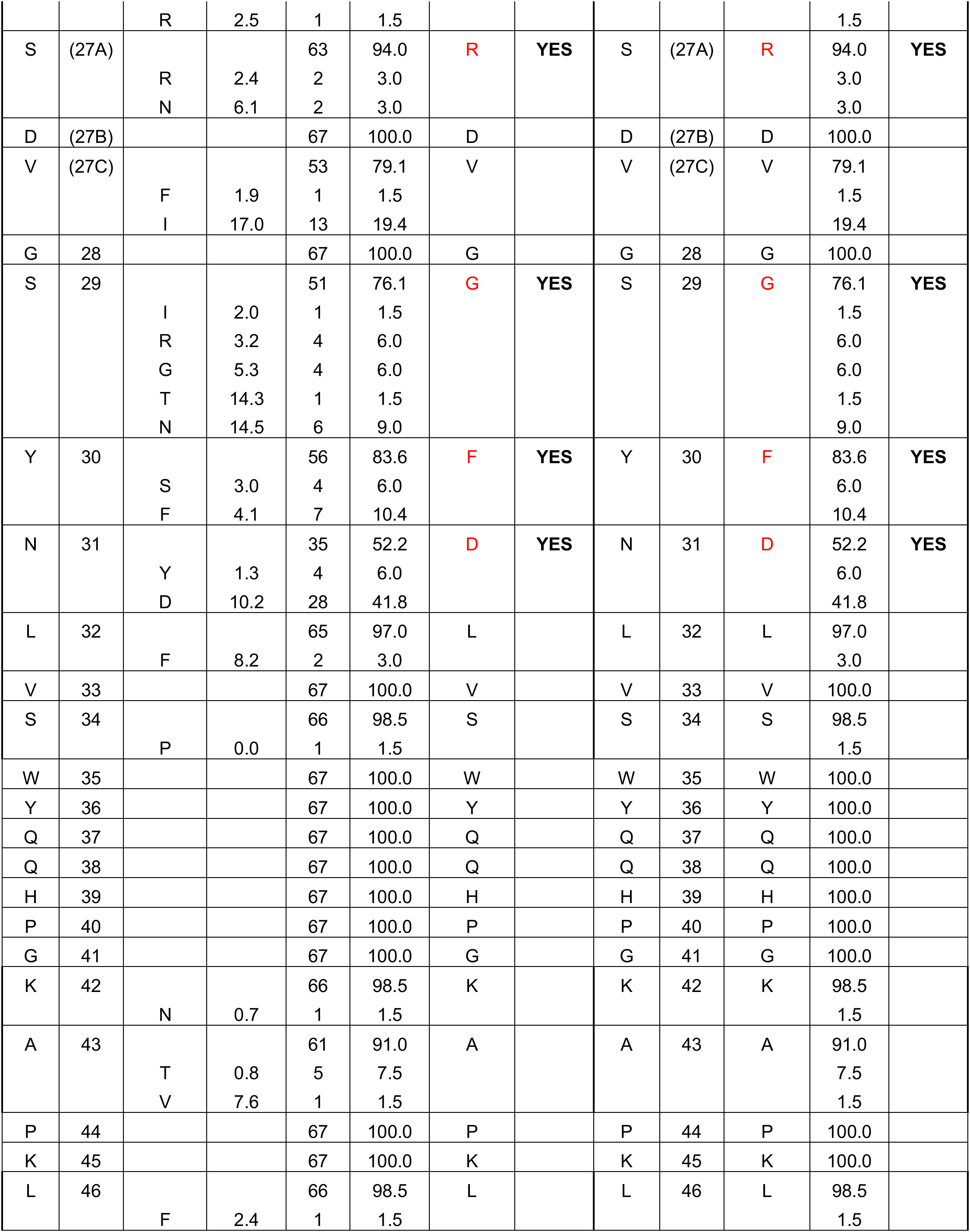
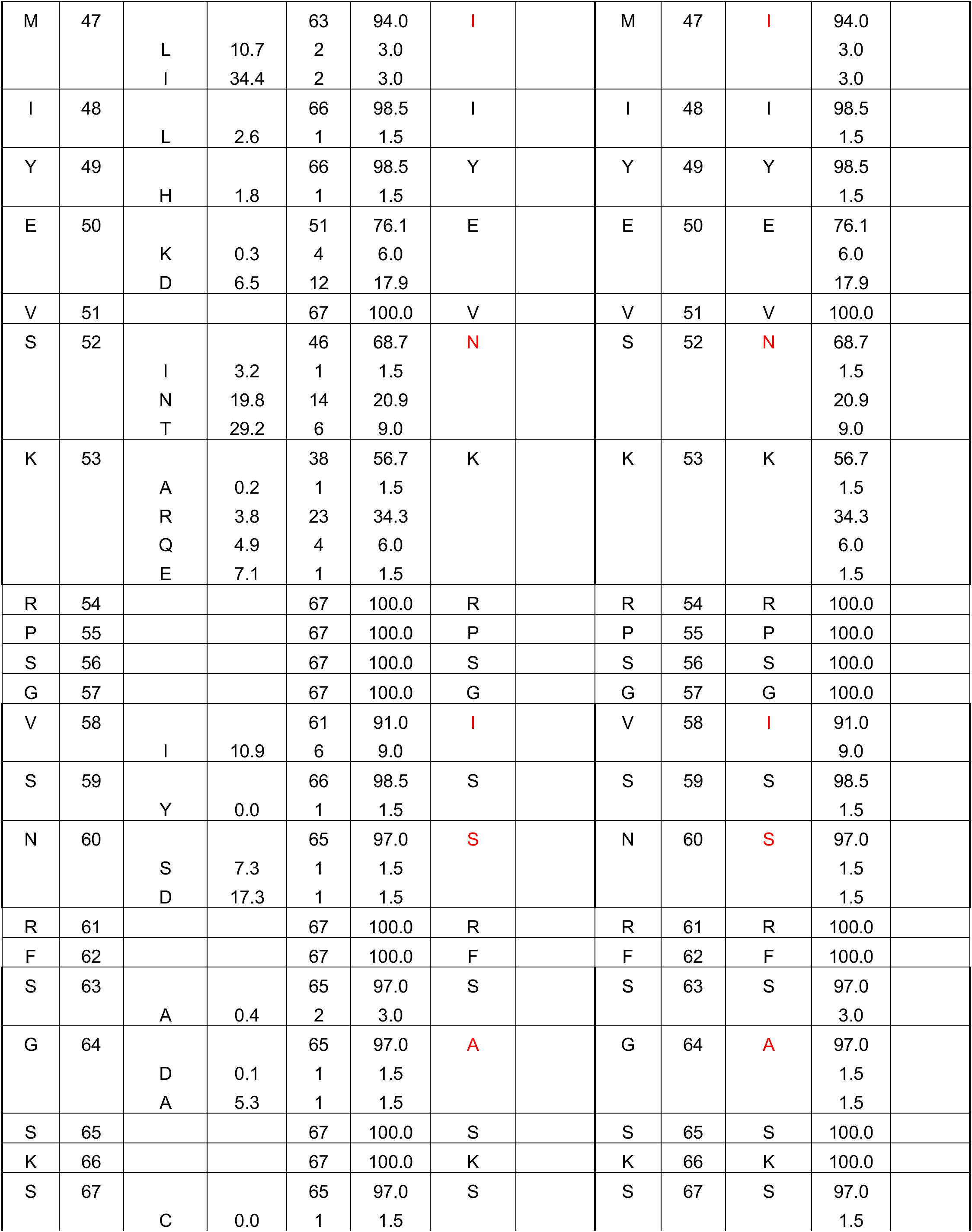
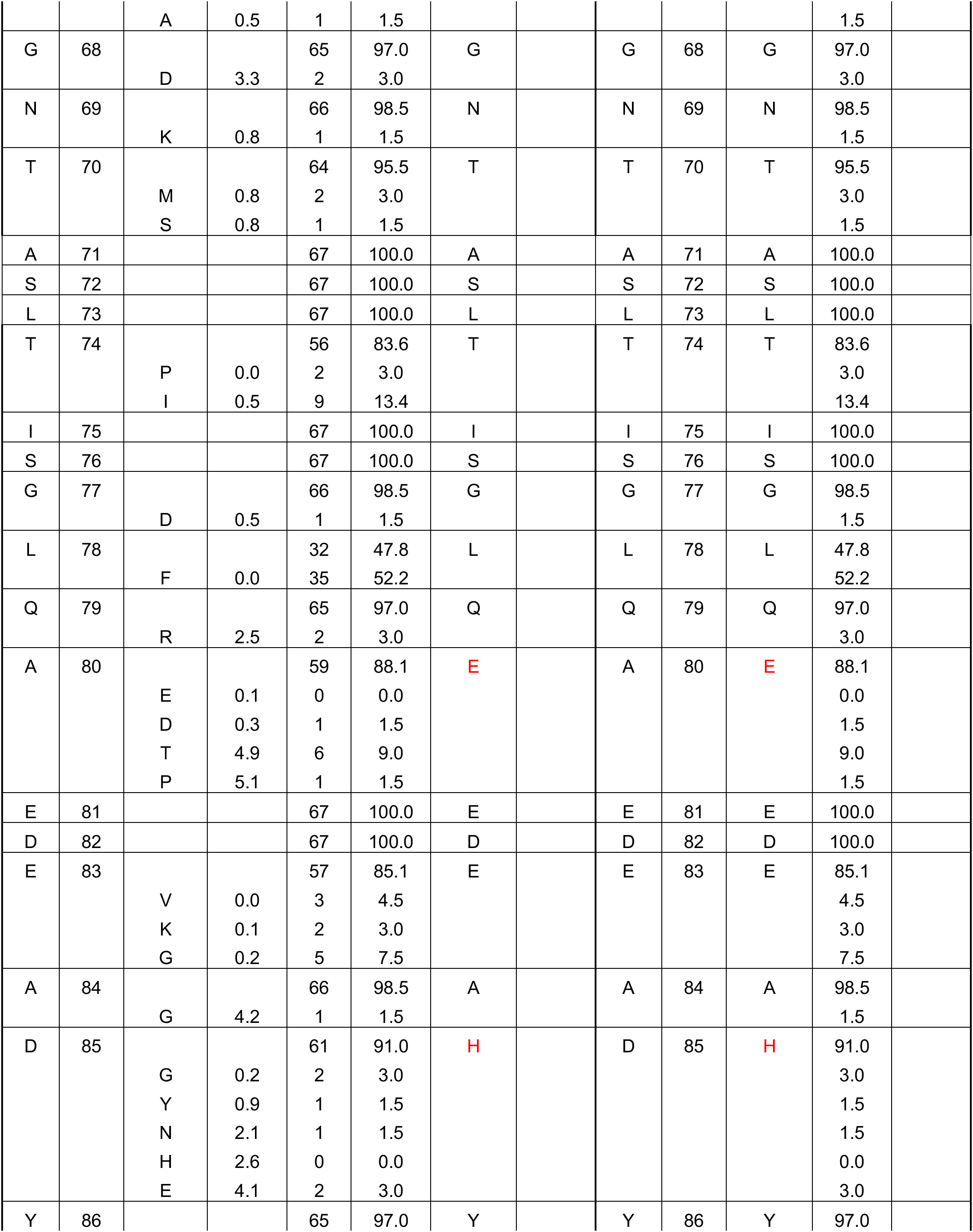
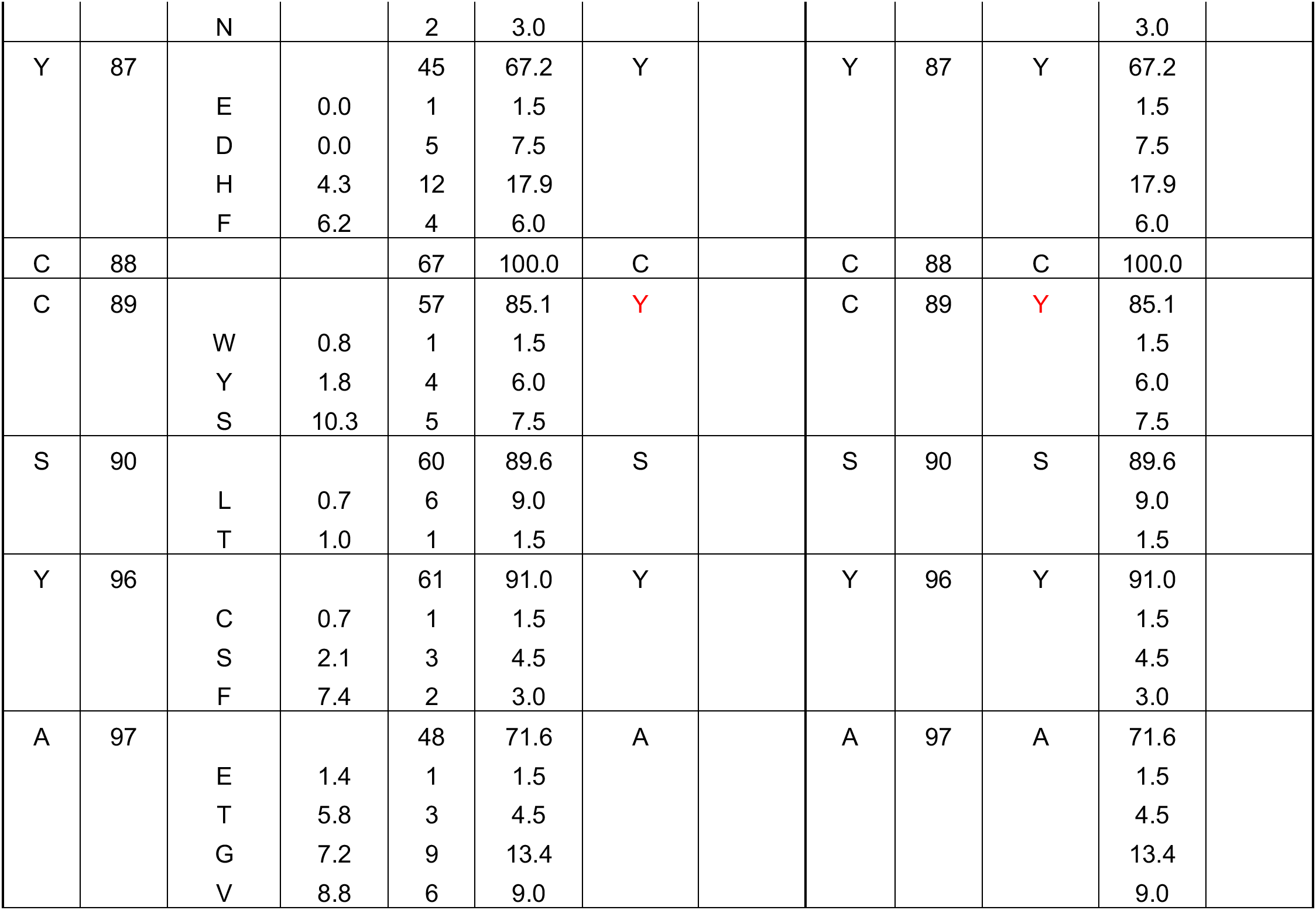
Mutational analysis of antibodies isolated from IOMA iGL transgenic mice.

**Table S5:**
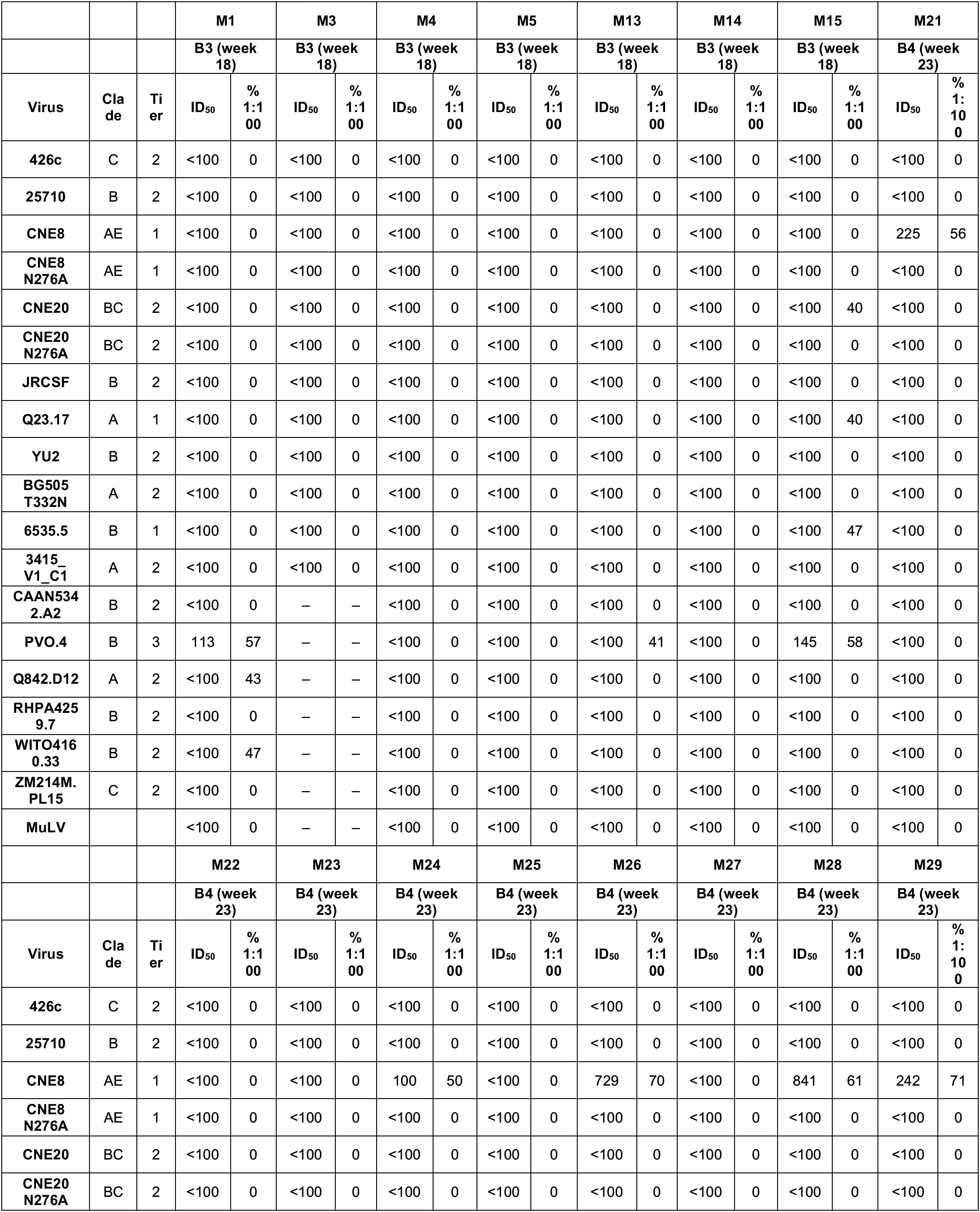
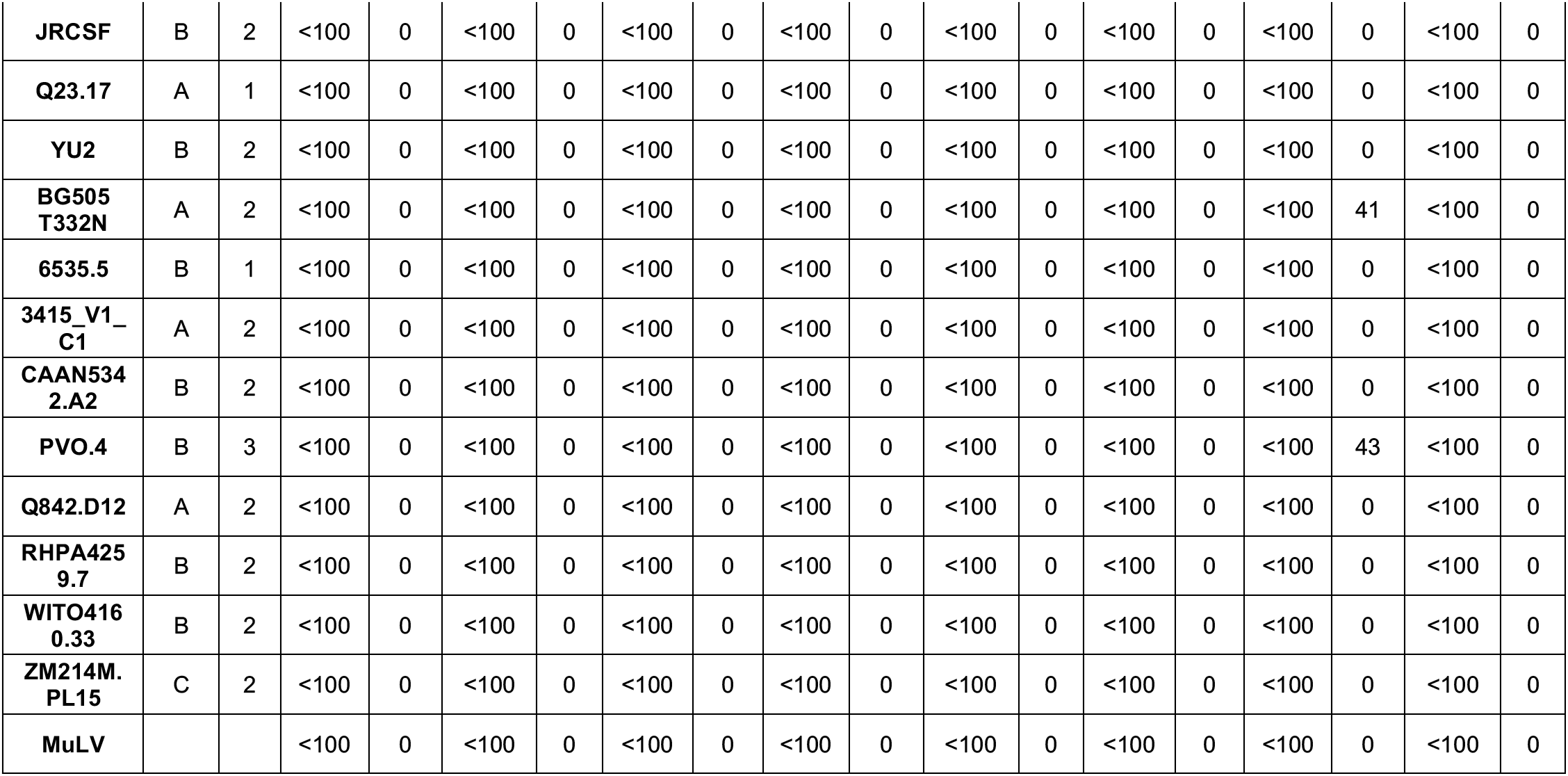
Serum neutralization in wildtype mice.

**Table S6:**
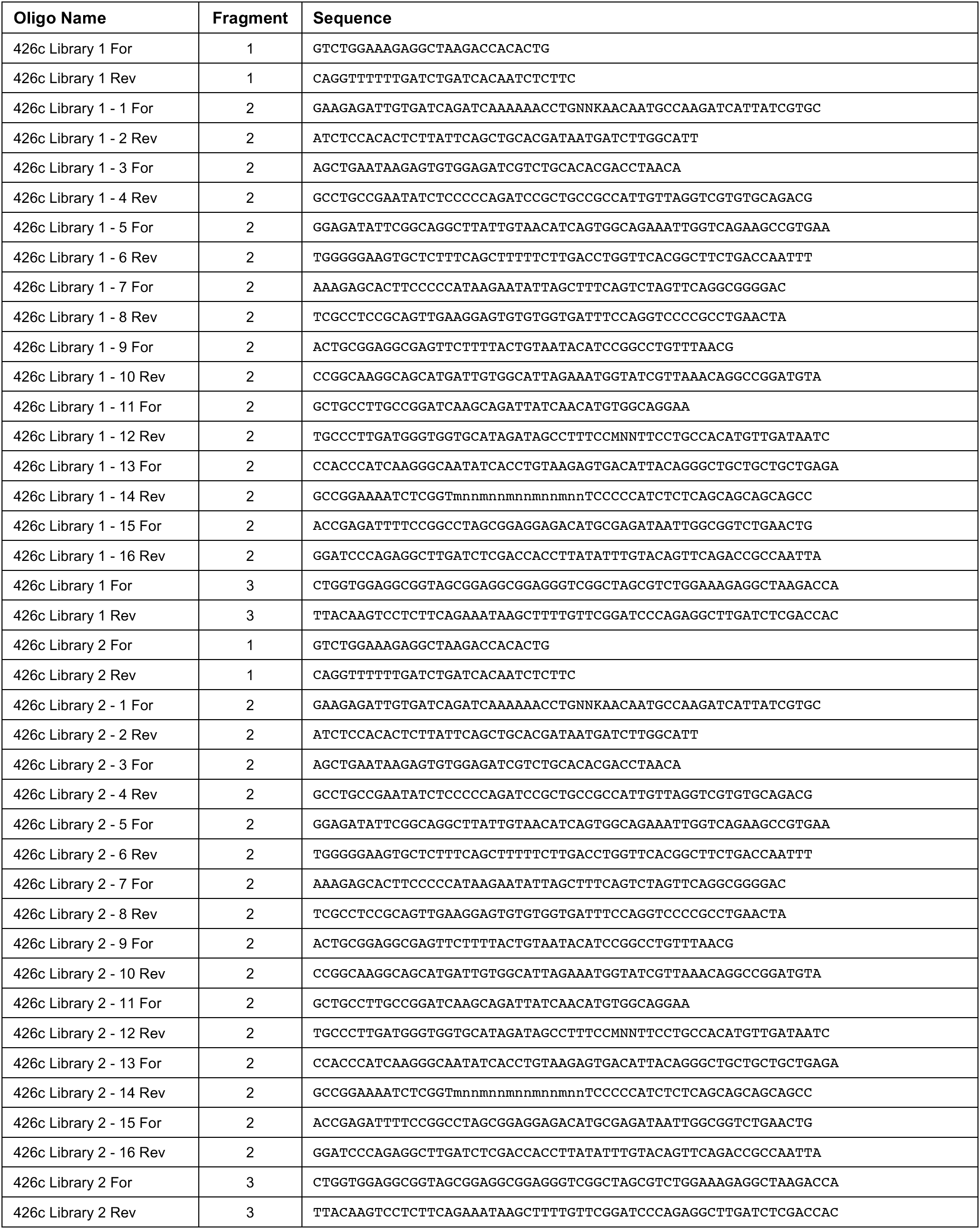
Oligonucleotides used to generate yeast display gp120 libraries.

**Table S7:**
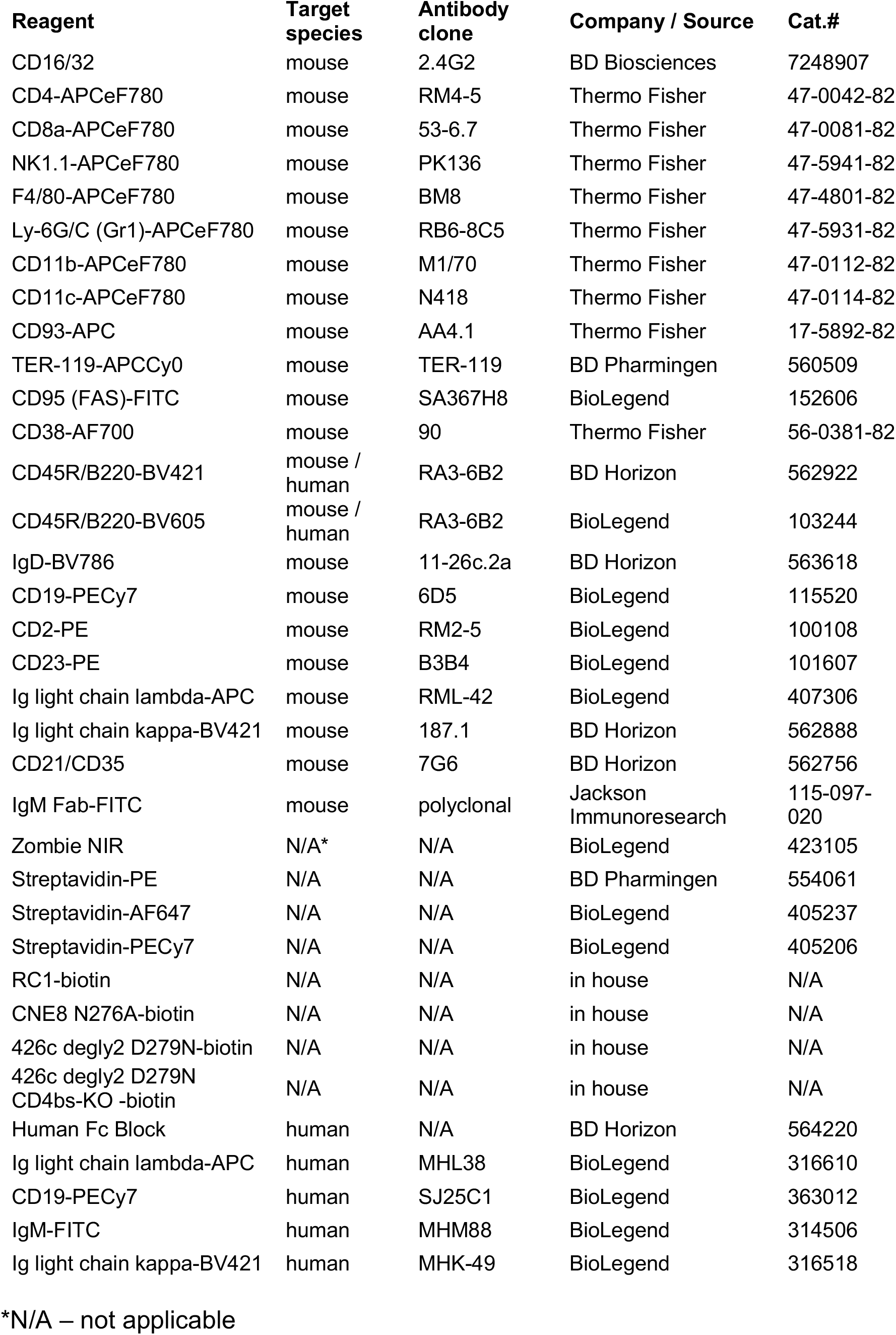
Flow cytometric reagents.

**Table S8:**
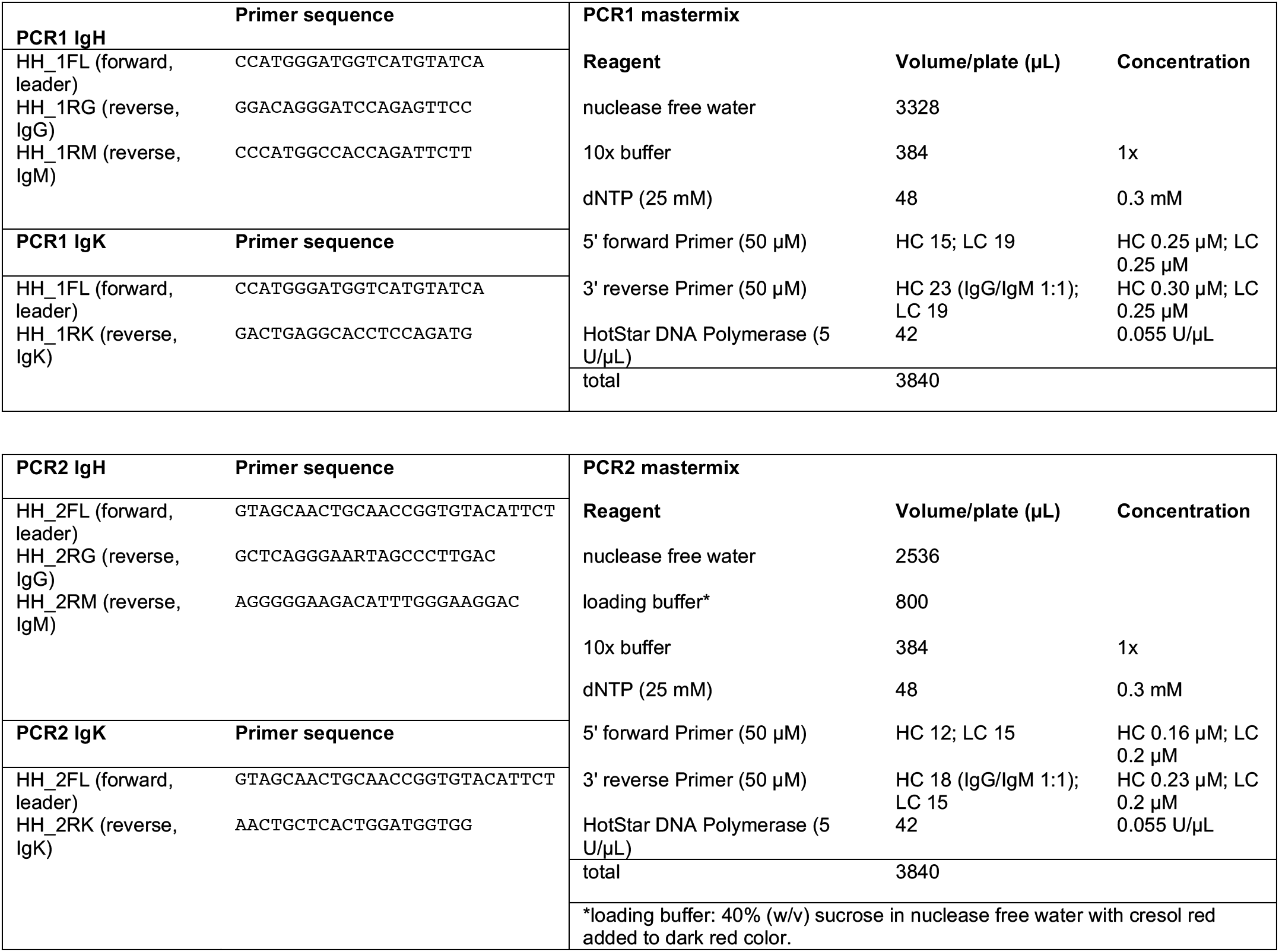
Single cell antibody cloning reaction conditions.

